# Rice bran supplementation modulates growth, microbiome and metabolome in weaning infants: a clinical trial in Nicaragua and Mali

**DOI:** 10.1101/530089

**Authors:** Luis E. Zambrana, Starin McKeen, Hend Ibrahim, Iman Zarei, Erica C. Borresen, Lassina Doumbia, Abdoulaye Bore, Alima Cissoko, Seydou Douyon, Karim Kone, Johann Perez, Claudia Perez, Ann Hess, Zaid Abdo, Lansana Sangare, Ababacar Maiga, Sylvia Becker-Dreps, Lijuan Yuan, Ousmane Koita, Samuel Vilchez, Elizabeth P. Ryan

## Abstract

Dietary rice bran supplementation during infant weaning from 6-12 months of age improved growth outcomes, modulated environmental enteric dysfunction biomarkers, and supported metabolism by the gut microbiome.

The authors declare no competing financial or non financial interests to disclose as defined by Nature Research. There are also no other interests that might be perceived to influence the results and/or discussion reported in this paper.

Correspondence and requests for materials should be addressed to Dr. Elizabeth Ryan.

**Abstract:** Rice bran supplementation provides nutrients, prebiotics and phytochemicals that enhance gut immunity, reduce enteric pathogens in mice and diarrhea in neonatal pigs, and warranted attention for improvement of environmental enteric dysfunction (EED) in children at risk. EED is a condition that drives childhood stunting via intestinal dysbiosis and impaired nutrient metabolism. This study investigated effects of rice bran supplementation on growth, EED biomarkers, gut microbiome and metabolome in weaning infants from 6 to 12 months old in Nicaragua and Mali. Healthy infants were randomized to a control group or rice bran group that received daily supplementation at increasing doses each month. Stool microbiomes were characterized using 16S rDNA amplicon sequencing. Stool metabolomes were analyzed using ultra-high-performance liquid-chromatography tandem mass-spectrometry. Statistical comparisons were completed at 6, 8, and 12 months of age. Daily consumption of rice bran was safe and feasible for infant growth, decreasing alpha-1 antitrypsin levels, and modulating gut microbiome and metabolome when compared to control. Rice bran merits investigation as a practical intervention strategy that could decrease EED prevalence and risk for children from low- and middle-income countries where rice is grown as a staple food, and bran is used as animal feed or wasted.

## Introduction

The prevalence of malnutrition in low and middle-income countries (LMIC) has negative consequences on growth of children during the first five years of life and has lifelong health consequences^1, 2^. There is an increased risk of death among children under 5 years of age due to underweight, stunting, or wasting conditions^2, 3^. Risk factors for undernutrition may include, but are not limited to: low birth weight, inadequate breastfeeding, improper complementary feeding, and recurrent infections^3, 4^ Diarrheal diseases are also some of the primary causes of undernutrition in children under five years of age^1,3,4^.

Environmental enteric dysfunction (EED) is an acquired subclinical condition of the small intestine among LMIC children^5–10^. Chronic exposure to enteric pathogens early in life is one likely contributor to EED^11^. The altered gastrointestinal functions in EED include intestinal nutrient malabsorption and increased intestinal permeability that leads to protein loss^6, 7^. Infant weaning has been identified as a critical window for intervention^12^. Previous intervention efforts in young children have targeted micronutrient deficiencies, such as Vitamin A, Zn and Fe^13–16^, oral rehydration salts for treating diarrhea^17^, antimicrobial use^18, 19^, and community hygiene improvements^20^.

Rice bran is a nutrient dense food with bioactive phytochemicals shown to prevent enteric pathogens and diarrheal disease in mice and pigs^21–24^, and favorably promotes gut health in adults^25, 26^. The effect of rice bran supplementation on host resistance to enteric infections and enhanced gut mucosal immunity was demonstrated for *Salmonella enterica* Typhimurium^23, 24^, rotavirus^27–29^, and norovirus^21^. Rice bran merits attention because it is widely available for consumption globally^30^, and particularly in LMIC regions where EED is prevalent^31^.

Stool EED biomarkers, gut microbiome^32^ and metabolome analysis^15, 33^ became important surrogate markers for analysis as intestinal tissue from infants is not easily accessible to evaluate. Stool myeloperoxidase (MPO)^34^, calprotectin (CAL)^35^, and neopterin (NEO)^36^ are indicators of inflammation; and alpha-1-antitrypsin (AAT)^34^ is an indicator of barrier lumen disruption. Chronic, elevated concentrations of all four biomarkers have been associated with poor linear growth in infants up to 24 months old^10,34–36^, and as the gut microbiome is maturing over the first 3 years of life^37, 38^. Gut microbiome composition and metabolism is influenced by delivery mode, environment, and nutrition^39, 40^. Recent studies have demonstrated that malnutrition and immature microbiomes of infants are only partially, and temporarily improved by some nutritional interventions^15, 41^. The nutritional composition and metabolic profile of rice bran, which comprises a large suite of bioactive molecules, showed benefits in animal studies that provided important rationale for investigation of dietary feasibility in weaning infants and for improving growth in LMIC children.

The major objective of this study was to investigate effects of dietary rice bran supplementation during infant weaning on growth, EED biomarkers, gut microbiome and metabolome from six to twelve months of age in Nicaragua and Mali. The findings support that daily consumption of rice bran for six months is tolerable, safe and feasible for children during weaning and was associated with improved growth and decreased gut permeability via favorable modulation of the gut microbiome and metabolome.

## Results

### Rice bran supplementation is safe and feasible for weaning infants

Daily rice bran consumption was completed in a randomized controlled trial with infants from 6 to 12 months of age. To study the effect of daily rice bran supplementation on the gut, monthly stool samples from 47 Nicaraguan and 48 Malian children were collected (average of 7 samples per child, total of 567 samples). The flow and number of infants from study recruitment to study completion is shown in **Fig. 1**. Baseline participant characteristics for Nicaragua and Mali are shown in **Table 1**. We collected information on demographic factors and infants’ household characteristics. In Nicaragua, 54.2% of infants were born via caesarean section in the control group and 30.4% in the rice bran group. All participants from Mali were delivered vaginally. For breastfeeding status, 96% of the control group and 83% in the rice bran group were consuming breast milk at six months old in Nicaragua and all children in the Mali group were consuming breast milk at the beginning and throughout the study. Of the 95 infants enrolled, 52 received antibiotics with 87 total antibiotic courses in the six-month period. Most courses consisted of systemic antibiotics given orally, with some delivered by injection for respiratory, skin, ear or diarrheal infections.

**Fig. 1.**
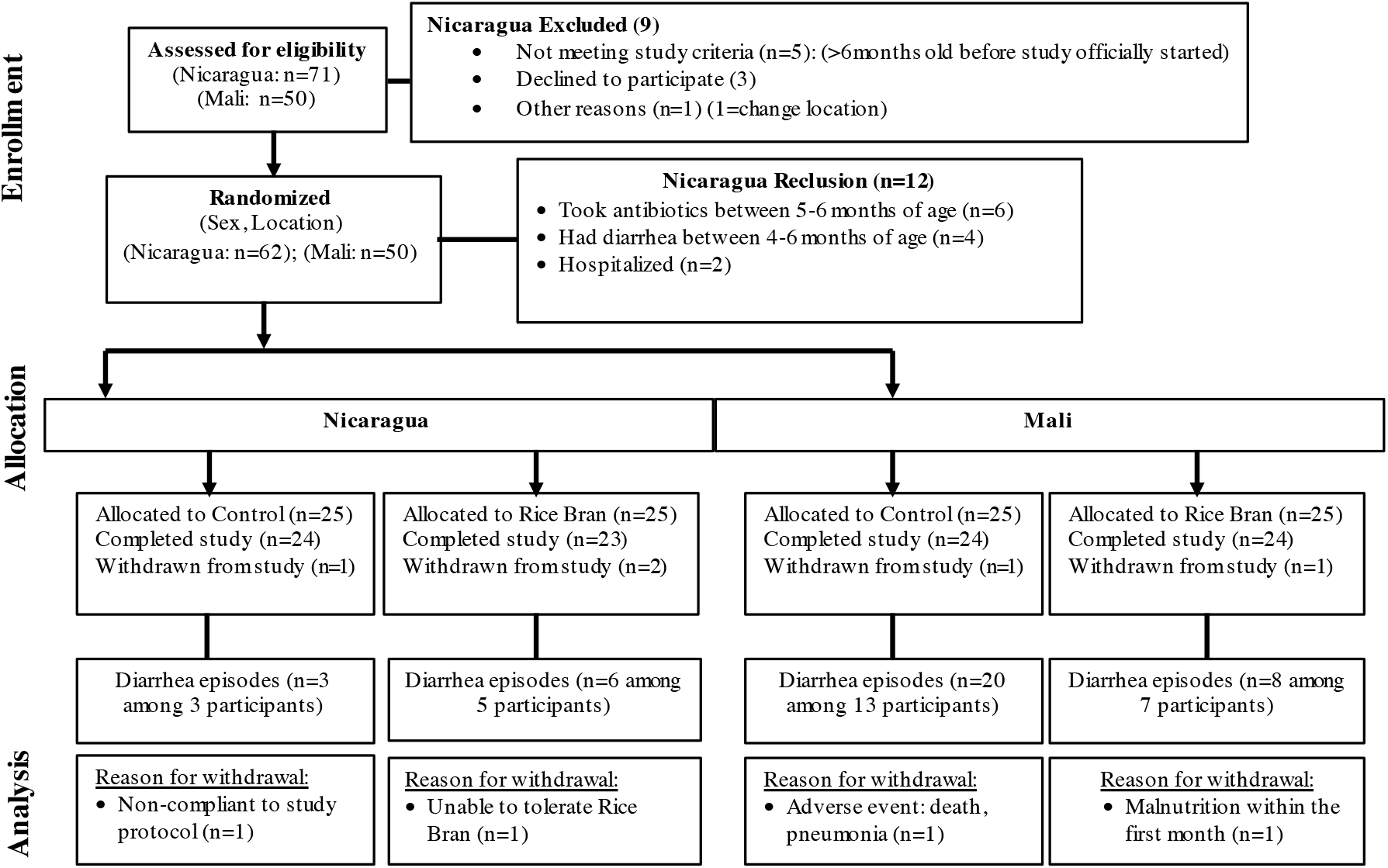
Study recruitment and participation based on CONSORT statement guidelines for clinical trials conducted in Nicaragua and Mali. (NCT02615886, NCT0255737315). 95 infants from León, Nicaragua and Dioro, Mali enrolled after meeting eligibility criteria, randomized by sex and location to one of two study arms. The number of diarrhea episodes and reasons for withdrawal were reported for each child.

**Table 1.**
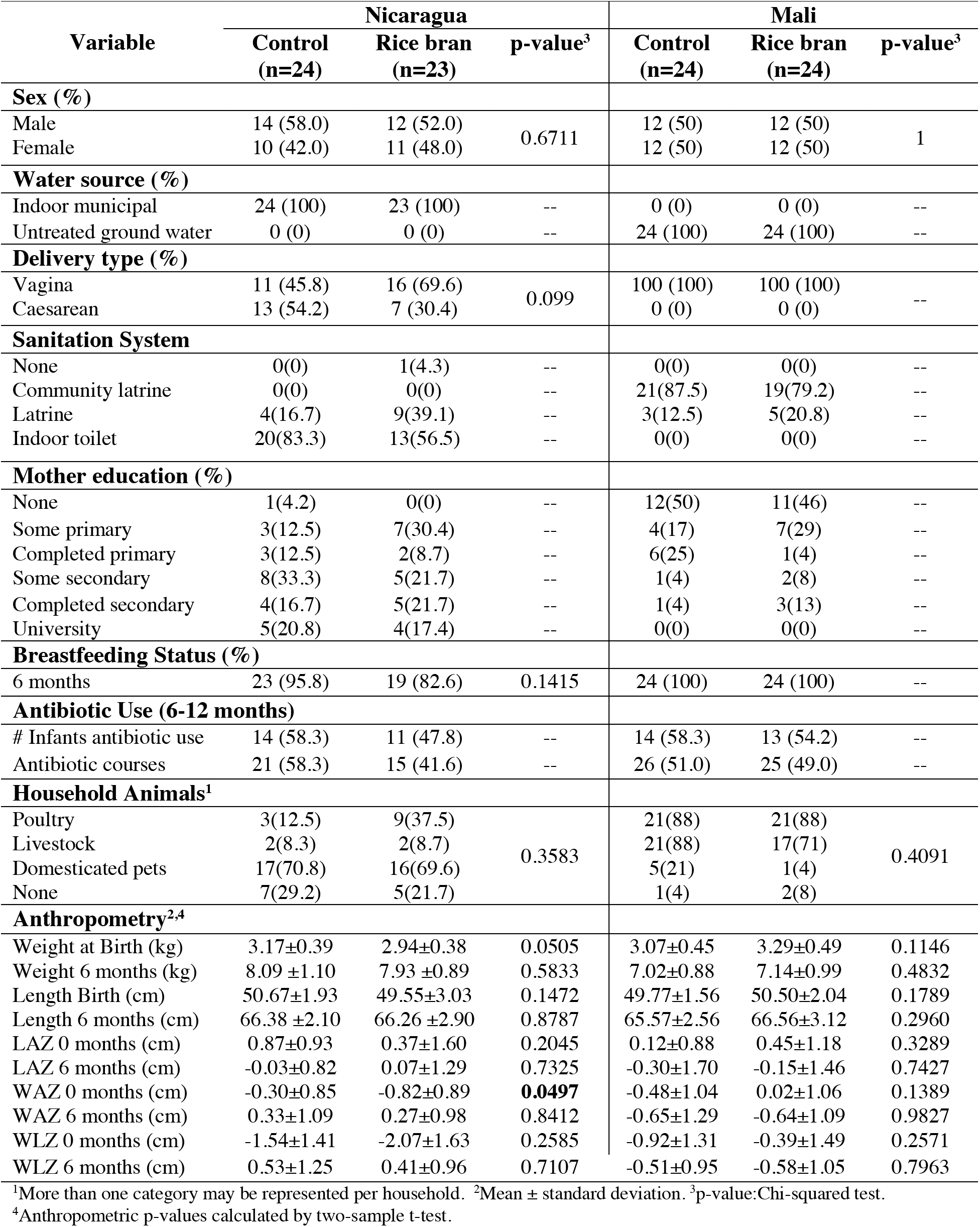
Baseline infant participant characteristics from Nicaragua and Mali.

| Variable | Nicaragua |  |  | Mali |  |  |
| --- | --- | --- | --- | --- | --- | --- |
|  | Control<br>(n=24) | Rice bran<br>(n=23) | p-value<sup>3</sup> | Control<br>(n=24) | Rice bran<br>(n=24) | p-value<sup>3</sup> |
| **Sex (%)** |  |  |  |  |  |  |
| Male | 14 (58.0) | 12 (52.0) | 0.6711 | 12 (50) | 12 (50) | 1 |
| Female | 10 (42.0) | 11 (48.0) |  | 12 (50) | 12 (50) |  |
| **Water source (%)** |  |  |  |  |  |  |
| Indoor municipal | 24 (100) | 23 (100) | -- | 0 (0) | 0 (0) | -- |
| Untreated ground water | 0 (0) | 0 (0) | -- | 24 (100) | 24 (100) | -- |
| **Delivery type (%)** |  |  |  |  |  |  |
| Vagina | 11 (45.8) | 16 (69.6) | 0.099 | 100 (100) | 100 (100) | -- |
| Caesarean | 13 (54.2) | 7 (30.4) |  | 0 (0) | 0 (0) |  |
| **Sanitation System** |  |  |  |  |  |  |
| None | 0(0) | 1(4.3) | -- | 0(0) | 0(0) | -- |
| Community latrine | 0(0) | 0(0) | -- | 21(87.5) | 19(79.2) | -- |
| Latrine | 4(16.7) | 9(39.1) | -- | 3(12.5) | 5(20.8) | -- |
| Indoor toilet | 20(83.3) | 13(56.5) | -- | 0(0) | 0(0) | -- |
| **Mother education (%)** |  |  |  |  |  |  |
| None | 1(4.2) | 0(0) | -- | 12(50) | 11(46) | -- |
| Some primary | 3(12.5) | 7(30.4) | -- | 4(17) | 7(29) | -- |
| Completed primary | 3(12.5) | 2(8.7) | -- | 6(25) | 1(4) | -- |
| Some secondary | 8(33.3) | 5(21.7) | -- | 1(4) | 2(8) | -- |
| Completed secondary | 4(16.7) | 5(21.7) | -- | 1(4) | 3(13) | -- |
| University | 5(20.8) | 4(17.4) | -- | 0(0) | 0(0) | -- |
| **Breastfeeding Status (%)** |  |  |  |  |  |  |
| 6 months | 23 (95.8) | 19 (82.6) | 0.1415 | 24 (100) | 24 (100) | -- |
| **Antibiotic Use (6-12 months)** |  |  |  |  |  |  |
| # Infants antibiotic use | 14 (58.3) | 11 (47.8) | -- | 14 (58.3) | 13 (54.2) | -- |
| Antibiotic courses | 21 (58.3) | 15 (41.6) | -- | 26 (51.0) | 25 (49.0) | -- |
| **Household Animals<sup>1</sup>** |  |  |  |  |  |  |
| Poultry | 3(12.5) | 9(37.5) | 0.3583 | 21(88) | 21(88) | 0.4091 |
| Livestock | 2(8.3) | 2(8.7) |  | 21(88) | 17(71) |  |
| Domesticated pets | 17(70.8) | 16(69.6) |  | 5(21) | 1(4) |  |
| None | 7(29.2) | 5(21.7) |  | 1(4) | 2(8) |  |
| **Anthropometry<sup>2,4</sup>** |  |  |  |  |  |  |
| Weight at Birth (kg) | 3.17±0.39 | 2.94±0.38 | 0.0505 | 3.07±0.45 | 3.29±0.49 | 0.1146 |
| Weight 6 months (kg) | 8.09 ±1.10 | 7.93 ±0.89 | 0.5833 | 7.02±0.88 | 7.14±0.99 | 0.4832 |
| Length Birth (cm) | 50.67±1.93 | 49.55±3.03 | 0.1472 | 49.77±1.56 | 50.50±2.04 | 0.1789 |
| Length 6 months (cm) | 66.38 ±2.10 | 66.26 ±2.90 | 0.8787 | 65.57±2.56 | 66.56±3.12 | 0.2960 |
| LAZ 0 months (cm) | 0.87±0.93 | 0.37±1.60 | 0.2045 | 0.12±0.88 | 0.45±1.18 | 0.3289 |
| LAZ 6 months (cm) | -0.03±0.82 | 0.07±1.29 | 0.7325 | -0.30±1.70 | -0.15±1.46 | 0.7427 |
| WAZ 0 months (cm) | -0.30±0.85 | -0.82±0.89 | **0.0497** | -0.48±1.04 | 0.02±1.06 | 0.1389 |
| WAZ 6 months (cm) | 0.33±1.09 | 0.27±0.98 | 0.8412 | -0.65±1.29 | -0.64±1.09 | 0.9827 |
| WLZ 0 months (cm) | -1.54±1.41 | -2.07±1.63 | 0.2585 | -0.92±1.31 | -0.39±1.49 | 0.2571 |
| WLZ 6 months (cm) | 0.53±1.25 | 0.41±0.96 | 0.7107 | -0.51±0.95 | -0.58±1.05 | 0.7963 |

<sup>1</sup>More than one category may be represented per household. <sup>2</sup>Mean ± standard deviation. <sup>3</sup>p-value:Chi-squared test.
<sup>4</sup>Anthropometric p-values calculated by two-sample t-test.

Dietary compliance to rice bran was averaged per month during the 6-month intervention with no adverse events related to rice bran intake at the increasing doses over time. Compliance to the rice bran intervention in Nicaragua was 90% and in Mali was > 99%. The feasibility and tolerability for infants to consume rice bran at increasing doses (1-5g/day) over the six month study period was demonstrated when mothers fed rice bran powder alone or reported consumption with drinking water, staple grain porridges (i.e. millet, sorghum, and white rice), soups, milk, fruits, juices, eggs, and fish when available. Rice bran supplementation increases infant growth

Anthropometric data was collected using standardized procedures across study sites at 6, 8, and 12 months of age for each child and included length-for-age Z-score (LAZ), weight-for-age Z-score (WAZ) and weight-for-length Z-score (WLZ). **Table 2** reports the z-scores analyzed by repeated measures and adjusted by treatment (rice bran) and age (6-8 months and 8-12 months) by country. Significant differences were observed for anthropometric measures between treatment, and ages in both cohorts. In Nicaraguan infants consuming rice bran, LAZ was significant over time (1.18 at 6-8 months, p-value= 0.0000; 0.35 at 8-12 months, p-value= 0.0002) and for WLZ at 6-8 months (p-value= 0.0000), with no changes detected in WAZ. Malian infants consuming rice bran had significant growth results for WAZ (6-8 months, p-value= 0.0001, 8-12 months, p-value= 0.0175) and WLZ (6-8 months, p-value= 0.0141, 8-12 months, p-value= 0.0134). **Fig. 2** displays LAZ, WAZ and WLZ over time and by country. The significant increase in LAZ at 8 months and 12 months in Nicaraguan infants that consumed rice bran was compared to control group (**Fig. 2A**, p<0.01). No significant differences were observed for WAZ and WLZ with this control group comparison (**Fig. 2B** and **2C**).

**Fig. 2.**
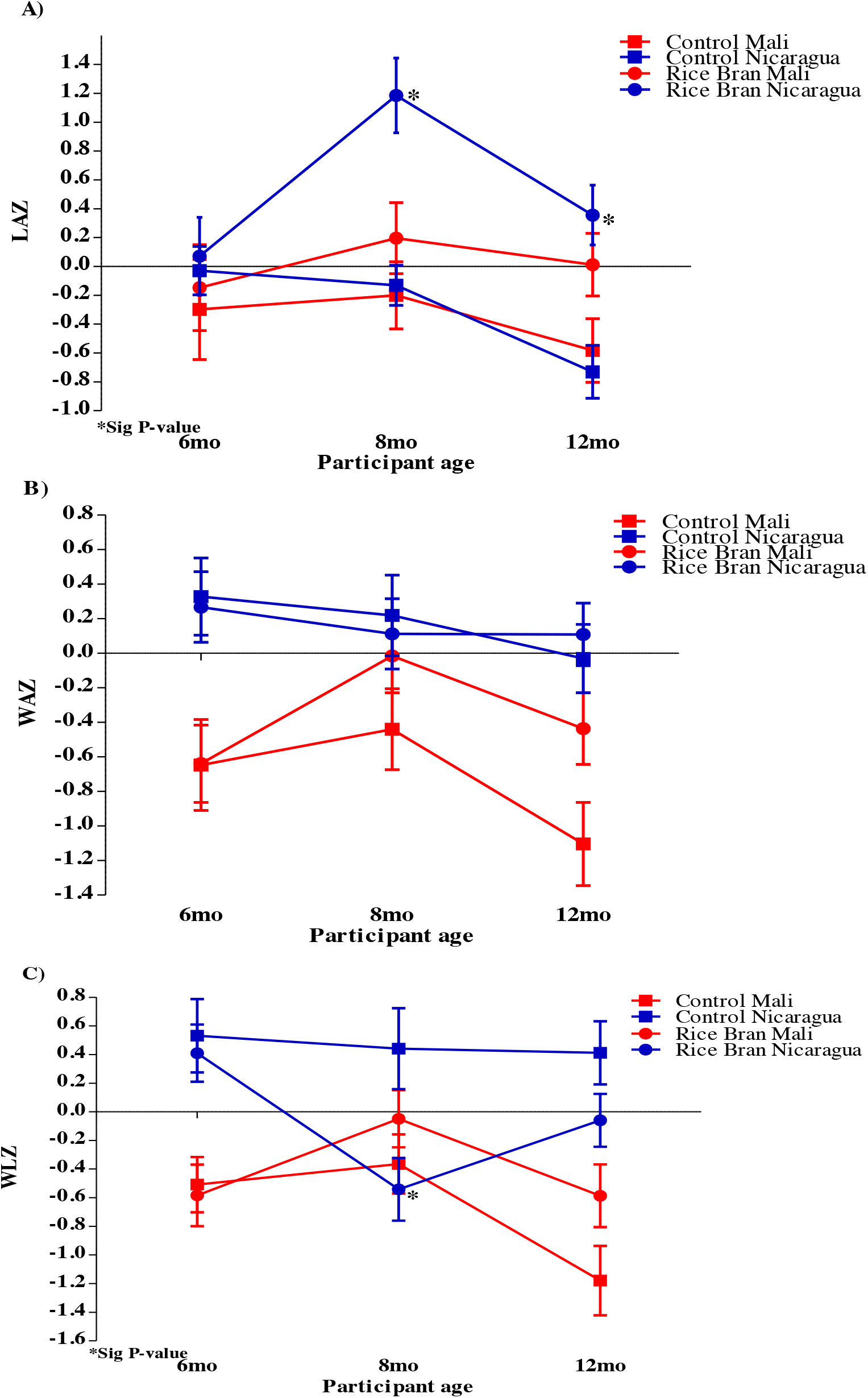
Anthropometric Z-scores for Nicaraguan and Malian infants in rice bran and control groups at 6, 8 and 12 months. A). Significant LAZ (p<0.05) at 8 and 12 months in the rice bran group compared to control for Nicaraguan infants. B). No WAZ significant changes between rice bran and control group in Nicaraguan and Malian infants. C). WLZ at 8 months was significantly lower for the rice bran group compared to control in Nicaragua.

**Table 2.**
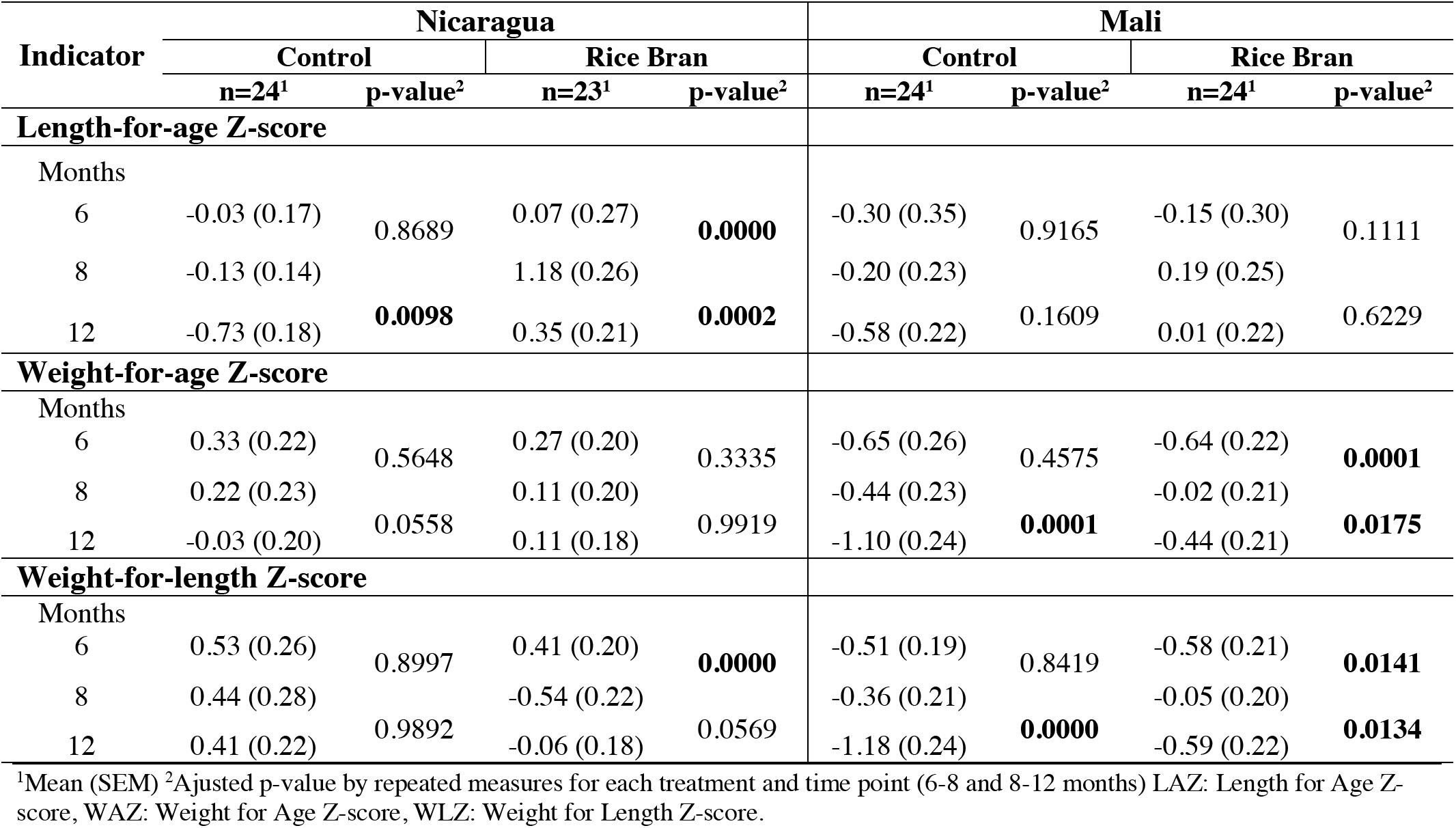
Anthropometric measures adjusted by treatment and age in Nicaraguan and Malian Infants

### Rice bran modulation of environmental enteric dysfunction biomarkers

Four EED stool biomarkers were selected for analysis at 6, 8 and 12 months of age using ELISA (see materials and methods). A significant decrease in AAT was observed at 12 months of age (p=0.0368) in Nicaragua infants that consumed rice bran compared to control (**Table 3**). No significant differences were detected in NEO, MPO and CAL between treatment groups, however a slight decrease in median concentrations of all stool EED biomarkers were observed in the rice bran group compared to control in both countries that can be applied for future study sample size and power calculations.

**Table 3.** Environmental enteric dysfunction (EED) biomarkers in stool at 6, 8, and 12 months of age for Nicaraguan and Malian infants.

| EED Biomarker | Nicaragua |  |  | Mali |  |  |
| --- | --- | --- | --- | --- | --- | --- |
|  | Control<sup>1</sup><br>(n=24) | Rice Bran<sup>1</sup><br>(n=23) | p-value<sup>2</sup> | Control<sup>1</sup><br>(n=24) | Rice Bran<sup>1</sup><br>(n=24) | p-value<sup>2</sup> |
| **Neopterin (nmol/L)** |  |  |  |  |  |  |
| 6 | 150.8 (182.2) | 208.6 (131.7) | 0.7220 | 20.2 (31.0) | 34.6 (28.0) | 0.5929 |
| 8 | 222.3 (241.1) | 144.8 (196.9) | 0.5771 | 36.2 (27.6) | 28.9 (28.5) | 0.4314 |
| 12 | 137.5 (285.2) | 182.4 (230.4) | 0.8727 | 12.0 (33.0) | 36.0 (32.6) | 0.9880 |
| **Myeloperoxidase (ng/ml)** |  |  |  |  |  |  |
| 6 | 277.0 (374.5) | 237.5 (376.5) | 0.0847 | 3970.6 (17776.6) | 5400.9 (20794.8) | 0.6139 |
| 8 | 331.1 (312.3) | 266.3 (236.4) | 0.8454 | 15838.7 (20511.9) | 7451.0 (12972.7) | 0.3763 |
| 12 | 182.0 (324.8) | 158.5 (376.7) | 0.3345 | 4846.4 (11266.9) | 3153.9 (14095.2) | 0.5437 |
| **Calprotectin (µg/g)** |  |  |  |  |  |  |
| 6 | 32.2 (108.7) | 88.0 (281.1) | 0.2394 | 51.7 (101.2) | 53.3 (106.4) | 0.7136 |
| 8 | 24.7 (87.5) | 58.2 (213.2) | 0.8023 | 130.0 (769.7) | 68.7 (126.8) | 0.0639 |
| 12 | 24.0 (74.4) | 50.5 (133.9) | 0.2629 | 35.0 (70.2) | 20.9 (56.0) | 0.2103 |
| **Alpha-1 Antitrypsin (ng/ml)** |  |  |  |  |  |  |
| 6 | 130.1 (177.7) | 109.5 (217.4) | 0.7199 | 247.5 (499.6) | 463.1 (891.8) | 0.0999 |
| 8 | 152.0 (115.4) | 73.5 (122.4) | 0.1221 | 619.2 (759.0) | 579.9 (899.9) | 0.2237 |
| 12 | 130.9 (129.8) | 70.8 (87.8) | **0.0368** | 663.7 (580.5) | 453.2 (807.5) | 0.4796 |

<sup>1</sup>Median (IQR)
<sup>2</sup>p-value by repeated measures comparing treatments at each time point.

### Rice bran modulates gut microbial communities in Nicaraguan and Malian infants

The microbiome was characterized and compared for 48 Malian and 47 Nicaraguan infants at 8 and 12 months of age in the rice bran and control study groups. DNA was isolated from stool samples and the V4 hypervariable region of the 16S rRNA gene was sequenced utilizing the Earth Microbiome Project protocol^42–46^. Sequences were preprocessed for quality assurance and classified into operational taxonomic units (OTUs) (see materials and methods), and the results were integrated to construct family and genus-level composition profiles for all 192 samples from both countries. No major differences were detected in alpha diversity indices (Observed, Shannon, InvSimpson and Richness) calculated for rice bran group and control group at 8 and 12 months (**Table S1**). Beta diversity analysis, depicted in the Nonmetric Multidimensional Scaling (NMDS) plot based on the Bray-Curtis distance measure, indicated complete country-level separation in the overall gut microbial community composition (**Fig. 3A**). This provided rationale for separating analyses for the microbiome and metabolite profiles by country.

**Fig. 3.**
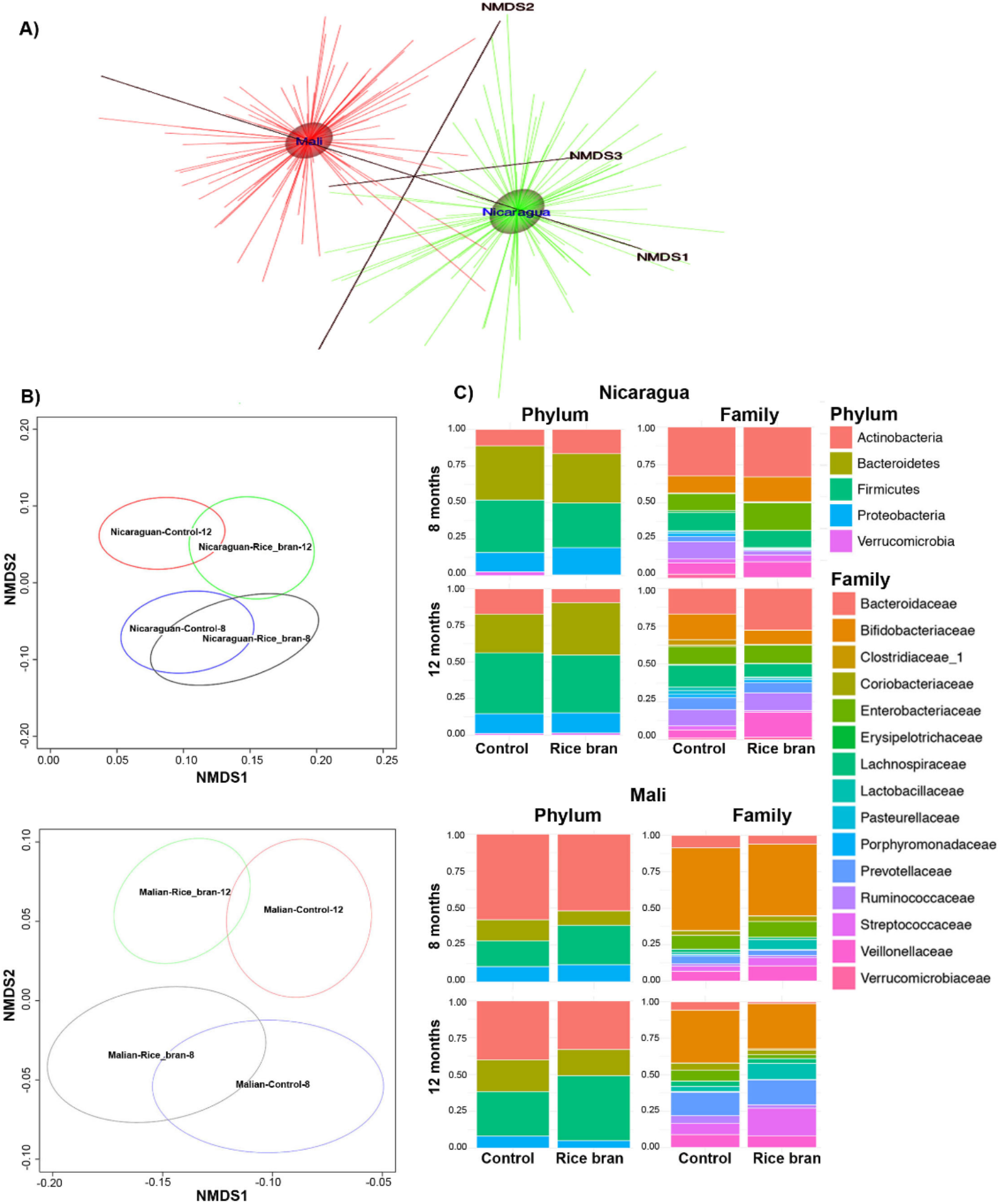
Rice bran and control infant stool microbiome at 8 and 12 months of age in Nicaragua and Mali. Nonmetric Multidimensional Scaling (NMDS) for A). Nicaragua and Mali all samples B). Control groups and rice bran groups at 8 and 12 months. NMDS was used on the OTU level to assess possible trends and clustering in the microbial community structure per treatment and time point. C). Bacterial taxa at phylum and family level in Nicaragua (top) and Mali (bottom). Bar-graphs show phylum and family relative abundance based on the resulting OTU table generated using the ggplot2 package in R. These plots were generated for the data at the phylum and the family levels and meant to describe the microbial community structure per sampled group and per time point (8 months and 12 months) under each of the treatment levels (control and rice bran).

**Fig. 3B** shows the NMDS plot separated by country. This figure highlights differences in the microbiome between two time periods, 8 and 12 months, which is more pronounced in the Malian samples. These figures show overlap of microbiomes within each time period indicating putative similarity between the microbial community structures during growth periods. This overlap was observed to a greater level at 8 months of age as compared to 12 months that may illustrate microbial adaptation to new exposures^31^.

**Fig. 4** illustrates the taxa (on the OTU level) with at least 2 log-fold change between the rice bran and control groups per country and per age group. These taxa were ordered based on significance, measured by the FDR-adjusted p-value, from bottom (most significant) to top (details are provided in **Table S2** for Nicaragua and **Table S3** for Mali). Given the measurable differences in growth between groups by 8 months of age, the effect of daily dietary supplementation of rice bran on the infant microbiome was compared to the control group at 8 months.

**Fig. 4.**
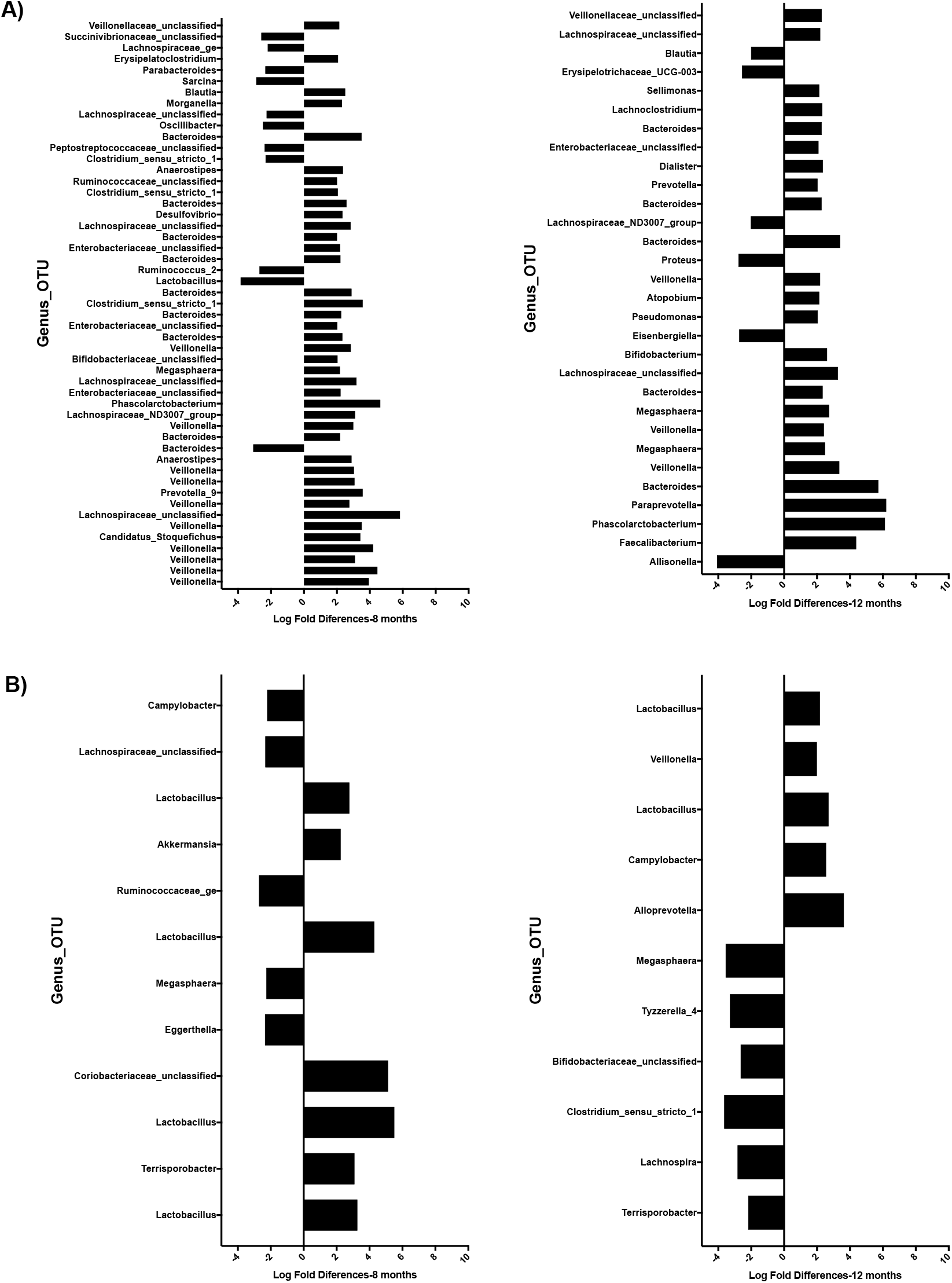
Microbiome differences between Nicaragua and Mali at 8 and 12 months between rice bran and control groups. Fold differences in relative percentage of OTUs different between control and rice bran groups at 8 months and 12 months. A) Nicaragua, and B) Mali. OTUs with fold difference more than 2 are shown for infants at 8 months (left) and 12 months (right). Fold difference for OTUs with FDR less than 0.05 is shown with the most significant OTUs on the bottom of each graph.

In Nicaragua, we identified 145 OTUs that were significantly different between control and rice bran across the samples (**Table S2**), 74 of which showed greater than or equal to 2 log fold differences between rice bran and control groups at 8 or 12 months (**Fig. 4A**, adjusted p-value <0.05). Seven of these 74 OTUs overlapped between the ages 8 and 12 months. In Mali, 42 bacterial OTUs were identified to significantly differ between control and rice bran across samples, and 19 showed more than 2 log fold changes between rice bran and control at 8 or 12 months (**Fig. 4B**, adjusted p-value < 0.05) with only three overlapping between the two age groups. Next we explored the country specific changes in genus level taxa that were responsive to rice bran intake. For Nicaraguan infants (**Table S2**), the notable rice bran responsive taxa which increased at 8 months of age compared to control group were Lachnospiraceae-unclassified-Otu0280 (log-FC 5.84, adjusted p-value 3.95E-08) Bifidobacterium-unclassified-Otu0314 (log-FC 2.04, adjusted p-value 1.38E-6), Ruminococcaceae-unclassified-Otu0238 (log-FC 2.01, adjusted p-value 0.00097), Veillonella (11different OTUs, log-FC > 2.0, adjusted p-value < 0.05), and Bacteroides (log-FC > 2.0, adjusted p-value < 0.05). The fold difference for genus level taxa that were lower in relative percent abundance for rice bran group were Bacteroides-Otu0192 (log-FC-3.08, adjusted p-value 2.34E-07), Parabacteroides-0tu0086 (log-FC −2.34, adjusted p-value 0.0074), Lachnospiraceae-unclassified-0tu0174 (log-FC −2.27, adjusted p-value 0.0033), Lactobacillus-0ut0053 (log-FC −3.85, adjusted p-value 1.09E-05), Oscillibacter (log-FC −2.49, adjusted p-value 0.0029) and Ruminococcaceae_2 (log-FC −2.69, adjusted p-value 1.98E-05).

In Mali infants, (**Table S3**), there were 2-fold increased differences observed for rice bran fed infants in Lactobacillus-0ut0356 (log-FC 3.2, adjusted p-value 1.35E-09) and decreased for Lachnospiraceae-unclassified-0tu0010 (log-FC −2.3, adjusted p-value 0.016). There were also distinctions among taxa between the infant gut microbiomes that was observed by age and country with respect to rice bran intake. At 12 months of age, the Nicaragua rice bran group had increased Paraprevotella (log-FC 6.2, adjusted p-value 4.27E-08), Phascolarctobacterium (log-FC 6.12, adjusted p-value 1.60E-08) Veillonella (log-FC 3.35, adjusted p-value 3.30E-07) and Bifidobacterium (log-FC 2.6, adjusted p-value 1.44E-05). Lower abundant taxa in rice bran infants at 12 months from Nicaragua were Lachnospiraceae_ND3007_group (log-FC −2.0, adjusted p-value 0.00029) and Alisonella (log-FC −4.0, adjusted p-value 1.60E-08). Malian rice bran fed infants at 12 months of age showed increased Lactobacillus_0tu0053 (log-FC 2.7, adjusted p-value 0.0098) and Alloprevotella (log-FC 3.6, adjusted p-value 0.00034). The significantly decreased fold difference in taxa of Malian infants between rice bran and control groups at 12 months were Bifidobacteriaceae_unclassified_0tu0265 (log-FC −2.6, adjusted p-value 3.95E-05) and Clostridium_sensu_stricto_1_0tu0076, and Terrisoporobacter (log-FC −2.1, adjusted p-value 7.03E-06).

There were 12 OTUs identified as rice bran responsive from this microbial community analysis that showed changes in both Mali and Nicaragua at either 8 or 12 months of age when compared to control. The highest area of overlap occurred for both Lactobacillaceae_Lactobacillus_Otu0024 and Lactobacillus_Otu0053 (**Table S4**). Other taxa that overlapped between countries with respect to rice bran intake included three distinct Bifidobacterium, Faecalibacterium, and Lachnospiriaceae.

### Metabolomics identified rice bran and microbial digested rice bran small molecules in stool of weaning infants

A total of 309 stool samples were collected from this 6-month prospective study to evaluate effects of rice bran supplementation compared to control infants from Nicaragua and Mali. ANOVA contrasts and Welch’s two-sample t-test were used to identify biochemicals that differed significantly between experimental groups at the 8-month time point. Stool metabolite analysis of children at 8 months of age in Nicaraguan and Malian infants resulted in the detection of 1449 biochemicals, of which 1016 metabolites had confirmed names and 433 compounds were of unknown structural identity (see **Table S5**). **Table 4** lists the fold differences calculated from the relative abundances of each stool metabolite between study diet groups at 8 months of age, whereby infants had been consuming rice bran daily for 2 months. There are 39 (Nicaragua) and 44 (Mali) stool metabolites with significant fold differences between children consuming rice bran compared to control. There were also 33 and 31 significantly different metabolites between groups that were classified as unknown for Nicaraguan and Malian infants respectively, (data shown in **Table S5**). Significant fold differences occurred for 15 amino acids, 2 peptides, 3 carbohydrates, 9 lipids, 1 cofactor and vitamin, and 9 xenobiotics (six of these considered as food components/plant-derived) in children consuming rice bran compared to control in Nicaragua. In Mali at 8 months of age, there were 6 amino acids, 1 energy, 14 lipids, 6 cofactor and vitamins, 5 nucleotides and 12 xenobiotics (7 classified as food components/ plant-derived) that showed significant differences in children consuming rice bran compared to control (**Table 4**).

**Table 4.**
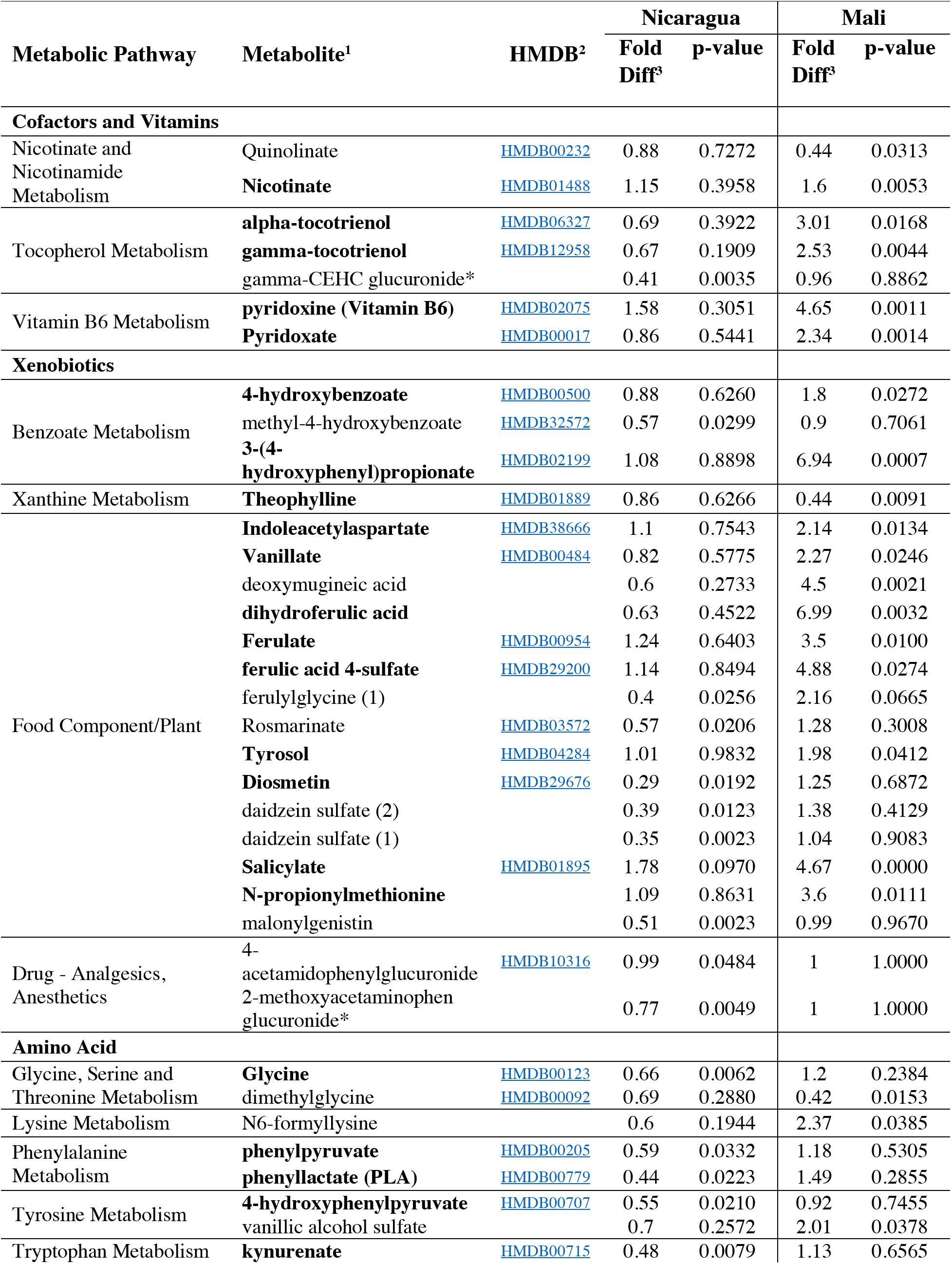
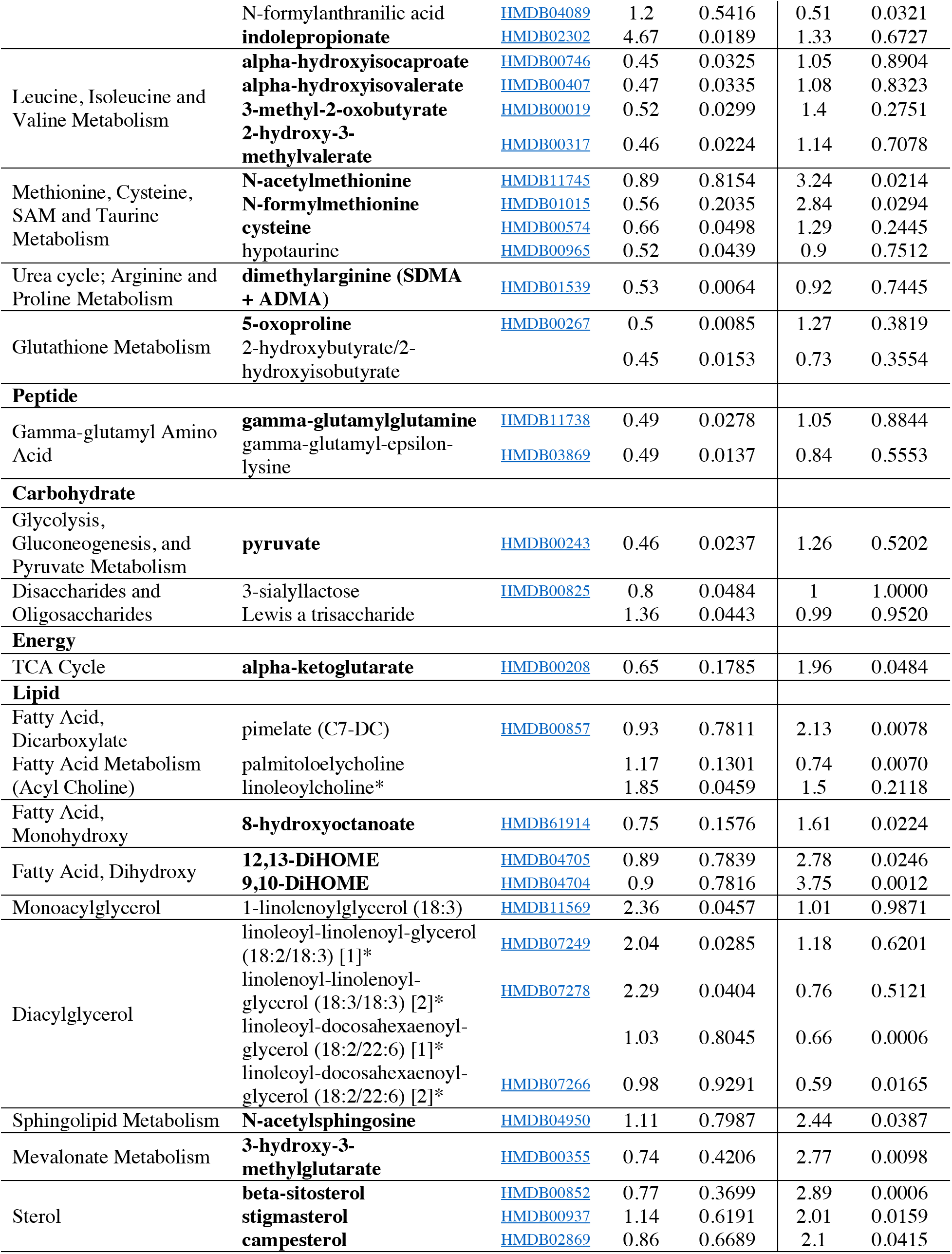
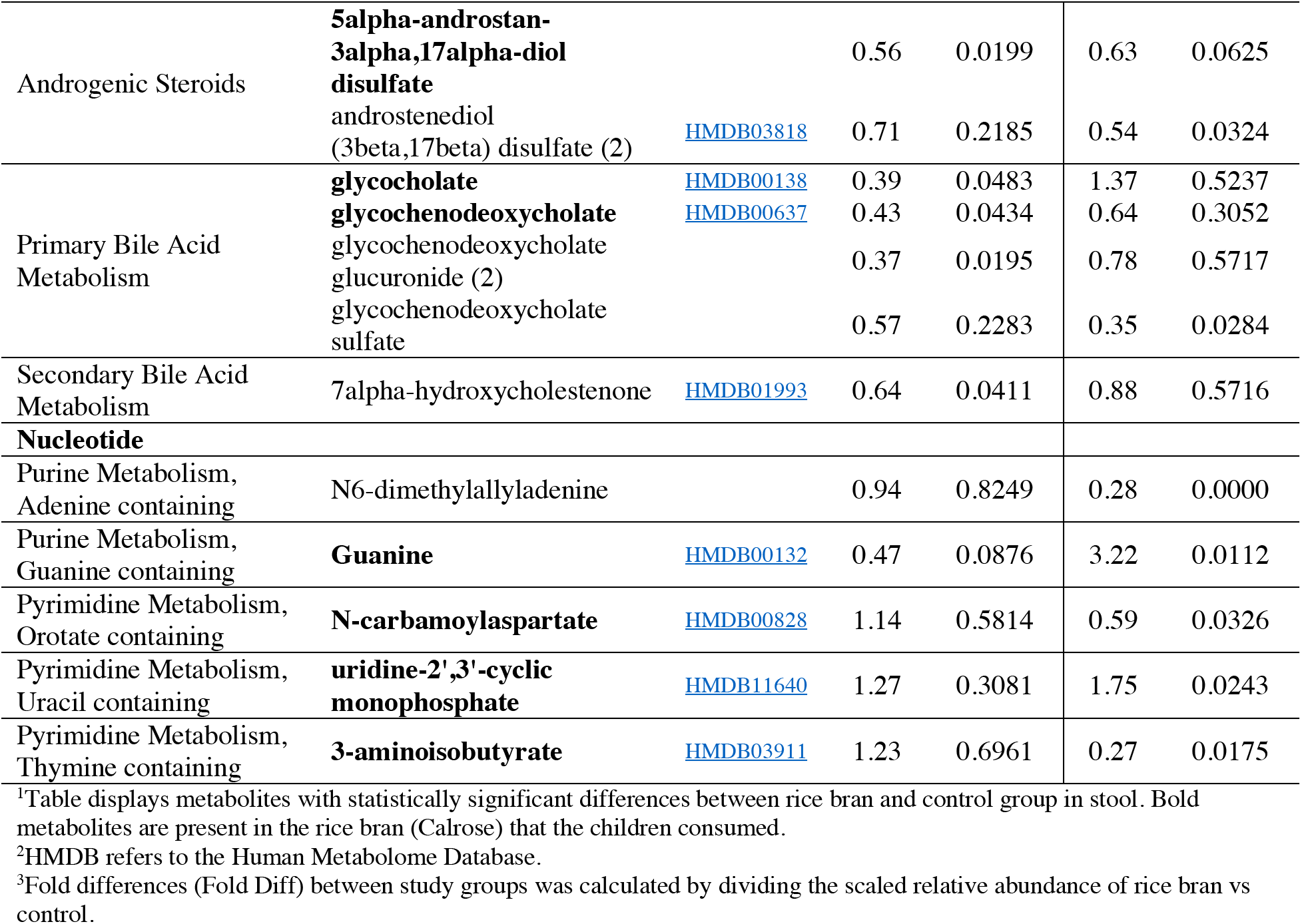
Stool metabolites significantly modulated by rice bran supplementation compared to control for Nicaragua & Mali infants at 8 months of age.

There were 62 stool metabolites from the Nicaraguan cohort at 8 months that showed significantly lower relative abundances (comparing fold differences between rice bran compared to control), and there were 10 significant stool metabolites with increased fold differences in abundance between groups. The stool metabolites of food and nutritional importance to highlight that resulted from increasing rice bran intake come from the tryptophan metabolism (indolepropionate), monoacylglycerol (1-linolenoylglycerol) and diacylglycerol metabolism (linoleoyl-linolenoyl-glycerol) pathways.

In contrast to Nicaragua, there were 54 distinct stool metabolites from the Mali cohort that had increased abundance and significant fold differences at 8 months between rice bran and control infants. Selected stool metabolites for gut health and nutrition relevance, as well as for originating from rice bran food metabolites, included Alpha-tocotrienol (vitamin E component), Pyridoxine (vitamin B6), Ferulic acid 4-sulfate, tyrosol, and N-acetyl sphingosine (**Table 4**). An estimated false discovery rate (q-value) was calculated to account for the multiple comparisons across metabolites that are typical of metabolomic-based studies.

## Discussion

This study demonstrated that rice bran supplementation was feasible, well tolerated, and safe for weaning infants with strong compliance to daily dietary intake in both LMIC countries. Rice bran supplementation in the diet supported growth of Nicaraguan and Malian infants with differences detected between groups by 8 months of age and improved gut permeability at 12 months. Nicaraguan infants fed rice bran had increased LAZ compared to the control, and in Mali, there was increased WAZ. Based on documented country-wide averages using the WHO scoring index^12^, the 95 healthy infants enrolled in this study had slightly higher WAZ, LAZ and WLZ scores.

There is a growing body of scientific evidence for a strong relationship between EED and growth deficits in children^7,10,47^. EED biomarkers were selected in this study because high concentrations of stool AAT and MPO were associated with decreased growth in children^47^, and Naylor et al. found that elevated AAT levels were associated with decreased oral rotavirus vaccine response^48^. Rice bran was shown to reduce AAT in neonatal pigs challenged with human Rotavirus infection^22^ and is of translational importance herein as rice bran intake improved AAT levels in Nicaraguan infants (**Table 2**). The lack of significant differences in levels of EED biomarkers for Mali may be due to the higher number of overall diarrheal episodes, the level of variability within individuals, and breastfeeding over time and across groups, yet these findings did concur with concentrations reported in the MAL-ED cohort^34^. EED biomarkers merit continuous review for relevance with growth outcomes due to extensive global variability in concentrations reported across studies^6,49–51^.

A study limitation and possible confounder of rice bran modulation to infant gut measures was that the percentage of exclusively breastfed infants was lower and delivery mode varied in the Nicaragua cohort (<50%) between groups, which was in contrast to the Mali site that had 100% of children that were breastfed and had vaginal delivery. These are key considerations to evaluating microbiomes of children with EED that have been characterized as less mature^52^, and with varied structure by geographical location^53^, diet deficiencies^54–58^, environmental exposures^53,59,60^ and host factors^59, 61^. Thus, we did expect changes in both the stool microbiome and metabolome that occurred over time with growth, and notably the microbial taxa and metabolites associated with the improved growth in rice bran groups also differed at 8 months in both countries.

Given that dietary rice bran intake has been previously shown to promote beneficial stool microbial communities, such as native gut probiotics in mice^23, 62^, pigs^21,22,29^ and adults^25,63,64^, we first evaluated significant genus level taxa differences between rice bran and control fed infants at 8 months and 12 months of age from both countries. In Nicaragua, Lactobacillus, Lachnospiraceae, Bifidobacterium, Ruminococcaceae and Veillonella were identified as responsive following rice bran consumption compared to an age and control matched group. These taxa have recognized saccharolytic mechanisms of action^65–67^, are known to produce and promote crossfeeding of short chain fatty acids^68, 69^, as well as provide competitive inhibition of pathogen colonization^70, 71^. The microbial enrichment of these communities as associated with infant growth outcomes has implications for assessing how gut microbes metabolize rice bran components. In Mali, the increased relative abundance of Lactobacillus in rice bran fed infants is also highly consistent with prior studies in young animals, and should be considered alongside evidence that human milk oligosaccharides also promote Lactobacillus in breastfed infants^72^.

Stool metabolites from infants fed rice bran showed significant fold differences amongst several essential amino acids, cofactors and vitamins, lipids, phytochemicals, and in energy metabolism pathways compared to the control groups. Stool metabolites originating from rice bran were identified in both Nicaragua and Malian infants, providing additional confirmation of compliance to the dietary intervention^73^. As predicted, we observed and reported exceptionally distinct profiles for both the stool microbiome and metabolome composition of infants between Mali and Nicaragua at all ages^40,53,74^ and therefore separately discussed the response to rice bran supplementation by region. For example, the nearly 5-fold increased stool detection of indolepropionate in rice bran infants compared to control from Nicaragua represents a tryptophan metabolite produced by the gut microbiota that may influence the developing immune system and intestinal homeostasis^75, 76^. Increased levels of N-acetylmethionine (nutritionally and metabolically equal to L-methionine) and N-formylmethionine in Malian infants also represented rice bran derived amino acids required for normal growth and development^77^. There are several cofactors and vitamins from rice bran supplementation, such as alpha and gamma-tocotrienol and pyridoxine (vitamin B6) that merit attention for demonstrating multiple health benefits such as synthesis of amino acids and neurotransmitter precursors, as well as preventing anemia and skin problems^73,78,79^. Additional microbial digested food components in the stool metabolome that come from rice bran were ferulic acid 4-sulfate, indoleacetylaspartate, and Tyrosol. A study limitation is that we had only supplemented for a 6 month window and according to WHO, growth assessment should be standardized and compared globally over the first two years of life^80, 81^. We put forth that rice bran metabolism by host and gut microbes between 12-36 months of age should be captured for the continuous assessment and influence on growth velocity during childhood^80^.

Furthermore, the increased separation noted between 8 and 12 months in this study showed modifications in gut microbial communities and metabolites by rice bran intake, and suggests there will be long-term impact on the overall microbiome composition as it continues to develop and mature^82^. The overall dietary pattern differences between mothers and infants weaning practices in each country were also considered a major source of variation as dietary diversity of gut microbe-food interactions were clearly observed in the stool metabolite profiling (see **Table S5**). Nevertheless, rice bran consumption was well tolerated at increasing dose supplementation amounts during the first 6 months of weaning without side effects or adverse interactions.

This was the first randomized controlled trial of rice bran supplementation in LMIC infants and provides compelling rationale for continued follow-up investigation of rice bran supplementation for reducing risk of malnutrition, as well as for eliciting changes during child growth that protect against enteric pathogens and diarrhea. The dose and feasibility outcomes from this study support development of rice bran based complementary weaning foods. Our findings also suggest that double blinded-controlled trial study designs with larger infant cohorts are warranted for long-term outcomes to be assessed until five years of age. Incorporating rice bran from local rice production and processing facilities should be a priority in subsequent trial designs, and with the goal of supporting the development of sustainable and affordable food products for weaning infants, particularly those residing in in LMIC where food and nutritional security remain.

## Materials and Methods

### Study design

A 6-month, prospective, randomized-controlled, dose escalation dietary intervention was conducted in a cohort of weaning infants residing in León, Nicaragua and in the community of Dioro, Mali, West Africa. Nicaraguan infants were recruited from public health rosters provided by the local Health Ministry from Perla Maria and Sutiava health sectors, and Malian infants were recruited from the Dioro Community Health Center. To be eligible, infants were screened between 4-5 months of age for health status, and then followed weekly for diarrhea episodes. Participants were excluded if they had experienced diarrhea or received antibiotic treatment within the previous month; had known allergies, or immune-compromising conditions (e.g. parasitic or malarial infections); had previously been hospitalized; and/or enrolled in a malnutrition treatment program. In Nicaragua, all eligible participants received 3 doses of the rotavirus vaccine per regular administration through the Immunization Program^83^. Rotavirus vaccination was not yet administrated to the Mali cohort. All Malian participants received vitamin A supplementation upon enrollment. Dietary intervention with rice bran started when infants were 6 months of age because WHO guidelines promote exclusive breastfeeding for the first six months of life^81, 84^.

The required ethical board reviews and approvals were completed for Mali and Nicaragua as provided by the Internal Review Board (IRB) of the Colorado State University Research Integrity and the Compliance Review office. In Mali, the Institut National de Recherche en Santé Publique (National Institute of Research in Public Health, FWA 00000892) approved the intervention, which occurred between October 2015 and May of 2016 and registered at clinicaltrial.gov as (NCT02557373) on 23 September 2015. Ethical review and approvals for the Nicaraguan intervention that occurred between March 2015 and 0ctober 2015 were provided by the IRBs of the Universidad Nacional Autónoma de Nicaragua – León, University of North Carolina at Chapel Hill, and Virginia Polytechnic Institute and State University and registered at clinicaltrial.gov on 26 November 2105 as (NCT02615886). Written informed consent was obtained from the infant’s parent or responsible guardian prior to any data collection. Infant participants that met the eligibility criteria were randomized within each health sector, and sex (Nicaragua) and geographic location of household and sex (Mali) to either rice bran or control group (see Fig. S1 for enrollment details). Randomization was completed using sequential enrollment for each site independently. Participants were randomized by CSU, enrolled and assigned to groups by study coordinators in each country. Complete study protocol is available online (http://csu-cvmbs.colostate.edu/academics/erhs/Pages/elizabeth-ryan-lab-global-health.aspx).

### Rice bran packaging for consumption

The United States Department of Agriculture-Agricultural Research Service (USDA-ARS) Dale Bumpers National Rice Research Center provided rice bran that was polished from the U.S. variety, Calrose. Rice bran is prone to fat oxidation and heat-stabilization was performed to increase shelf-life by heating the bran at 100 degrees Celsius for five minutes to inactivate the lipase/lipoxygenase enzymes that cause rancidity^85^. The rice bran was sifted to remove any debris (rice husk, rice grain). Packaging of the rice bran was completed by Western Innovations, Inc. (Denver, CO) where 22 kg of rice bran was weighed into 1g increments, separated into water-proof sachets, and heat-sealed to ensure the rice bran would be administered with accurate doses to infants.

Fourteen sachets (1g/sachet) were filled into a 4” x 3” x 2” box that was labeled for study participants and included nutrient information. These boxes were stored in a cool, dark, dry place until they were provided to study participants.

### Nicaragua and Mali intervention

The study team (doctor, nurse and study coordinator) in Nicaragua and the community health workers (CHWs) in Mali provided a 2-week supply of rice bran at each routine home visit and instructed the participant’s parent or guardian to add the daily amount of rice bran to the participant’s food. At 6-7 months of age, participants in the rice bran group consumed 1g of rice bran/day (1 sachet). Between the ages of 7-8 months, participants consumed 2g of rice bran/day (2 sachets). At 8-10 months of age, participants consumed 3g of rice bran/day (3 sachets). The amount increased to 4g of rice bran/day (4 sachets) from 10-11 months, and 5g of rice bran/day (5 sachets) from 11-12 months of age, respectively. The rice bran was added to appropriate weaning foods, such as rice cereal, yogurt, fruit and natural juices, vegetables, and soups. At the beginning of the intervention (six months of age), infant’s parents or guardians were instructed and monitored daily for one week by study personnel to ensure that guardians knew how to administer and record the amount of rice bran consumed. Compliance to the rice bran intervention was calculated from records that had the dose/amount of rice bran consumed circled in daily increments of none (0%), half (50%), or all (100%). The study team also collected any unused boxes or sachets during these visits. Participants in the control group did not receive any rice bran and there were no reports of brown rice intake during the 6-month study duration.

In Nicaragua, study personnel visited all infants weekly. In Mali, the CHWs visited each participant’s household daily for the duration of the 6-month study to assess compliance and diarrhea episodes. If a participant had a diarrhea episode, the study team would collect a stool sample, and collect information that included the diarrhea onset date, how long the episode lasted, numbers of bowel movements within 24 hours, any associated signs and symptoms (e.g. nausea, vomiting, fever), if any other family members had diarrhea, and if any treatment was provided (e.g. antibiotics, rehydration).

The study team in Nicaragua collected data for control group participants at 6, 8, and 12 months old, and rice bran group every month. The anthropometric measures (weight and length) were collected via a portable stadiometer and weighing balance. Mali participants visited the Community Health Clinic every month. Length was measured in supine position using a reclining length-board. Length was collected to the nearest centimeter and weight to the nearest 0.1 kg. Anthropometric measures were calculated for LAZ, WAZ, and WLZ scores following the World Health Organizations (WHO) child growth standards using the WHO Anthro software (version 3.2.2)^86^.

Diapers were provided to all study participants. Stool was collected directly from soiled diapers. Freshly collected stool was diluted 20-fold and homogenized in a sterile pre-reduced anaerobic saline – 0.1 M potassium phosphate buffer (pH 7.2) containing 20% glycerol (vol/vol). Four aliquot suspensions were prepared in 15 mL falcon tubes, transported on dry ice to the UNAN-León-Center of Infectious Diseases Laboratories (and liquid nitrogen in Bamako, Mali), immediately transferred to a −80°C freezer, shipped in a liquid nitrogen chilled dry shipping dewar to Colorado State University, where they were relocated into a −80°C freezer prior to analysis.

A study questionnaire was completed by the participant’s caretaker (e.g. mother, father, or grandparent) to assess for duration of breastfeeding, types of and timing of introductions to complementary foods, as well as antibiotic use. The breastfeeding questions included whether or not the child was receiving breast milk, and/or had the child been receiving received formula. The complementary feeding history included a list of common Nicaraguan and Malian foods that are normally introduced to infants during weaning. Infant’s parents or guardians recorded how often the infant consumed each of the eleven foods. The questionnaire also recorded if a participant had received treatment with antibiotics since the last visit, the reason for taking the antibiotic, the name of the antibiotic, as well as the length of time the participant had been taking the antibiotic. A household survey was also completed at the beginning of the trial to collect mother’s education level, drinking water source, household flooring type, and animals present in the household. Analysis of breastfeeding and formula feeding patterns, complementary feeding practices, and associations with nutritional status at 6-months old (i.e. baseline) were previously reported for Nicaragua^87^. Monthly visits to the Mali community health clinic provided monitoring for malnutrition and severe adverse events; no adverse events were reported in the rice bran intervention group. There was one participant death reported in the control group (respiratory infection) and another withdrew to receive malnutrition treatment in the second month of the study. Diarrheal episodes were recorded, and a sample was collected in both countries using the same protocol.

### Stool analysis for EED markers

Stool biomarkers were selected to report gut inflammation and epithelial integrity as indicators of EED. These included neopterin (NEO), myeloperoxidase (MPO), calprotectin, (CAL) and alpha-1 antitrypsin (AAT)^88^. Suspended stool samples from 6, 8, and 12-month collections were centrifuged at 3,000 RPM to remove debris, following agitation, and the remaining supernatant was used for Enzyme-Linked-Immunosorbant-Assay (ELISA) determination of EED biomarker concentrations. Laboratory analysis protocols included in commercial kits were followed. Concentrations of CAL were determined at a 1:360 final dilution factor (Eagle Biosciences-Nashua, NH. Ref: CAL35-K01). Samples were diluted to 1:100 for determination of NE0 concentrations (GenWay Biotech Inc-San Diego, CA, USA). MP0 concentrations were determined at a 1:500 dilution factor (Immundiagnostik AG-Bensheim, Germany). Samples were diluted to 1:12,500 for determination of AAT concentrations (Immuchrom GMBH-Heppenheim, Germany), and dilution factors accounted for stool suspension ratios (20-fold). Final concentrations were determined from averages of replicate assays and duplicate optical density readings, and interpolated using Graphpad 6.0 according to standards measured on each 96-well plate.

### Stool microbiome analysis for Nicaragua and Mali

The infant stool was collected at 6, 8 and 12 months of age from diapers and placed in a 1:19 ratio with Phosphate Buffered Saline + Glycerol solution. Diarrhea samples were collected using the same protocol. Suspended stool samples were vortexed before centrifuging at 3000 RPM to separate the stool debris. The remaining supernatant was used for Enzyme-Linked-Immunosorbant-Assay (ELISA) determination of EED biomarkers whereas; DNA was extracted for 16S microbial analysis from the stool pellet. DNA extraction was conducted using MoBio PowerSoil Kit (Reference number 12888, MoBio Laboratories Inc., Solana Beach, CA). PCR amplification of 390 bp amplicons was done in 50 μl reaction using Fischer Hot Start Master Mix and EMP standard protocols^42–46^. SPRI magnetic beads were used to purify DNA, and flourimetric quantification of Sybr Green tags was used to confirm adequate concentration of DNA. The pooled library was created with 50 ng DNA per sample and quantified using Kapa Kit (Kapa Biosystems). The pooled library was run on Illumina-MiSeq with 15% PhiX mock library to reduce discrepancies in read clustering, using the Illumina V2 500 cycle kit (2 × 250/250 paired-end reads).

### Microbiome data processing and analysis

Sequence data were processed using mothur^89^ version 1.39.5 and using a custom pipeline that provides an adjustment on the developers’ standard operating procedure (SOP) for OTU calling and taxonomic classification of MiSeq data first presented in Kozich, et al., 2013^90^. For alignment and classification within this SOP we used the SILVA database^91^ version 128. Clustering, for OTU identification, was performed using VSEARCH using the distance based greedy clustering (DGC) option as implanted in mothur and utilizing 0.97 sequence similarity cutoff. We also used a cutoff of one read that was subtracted from all OTU read counts to guard against overestimation of sample richness. Rarefaction curves were generated using the package vegan^92^ as implemented in R version 3.4.4^93^ to assess diversity and suitability of depth of coverage per sample. The resulting OTU table was utilized in further data analyses as follows.

Exploring the data: Bar-graphs for relative abundance data based on the resulting OTU table were generated using the ggplot2^94^ package in R. These plots were generated for the data at the genus and the family levels and meant to describe the microbial community structure per sampled infant and per time point under each of the treatment levels.

Data were normalized using cumulative sum scaling (CSS)^95^ prior to beta diversity and log-fold change analyses. Nonmetric Multidimensional Scaling (NMDS)^96^ was used on the 0TU level to assess possible trends and clustering in the microbial community structure comparing the two countries, the treatment conditions and the two time points, using the vegan package and utilizing Bray-Curtis dissimilarity^96^. Data were separated per country and the metagenomeSeq^97^ package in R^93^ was used to fit a zero inflated normal (ZIN) model to test for log-fold change differences between the rice bran treatment and control per age group. Benjamini and Hochberg’s^98^ false discovery rate (FDR) method was used to correct for multiple testing and compute the adjusted p-values used to determine significance of differences in the log-fold change of OTU abundance.

### Stool metabolomics analysis

Stool samples were sent to Metabolon Inc. (Durham, NC, USA) for non-targeted metabolite profiling via ultrahigh-performance liquid chromatography tandem mass spectrometry (UPLC-MS/MS). All samples were accessioned into the Metabolon Library Information Management Systems (LIMS) and prepared using the automated MicroLab Star^®^ system (Hamilton Company, Switzerland). Eight to ten recovery standards were added prior to the first step in the extraction process for quality control purposes. Extraction was performed using 80% ice-cold methanol under vigorous shaking for 2 min (Glen Mills GenoGrinder 2000) followed by centrifugation to remove protein, dissociate small molecules bound to protein or trapped in the precipitated protein matrix. Each stool extract was divided into five fractions: two for analysis by two separate reverse phase UPLC-MS/MS methods with positive ion mode electrospray ionization, 1 for analysis by reverse phase UPLC-MS/MS methods with negative ion mode electrospray ionization, 1 for hydrophilic interaction liquid chromatography UPLC-MS/MS with negative ion mode electrospray ionization, and 1 sample for backup. All samples were placed briefly on Concentration Evaporator (TurboVap^®^ Zymark) to remove organic solvent. UPLC-MS/MS methods utilized a Waters ACQUITY ultra-performance liquid chromatography and a Thermo Scientific Q-Exactive high resolution/accurate mass spectrometer interfaced with a heated electrospray ionization (HESI-II) source and Orbitrap mass analyzer operated at 35,000 mass resolution. Raw data was extracted, peak-identified and processed for quality control using Metabolon’s hardware and software.

### Statistical analysis

Statistical analyses for anthropometric measures (length, weight, LAZ, WAZ, and WLZ) and stool EED biomarkers were completed using SAS 9.4 (Cary, NC, USA). The sample size was calculated for achieving greater than 85% power and based on expected changes in selected stool metabolites following dietary rice bran consumption for one month^25^. Normality was evaluated by visual inspection. For anthropometric variables, two-sample t-tests were used to compare means for the 2 treatment groups (rice bran and control) separately at birth and 6 months (prior to start of treatment). A repeated measures analysis was performed for each response variable separately using SAS Proc Mixed. Specifically, treatment (rice bran or control) and age (6, 8 or 12 months), and treatment-age interaction were included in the model as fixed effects. The participant was included as a random effect to account for repeated measures. At each age, treatment groups were compared using contrasts of the model. A similar repeated measures analysis was used EED biomarkers, but log transformation was used to satisfy model assumptions. For stool metabolites, Welch’s two-sample t-test was used to analyze statistical significance between groups’ stool metabolites, after participating in the 6-month dietary trial. A p-value of ≤0.05 was used for statistical significance. An estimated false discovery rate (q-value) was calculated to account for the multiple comparisons across metabolites that are typical of metabolomic-based studies.

## Acknowledgements

The authors thank the study participants, community health workers, and local clinical staff for their assistance in the conduct of the clinical trials.

## Author contributions

For Nicaragua trial: SV, SBD, LY and EPR designed the research trial and maintained study oversight. ECB, SV, JP, CP and LZ conducted research in field (including sample collection and processing), and IZ, SM, JP, HI and CP analyzed stool samples in the lab. For Mali trial: EPR, OK, and AM designed the approved research protocols for the rice bran intervention with assistance from AB, AC, ECB, and LS. AB and AC coordinated the study in the community, and LS oversaw sample collection and field laboratory operations. LD, SD, and KK provided community and participant engagement, collected samples and data, and carried out laboratory analyses. SM, LZ, IZ, ECB, HI, ZA and LZ carried out further laboratory and data analyses in the USA. AH, ZA, HI and LZ compiled the growth, biomarkers, microbial communities and metabolomics datasets and performed integrated statistical data analysis for both Mali and Nicaragua sites. LZ, ECB, SM, HI, SV, OK, and EPR collectively interpreted data, carried out the literature search, constructed tables and figures, and contributed to manuscript preparation. LZ, ECB, SM, HI, ZA and EPR wrote the paper. EPR had primary responsibility for the final product. All authors read and approved the final manuscript. The authors declare that they have no competing interests.

## Data and materials availability

*16S* sequence data were submitted to the National Center for Biotechnology Information SRA under accession no. (SRP159269) and Bio-project (PRJNA488807).

