## Supplementary tables and figures for "Rice bran supplementation modulates growth, microbiome and metabolome in weaning infants: a clinical trial in Nicaragua and Mali"

**Title:**

**Authors and Affiliations:**

Luis E. Zambrana[1,2], Starin McKeen[1], Hend Ibrahim[1,3], Iman Zarei[1], Erica C. Borresen[1], Lassina Doumbia[4], Abdoulaye Bore[4], Alima Cissoko[4], Seydou Douyon[4], Karim Kone[4], Johann Perez[2], Claudia Perez[2], Ann Hess[5], Zaid Abdo[6], Lansana Sangare[4], Ababacar Maiga[4], Sylvia Becker-Dreps[7], Lijuan Yuan[8], Ousmane Koita[4*], Samuel Vilchez[2*], & Elizabeth P. Ryan[1]*

**Fig. S1.** Overview of study design for both Nicaragua & Mali cohorts. A diagram of the eligibility criteria, intervention, participant information, and analysis of samples.

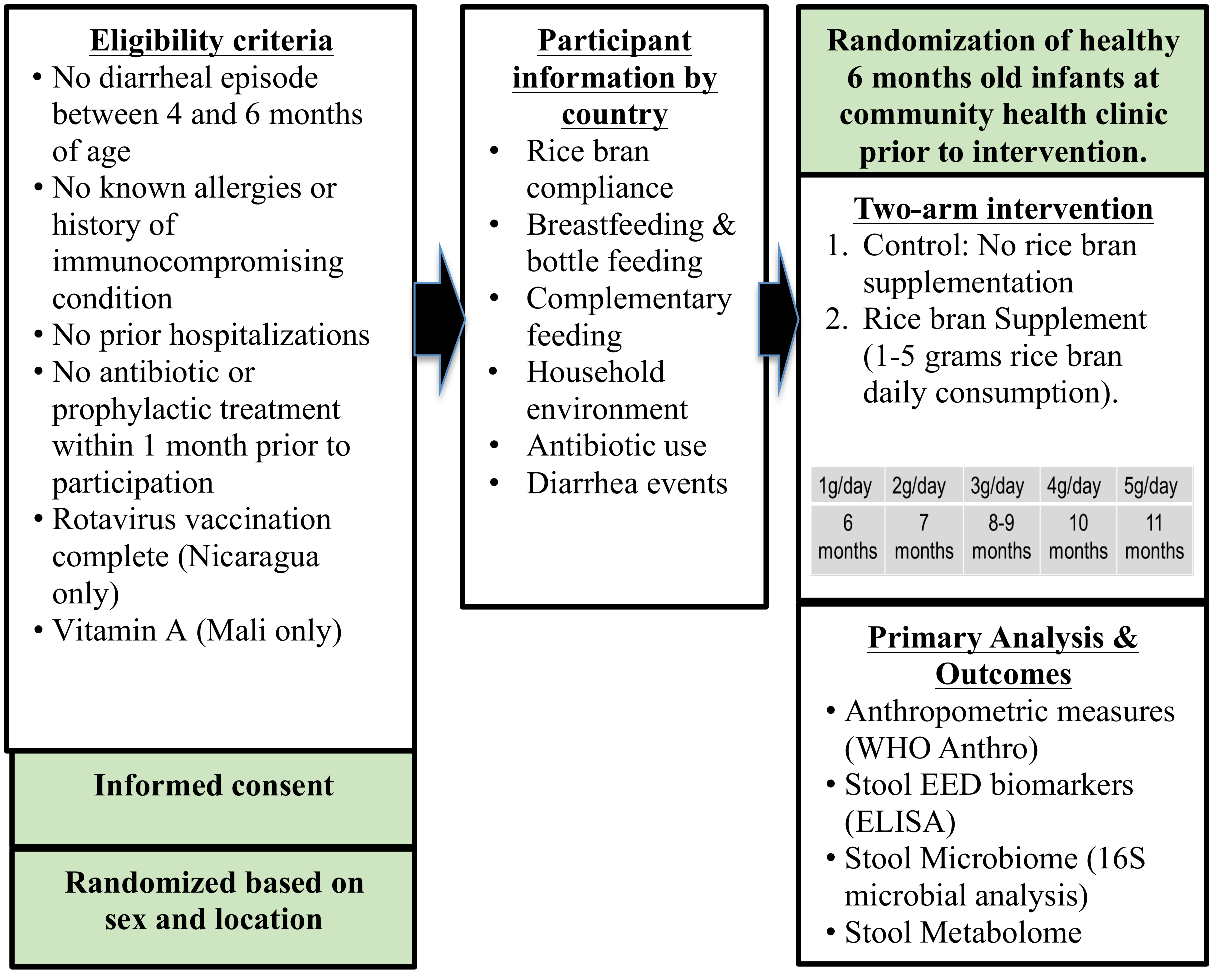

### Supplemental Tables

**Table S1.** The alpha diversity indices for gut microbial communities analyzed from Nicaraguan and Malian Infants at 8 and 12 months of age (data shown for all control and rice bran infants).

| **Nicaragua_Control** | | | | | | | | |
|---|---|---|---|---|---|---|---|---|
| **SampleID** | **country** | **group** | **sex** | **month** | **Observed** | **Shannon** | **InvSimpson** | **Richness** |
| COSU00591 | Nicaraguan | Control | F | 8 | 118 | 2.35 | 6.09 | 66.75 |
| COSU00592 | Nicaraguan | Control | F | 8 | 134 | 2.39 | 7.23 | 78.37 |
| COSU00594 | Nicaraguan | Control | F | 8 | 293 | 1.95 | 3.83 | 86.18 |
| COSU00596 | Nicaraguan | Control | F | 8 | 173 | 2.40 | 4.75 | 110.96 |
| COSU00603 | Nicaraguan | Control | F | 8 | 116 | 2.73 | 8.18 | 86.90 |
| COSU00606 | Nicaraguan | Control | F | 8 | 95 | 2.22 | 6.27 | 53.84 |
| COSU00608 | Nicaraguan | Control | F | 8 | 105 | 2.67 | 10.41 | 73.89 |
| COSU00610 | Nicaraguan | Control | F | 8 | 114 | 1.88 | 3.72 | 81.72 |
| COSU00611 | Nicaraguan | Control | F | 8 | 72 | 1.85 | 4.22 | 51.73 |
| COSU00593 | Nicaraguan | Control | M | 8 | 96 | 2.16 | 4.90 | 56.81 |
| COSU00595 | Nicaraguan | Control | M | 8 | 132 | 2.21 | 4.02 | 78.70 |
| COSU00597 | Nicaraguan | Control | M | 8 | 140 | 3.11 | 13.28 | 95.60 |
| COSU00598 | Nicaraguan | Control | M | 8 | 89 | 1.92 | 4.22 | 57.26 |
| COSU00599 | Nicaraguan | Control | M | 8 | 94 | 1.93 | 4.48 | 52.51 |
| COSU00600 | Nicaraguan | Control | M | 8 | 77 | 1.95 | 5.09 | 52.17 |
| COSU00601 | Nicaraguan | Control | M | 8 | 100 | 2.51 | 7.08 | 83.89 |
| COSU00602 | Nicaraguan | Control | M | 8 | 92 | 2.21 | 5.32 | 68.75 |
| COSU00604 | Nicaraguan | Control | M | 8 | 98 | 2.26 | 5.20 | 83.26 |
| COSU00605 | Nicaraguan | Control | M | 8 | 132 | 2.49 | 6.41 | 103.44 |
| COSU00607 | Nicaraguan | Control | M | 8 | 178 | 2.64 | 6.48 | 115.70 |
| COSU00609 | Nicaraguan | Control | M | 8 | 105 | 1.97 | 3.97 | 58.00 |
| COSU00612 | Nicaraguan | Control | M | 8 | 96 | 1.86 | 3.57 | 58.24 |
| COSU00635 | Nicaraguan | Control | F | 12 | 156 | 2.52 | 7.02 | 88.29 |
| COSU00636 | Nicaraguan | Control | F | 12 | 99 | 1.79 | 2.81 | 72.23 |
| COSU00638 | Nicaraguan | Control | F | 12 | 139 | 2.22 | 5.89 | 80.13 |
| COSU00640 | Nicaraguan | Control | F | 12 | 165 | 2.69 | 7.34 | 114.71 |
| COSU00646 | Nicaraguan | Control | F | 12 | 194 | 2.53 | 5.04 | 117.05 |
| COSU00649 | Nicaraguan | Control | F | 12 | 115 | 1.88 | 3.15 | 70.62 |
| COSU00651 | Nicaraguan | Control | F | 12 | 141 | 2.88 | 9.77 | 93.71 |

| | | | | | | | | |
|---|---|---|---|---|---|---|---|---|
| COSU00653 | Nicaraguan | Control | F | 12 | 116 | 2.91 | 10.95 | 88.02 |
| COSU00654 | Nicaraguan | Control | F | 12 | 75 | 1.73 | 3.20 | 54.28 |
| COSU00639 | Nicaraguan | Control | M | 12 | 157 | 2.89 | 9.89 | 107.53 |
| COSU00641 | Nicaraguan | Control | M | 12 | 160 | 2.70 | 6.19 | 115.84 |
| COSU00643 | Nicaraguan | Control | M | 12 | 96 | 2.34 | 5.54 | 61.45 |
| COSU00644 | Nicaraguan | Control | M | 12 | 103 | 2.39 | 6.27 | 79.01 |
| COSU00645 | Nicaraguan | Control | M | 12 | 103 | 2.41 | 6.58 | 78.79 |
| COSU00647 | Nicaraguan | Control | M | 12 | 231 | 3.76 | 26.09 | 168.33 |
| COSU00648 | Nicaraguan | Control | M | 12 | 136 | 2.59 | 9.13 | 86.78 |
| COSU00650 | Nicaraguan | Control | M | 12 | 181 | 2.88 | 5.64 | 124.09 |
| COSU00652 | Nicaraguan | Control | M | 12 | 151 | 3.03 | 10.07 | 91.66 |
| COSU00655 | Nicaraguan | Control | M | 12 | 104 | 2.14 | 3.88 | 65.58 |
| **Nicaragua_Rice bran** | | | | | | | | |
| **SampleID** | **country** | **group** | **sex** | **month** | **Observed** | **Shannon** | **InvSimpson** | **Richness** |
| COSU00613 | Nicaraguan | Rice_bran | F | 8 | 204 | 2.70 | 7.91 | 115.86 |
| COSU00614 | Nicaraguan | Rice_bran | F | 8 | 150 | 2.54 | 8.62 | 94.77 |
| COSU00616 | Nicaraguan | Rice_bran | F | 8 | 78 | 1.40 | 2.44 | 50.26 |
| COSU00620 | Nicaraguan | Rice_bran | F | 8 | 192 | 2.59 | 7.26 | 99.33 |
| COSU00621 | Nicaraguan | Rice_bran | F | 8 | 62 | 1.93 | 4.98 | 50.53 |
| COSU00622 | Nicaraguan | Rice_bran | F | 8 | 140 | 1.56 | 2.92 | 69.77 |
| COSU00623 | Nicaraguan | Rice_bran | F | 8 | 152 | 2.22 | 4.02 | 91.98 |
| COSU00629 | Nicaraguan | Rice_bran | F | 8 | 162 | 2.76 | 8.76 | 110.76 |
| COSU00631 | Nicaraguan | Rice_bran | F | 8 | 206 | 2.08 | 3.71 | 103.54 |
| COSU00634 | Nicaraguan | Rice_bran | F | 8 | 123 | 2.37 | 7.71 | 69.00 |
| COSU00615 | Nicaraguan | Rice_bran | M | 8 | 102 | 1.99 | 3.51 | 64.13 |
| COSU00617 | Nicaraguan | Rice_bran | M | 8 | 115 | 2.55 | 7.11 | 80.05 |
| COSU00618 | Nicaraguan | Rice_bran | M | 8 | 119 | 2.13 | 5.82 | 75.76 |
| COSU00619 | Nicaraguan | Rice_bran | M | 8 | 167 | 2.11 | 3.98 | 83.64 |
| COSU00624 | Nicaraguan | Rice_bran | M | 8 | 177 | 2.56 | 8.08 | 92.85 |
| COSU00625 | Nicaraguan | Rice_bran | M | 8 | 54 | 1.59 | 3.41 | 39.47 |
| COSU00626 | Nicaraguan | Rice_bran | M | 8 | 351 | 3.20 | 12.16 | 187.50 |
| COSU00627 | Nicaraguan | Rice_bran | M | 8 | 104 | 0.83 | 1.50 | 48.07 |
| COSU00628 | Nicaraguan | Rice_bran | M | 8 | 710 | 2.11 | 4.92 | 97.68 |
| COSU00630 | Nicaraguan | Rice_bran | M | 8 | 213 | 2.64 | 7.07 | 102.74 |
| COSU00632 | Nicaraguan | Rice_bran | M | 8 | 72 | 1.36 | 2.64 | 45.41 |
| COSU00633 | Nicaraguan | Rice_bran | M | 8 | 210 | 2.66 | 8.89 | 109.11 |
| COSU00656 | Nicaraguan | Rice_bran | F | 12 | 154 | 2.35 | 6.15 | 91.10 |
| COSU00657 | Nicaraguan | Rice_bran | F | 12 | 225 | 3.01 | 9.53 | 134.07 |
| COSU00659 | Nicaraguan | Rice_bran | F | 12 | 138 | 2.62 | 8.69 | 90.78 |

| | | | | | | | | |
|---|---|---|---|---|---|---|---|---|
| COSU00663 | Nicaraguan | Rice_bran | F | 12 | 203 | 3.13 | 11.36 | 126.47 |
| COSU00664 | Nicaraguan | Rice_bran | F | 12 | 137 | 2.02 | 5.32 | 83.94 |
| COSU00665 | Nicaraguan | Rice_bran | F | 12 | 174 | 2.42 | 5.25 | 91.34 |
| COSU00666 | Nicaraguan | Rice_bran | F | 12 | 118 | 2.09 | 4.46 | 69.68 |
| COSU00672 | Nicaraguan | Rice_bran | F | 12 | 247 | 3.21 | 15.51 | 132.12 |
| COSU00674 | Nicaraguan | Rice_bran | F | 12 | 96 | 1.04 | 1.80 | 47.66 |
| COSU00677 | Nicaraguan | Rice_bran | F | 12 | 163 | 2.39 | 5.78 | 95.68 |
| COSU00678 | Nicaraguan | Rice_bran | F | 12 | 210 | 2.32 | 5.87 | 95.95 |
| COSU00658 | Nicaraguan | Rice_bran | M | 12 | 100 | 1.72 | 3.01 | 61.75 |
| COSU00660 | Nicaraguan | Rice_bran | M | 12 | 185 | 2.94 | 9.95 | 127.45 |
| COSU00661 | Nicaraguan | Rice_bran | M | 12 | 163 | 2.47 | 7.35 | 99.53 |
| COSU00662 | Nicaraguan | Rice_bran | M | 12 | 121 | 2.24 | 6.16 | 79.40 |
| COSU00667 | Nicaraguan | Rice_bran | M | 12 | 256 | 2.81 | 6.65 | 147.79 |
| COSU00668 | Nicaraguan | Rice_bran | M | 12 | 117 | 2.14 | 4.10 | 81.01 |
| COSU00669 | Nicaraguan | Rice_bran | M | 12 | 248 | 2.89 | 9.92 | 133.17 |
| COSU00670 | Nicaraguan | Rice_bran | M | 12 | 281 | 2.52 | 5.57 | 128.36 |
| COSU00671 | Nicaraguan | Rice_bran | M | 12 | 207 | 2.49 | 7.50 | 109.91 |
| COSU00673 | Nicaraguan | Rice_bran | M | 12 | 207 | 2.55 | 7.16 | 101.09 |
| COSU00675 | Nicaraguan | Rice_bran | M | 12 | 108 | 1.78 | 3.81 | 64.76 |
| COSU00676 | Nicaraguan | Rice_bran | M | 12 | 198 | 2.77 | 10.24 | 103.28 |
| **Mali_Control** | | | | | | | | |
| **SampleID** | **country** | **group** | **sex** | **month** | **Observed** | **Shannon** | **InvSimpson** | **Richness** |
| COSU00720 | Malian | Control | F | 8 | 76 | 1.66 | 3.28 | 49.63 |
| COSU00722 | Malian | Control | F | 8 | 53 | 1.64 | 2.60 | 46.01 |
| COSU00723 | Malian | Control | F | 8 | 71 | 0.49 | 1.18 | 32.87 |
| COSU00724 | Malian | Control | F | 8 | 44 | 0.97 | 1.57 | 35.66 |
| COSU00728 | Malian | Control | F | 8 | 67 | 1.73 | 2.71 | 59.74 |
| COSU00730 | Malian | Control | F | 8 | 78 | 1.50 | 2.20 | 52.32 |
| COSU00734 | Malian | Control | F | 8 | 52 | 1.67 | 4.22 | 42.21 |
| COSU00739 | Malian | Control | F | 8 | 39 | 0.75 | 1.41 | 28.54 |
| COSU00740 | Malian | Control | F | 8 | 103 | 1.93 | 3.01 | 63.90 |
| COSU00721 | Malian | Control | M | 8 | 54 | 1.69 | 3.47 | 49.76 |
| COSU00725 | Malian | Control | M | 8 | 51 | 1.81 | 3.50 | 45.99 |
| COSU00726 | Malian | Control | M | 8 | 137 | 3.09 | 14.13 | 107.16 |
| COSU00727 | Malian | Control | M | 8 | 80 | 1.38 | 1.94 | 54.51 |
| COSU00729 | Malian | Control | M | 8 | 53 | 1.14 | 1.69 | 44.37 |
| COSU00731 | Malian | Control | M | 8 | 66 | 1.59 | 2.39 | 55.52 |
| COSU00732 | Malian | Control | M | 8 | 96 | 1.60 | 2.31 | 58.47 |
| COSU00733 | Malian | Control | M | 8 | 104 | 1.76 | 3.28 | 66.50 |

| | | | | | | | | |
|---|---|---|---|---|---|---|---|---|
| COSU00735 | Malian | Control | M | 8 | 97 | 1.93 | 3.56 | 56.55 |
| COSU00736 | Malian | Control | M | 8 | 145 | 2.13 | 3.63 | 84.18 |
| COSU00737 | Malian | Control | M | 8 | 41 | 1.09 | 2.01 | 28.40 |
| COSU00738 | Malian | Control | M | 8 | 122 | 2.14 | 3.72 | 78.20 |
| COSU00763 | Malian | Control | F | 12 | 47 | 1.89 | 3.92 | 44.70 |
| COSU00765 | Malian | Control | F | 12 | 102 | 2.11 | 4.23 | 71.04 |
| COSU00766 | Malian | Control | F | 12 | 109 | 2.29 | 4.14 | 78.86 |
| COSU00767 | Malian | Control | F | 12 | 75 | 1.94 | 3.99 | 57.87 |
| COSU00772 | Malian | Control | F | 12 | 81 | 1.98 | 3.33 | 56.28 |
| COSU00774 | Malian | Control | F | 12 | 74 | 1.88 | 4.09 | 58.95 |
| COSU00778 | Malian | Control | F | 12 | 115 | 2.52 | 8.52 | 84.21 |
| COSU00782 | Malian | Control | F | 12 | 71 | 1.62 | 2.78 | 66.42 |
| COSU00783 | Malian | Control | F | 12 | 195 | 2.35 | 5.48 | 117.78 |
| COSU00784 | Malian | Control | F | 12 | 108 | 1.89 | 3.16 | 78.65 |
| COSU00785 | Malian | Control | F | 12 | 187 | 2.65 | 6.77 | 118.63 |
| COSU00764 | Malian | Control | M | 12 | 80 | 2.08 | 4.64 | 60.53 |
| COSU00768 | Malian | Control | M | 12 | 104 | 2.49 | 6.19 | 83.79 |
| COSU00770 | Malian | Control | M | 12 | 132 | 2.15 | 4.42 | 88.75 |
| COSU00771 | Malian | Control | M | 12 | 100 | 2.35 | 6.71 | 70.71 |
| COSU00773 | Malian | Control | M | 12 | 90 | 1.20 | 1.73 | 60.06 |
| COSU00775 | Malian | Control | M | 12 | 64 | 1.66 | 3.28 | 49.08 |
| COSU00776 | Malian | Control | M | 12 | 112 | 2.29 | 5.06 | 78.40 |
| COSU00777 | Malian | Control | M | 12 | 131 | 2.49 | 8.19 | 84.32 |
| COSU00779 | Malian | Control | M | 12 | 149 | 2.61 | 6.23 | 106.79 |
| COSU00780 | Malian | Control | M | 12 | 131 | 1.92 | 3.12 | 92.67 |
| COSU00781 | Malian | Control | M | 12 | 141 | 2.20 | 5.19 | 77.19 |
| **Mali_Rice bran** | | | | | | | | |
| **SampleID** | **country** | **group** | **sex** | **month** | **Observed** | **Shannon** | **InvSimpson** | **Richness** |
| COSU00744 | Malian | Rice_bran | F | 8 | 44 | 1.03 | 1.68 | 32.95 |
| COSU00745 | Malian | Rice_bran | F | 8 | 93 | 1.87 | 3.79 | 77.37 |
| COSU00746 | Malian | Rice_bran | F | 8 | 115 | 1.64 | 2.29 | 84.78 |
| COSU00747 | Malian | Rice_bran | F | 8 | 48 | 1.36 | 2.02 | 37.41 |
| COSU00749 | Malian | Rice_bran | F | 8 | 78 | 1.91 | 4.20 | 56.14 |
| COSU00750 | Malian | Rice_bran | F | 8 | 92 | 2.25 | 3.93 | 72.36 |
| COSU00755 | Malian | Rice_bran | F | 8 | 62 | 2.16 | 5.32 | 53.12 |
| COSU00756 | Malian | Rice_bran | F | 8 | 111 | 2.40 | 7.05 | 74.61 |
| COSU00758 | Malian | Rice_bran | F | 8 | 44 | 0.92 | 1.49 | 32.33 |
| COSU00759 | Malian | Rice_bran | F | 8 | 108 | 1.66 | 2.29 | 65.16 |
| COSU00741 | Malian | Rice_bran | M | 8 | 70 | 1.90 | 4.07 | 48.68 |

| | | | | | | | | |
|---|---|---|---|---|---|---|---|---|
| COSU00742 | Malian | Rice_bran | M | 8 | 51 | 1.55 | 2.72 | 38.98 |
| COSU00748 | Malian | Rice_bran | M | 8 | 85 | 1.96 | 3.17 | 67.81 |
| COSU00751 | Malian | Rice_bran | M | 8 | 60 | 1.23 | 1.98 | 49.58 |
| COSU00752 | Malian | Rice_bran | M | 8 | 60 | 1.60 | 2.56 | 48.05 |
| COSU00753 | Malian | Rice_bran | M | 8 | 62 | 1.49 | 2.07 | 49.66 |
| COSU00754 | Malian | Rice_bran | M | 8 | 107 | 2.22 | 5.14 | 75.16 |
| COSU00757 | Malian | Rice_bran | M | 8 | 156 | 2.07 | 5.11 | 73.08 |
| COSU00760 | Malian | Rice_bran | M | 8 | 47 | 2.12 | 5.93 | 41.35 |
| COSU00761 | Malian | Rice_bran | M | 8 | 96 | 1.99 | 3.09 | 72.28 |
| COSU00789 | Malian | Rice_bran | F | 12 | 95 | 2.09 | 4.29 | 69.27 |
| COSU00790 | Malian | Rice_bran | F | 12 | 99 | 2.08 | 4.51 | 74.15 |
| COSU00791 | Malian | Rice_bran | F | 12 | 384 | 2.21 | 4.85 | 96.42 |
| COSU00792 | Malian | Rice_bran | F | 12 | 110 | 2.03 | 2.98 | 88.57 |
| COSU00793 | Malian | Rice_bran | F | 12 | 39 | 1.23 | 1.81 | 38.25 |
| COSU00795 | Malian | Rice_bran | F | 12 | 68 | 1.46 | 2.83 | 53.32 |
| COSU00796 | Malian | Rice_bran | F | 12 | 91 | 2.13 | 4.37 | 70.60 |
| COSU00801 | Malian | Rice_bran | F | 12 | 193 | 2.18 | 3.90 | 82.71 |
| COSU00802 | Malian | Rice_bran | F | 12 | 146 | 2.74 | 8.18 | 102.02 |
| COSU00804 | Malian | Rice_bran | F | 12 | 95 | 2.02 | 4.57 | 77.93 |
| COSU00805 | Malian | Rice_bran | F | 12 | 122 | 1.91 | 3.52 | 69.79 |
| COSU00806 | Malian | Rice_bran | F | 12 | 69 | 2.27 | 5.25 | 51.84 |
| COSU00786 | Malian | Rice_bran | M | 12 | 77 | 2.27 | 5.81 | 53.66 |
| COSU00787 | Malian | Rice_bran | M | 12 | 61 | 1.42 | 2.09 | 49.42 |
| COSU00788 | Malian | Rice_bran | M | 12 | 80 | 2.25 | 6.47 | 56.76 |
| COSU00794 | Malian | Rice_bran | M | 12 | 51 | 1.77 | 3.33 | 39.14 |
| COSU00797 | Malian | Rice_bran | M | 12 | 118 | 2.17 | 3.87 | 93.88 |
| COSU00798 | Malian | Rice_bran | M | 12 | 77 | 2.56 | 7.74 | 77.00 |
| COSU00799 | Malian | Rice_bran | M | 12 | 103 | 2.56 | 6.19 | 80.41 |
| COSU00800 | Malian | Rice_bran | M | 12 | 112 | 1.97 | 2.78 | 79.47 |
| COSU00803 | Malian | Rice_bran | M | 12 | 67 | 1.70 | 3.46 | 50.94 |
| COSU00807 | Malian | Rice_bran | M | 12 | 69 | 2.24 | 5.28 | 55.48 |
| COSU00808 | Malian | Rice_bran | M | 12 | 134 | 2.34 | 4.93 | 97.35 |

**Table S2.** The differences in gut microbial communities from infants fed rice bran and control. Data presented as log-fold changes in the Nicaraguan microbiome at 8 and 12 months of age for rice bran and control groups.

| **Nicaragua Rice bran vs Control at 8 months** | | | | | |
|---|---|---|---|---|---|
| **Phylum** | **Family** | **Genus** | **OTUs** | **logFC** | **adj.P.Val** |
| Actinobacteria | Bifidobacteriaceae | Bifidobacteriaceae_unclassified | Otu0314 | 2.04 | 1.4E-06 |
| | | Bifidobacteriaceae_unclassified | Otu0294 | 1.84 | 9.8E-06 |
| | | Bifidobacterium | Otu0208 | 1.61 | 2.2E-03 |
| | | Bifidobacterium | Otu0275 | 1.19 | 4.5E-03 |
| | | Bifidobacteriaceae_unclassified | Otu0183 | 1.29 | 5.8E-03 |
| | | Bifidobacteriaceae_unclassified | Otu0265 | 1.09 | 9.5E-03 |
| | | Bifidobacteriaceae_unclassified | Otu0272 | 1.23 | 1.2E-02 |
| | | Bifidobacteriaceae_unclassified | Otu0390 | 0.81 | 4.1E-02 |
| Bacteroidetes | Prevotellaceae | Prevotella_9 | Otu0121 | 3.57 | 1.4E-07 |
| | Bacteroidaceae | Bacteroides | Otu0192 | -3.08 | 2.3E-07 |
| | | Bacteroides | Otu0407 | 2.20 | 2.3E-07 |
| | | Bacteroides | Otu0321 | 2.34 | 3.2E-06 |
| | | Bacteroides | Otu0379 | 2.26 | 5.7E-06 |
| | | Bacteroides | Otu0266 | 2.89 | 9.8E-06 |
| | | Bacteroides | Otu0312 | 2.22 | 2.9E-05 |
| | | Bacteroides | Otu0433 | 2.01 | 5.0E-05 |
| | Prevotellaceae | Prevotella_9 | Otu0396 | 1.82 | 8.6E-05 |
| | Bacteroidaceae | Bacteroides | Otu0517 | 1.50 | 9.7E-05 |
| | | Bacteroides | Otu0442 | 1.60 | 2.4E-04 |
| | | Bacteroides | Otu0254 | 1.28 | 2.7E-04 |
| | | Bacteroides | Otu0414 | 1.55 | 3.5E-04 |
| | | Bacteroides | Otu0292 | 1.89 | 3.8E-04 |
| | | Bacteroides | Otu0456 | 1.43 | 4.3E-04 |
| | | Bacteroides | Otu0129 | 2.59 | 4.3E-04 |
| | | Bacteroides | Otu0319 | 1.78 | 8.2E-04 |
| | | Bacteroides | Otu0358 | 1.90 | 9.0E-04 |

| | | | | | |
|---|---|---|---|---|---|
| | | Bacteroides | Otu0334 | 1.66 | 1.0E-03 |
| | | Bacteroides | Otu0437 | 1.35 | 1.3E-03 |
| | | Bacteroides | Otu0311 | 1.72 | 2.2E-03 |
| | | Bacteroides | Otu0051 | 3.50 | 2.8E-03 |
| | | Bacteroides | Otu0502 | 1.25 | 3.1E-03 |
| | | Bacteroides | Otu0270 | 1.83 | 3.3E-03 |
| | | Bacteroides | Otu0368 | 1.51 | 4.2E-03 |
| | | Bacteroides | Otu0428 | 1.18 | 4.6E-03 |
| | | Bacteroides | Otu0410 | -1.24 | 5.1E-03 |
| | | Bacteroides | Otu0224 | 1.52 | 5.1E-03 |
| | Rikenellaceae | Alistipes | Otu0148 | 1.34 | 5.6E-03 |
| | Bacteroidaceae | Bacteroides | Otu0287 | 1.50 | 5.8E-03 |
| | | Bacteroides | Otu0169 | 1.90 | 6.1E-03 |
| | Porphyromonadaceae | Parabacteroides | Otu0086 | -2.34 | 7.4E-03 |
| | Bacteroidaceae | Bacteroides | Otu0249 | 1.16 | 7.4E-03 |
| | | Bacteroides | Otu0487 | 1.06 | 7.6E-03 |
| | Prevotellaceae | Prevotella | Otu0426 | -1.45 | 7.6E-03 |
| | Bacteroidaceae | Bacteroides | Otu0114 | 1.70 | 7.6E-03 |
| | | Bacteroides | Otu0530 | -1.06 | 9.2E-03 |
| | | Bacteroides | Otu0274 | 1.51 | 1.1E-02 |
| | Prevotellaceae | Paraprevotella | Otu0189 | 1.41 | 1.2E-02 |
| | Bacteroidaceae | Bacteroides | Otu0295 | 1.31 | 1.2E-02 |
| | | Bacteroides | Otu0405 | 1.23 | 1.4E-02 |
| | | Bacteroides | Otu0322 | 1.44 | 1.4E-02 |
| | | Bacteroides | Otu0088 | 1.82 | 1.8E-02 |
| | | Bacteroides | Otu0170 | 1.67 | 2.1E-02 |
| | | Bacteroides | Otu0279 | 1.35 | 2.3E-02 |
| | | Bacteroides | Otu0283 | 1.29 | 2.7E-02 |
| | | Bacteroides | Otu0205 | 1.32 | 4.9E-02 |
| Firmicutes | Veillonellaceae | Veillonella | Otu0245 | 3.94 | 4.8E-10 |
| | | Veillonella | Otu0289 | 4.47 | 7.6E-10 |
| | | Veillonella | Otu0241 | 3.10 | 1.4E-08 |
| | | Veillonella | Otu0253 | 4.21 | 1.4E-08 |

| | | | | |
|---|---|---|---|---|
| Erysipelotrichaceae | Candidatus_Stoquefichus | Otu0341 | 3.42 | 1.4E-08 |
| Veillonellaceae | Veillonella | Otu0316 | 3.51 | 2.2E-08 |
| Lachnospiraceae | Lachnospiraceae_unclassified | Otu0280 | 5.84 | 4.0E-08 |
| Veillonellaceae | Veillonella | Otu0347 | 2.76 | 6.1E-08 |
| | Veillonella | Otu0403 | 3.07 | 1.9E-07 |
| | Veillonella | Otu0397 | 3.04 | 1.9E-07 |
| Lachnospiraceae | Anaerostipes | Otu0284 | 2.89 | 1.9E-07 |
| Veillonellaceae | Veillonella | Otu0165 | 3.00 | 2.7E-07 |
| Lachnospiraceae | Lachnospiraceae_ND3007_group | Otu0242 | 3.10 | 2.9E-07 |
| Acidaminococcaceae | Phascolarctobacterium | Otu0066 | 4.64 | 3.3E-07 |
| Lachnospiraceae | Lachnospiraceae_unclassified | Otu0157 | 3.19 | 1.2E-06 |
| Veillonellaceae | Megasphaera | Otu0258 | 2.18 | 1.4E-06 |
| Ruminococcaceae | Faecalibacterium | Otu0514 | 1.80 | 1.8E-06 |
| Veillonellaceae | Veillonella | Otu0298 | 2.84 | 3.2E-06 |
| Clostridiaceae_1 | Clostridium_sensu_stricto_1 | Otu0099 | 3.57 | 8.5E-06 |
| Lactobacillaceae | Lactobacillus | Otu0053 | -3.85 | 1.1E-05 |
| Veillonellaceae | Megasphaera | Otu0350 | 1.59 | 2.0E-05 |
| Ruminococcaceae | Ruminococcus_2 | Otu0175 | -2.70 | 2.0E-05 |
| | Flavonifractor | Otu0553 | 1.55 | 4.0E-05 |
| | Ruminococcaceae_unclassified | Otu0374 | 1.49 | 5.2E-05 |
| Streptococcaceae | Streptococcus | Otu0201 | 1.57 | 6.1E-05 |
| Lachnospiraceae | Lachnospiraceae_unclassified | Otu0101 | 2.84 | 1.3E-04 |
| Veillonellaceae | Megasphaera | Otu0330 | 1.12 | 4.5E-04 |
| | Allisonella | Otu0209 | 1.48 | 5.3E-04 |
| Lachnospiraceae | Lachnospiraceae_unclassified | Otu0297 | -1.92 | 5.7E-04 |
| Clostridiaceae_1 | Clostridium_sensu_stricto_1 | Otu0211 | 2.06 | 8.6E-04 |
| Ruminococcaceae | Ruminococcaceae_unclassified | Otu0238 | 2.01 | 9.7E-04 |
| Erysipelotrichaceae | Holdemania | Otu0419 | 1.52 | 1.3E-03 |
| Lachnospiraceae | Anaerostipes | Otu0204 | 2.37 | 1.6E-03 |
| Clostridiaceae_1 | Clostridium_sensu_stricto_1 | Otu0191 | -2.33 | 1.6E-03 |
| Peptostreptococcaceae | Peptostreptococcaceae_unclassified | Otu0124 | -2.38 | 1.9E-03 |
| Lactobacillaceae | Lactobacillus | Otu0024 | 1.78 | 2.4E-03 |
| Ruminococcaceae | Oscillibacter | Otu0163 | -2.49 | 2.9E-03 |

| | | | | | |
|---|---|---|---|---|---|
| | Veillonellaceae | Megasphaera | Otu0421 | 1.03 | 2.9E-03 |
| | Lachnospiraceae | Lachnospiraceae_unclassified | Otu0174 | -2.27 | 3.3E-03 |
| | Ruminococcaceae | Faecalibacterium | Otu0206 | 1.48 | 3.3E-03 |
| | Lachnospiraceae | Roseburia | Otu0216 | 1.95 | 3.6E-03 |
| | | Tyzzerella_4 | Otu0435 | 1.06 | 5.1E-03 |
| | | Blautia | Otu0070 | 2.51 | 5.6E-03 |
| | Clostridiaceae_1 | Sarcina | Otu0031 | -2.89 | 6.1E-03 |
| | Erysipelotrichaceae | Erysipelatoclostridium | Otu0038 | 2.07 | 8.0E-03 |
| | Ruminococcaceae | Faecalibacterium | Otu0389 | 1.29 | 9.4E-03 |
| | Clostridiaceae_1 | Clostridium_sensu_stricto_1 | Otu0118 | 1.92 | 1.2E-02 |
| | Lachnospiraceae | Lachnospiraceae_ge | Otu0090 | -2.20 | 1.4E-02 |
| | Ruminococcaceae | Faecalibacterium | Otu0173 | 1.40 | 1.4E-02 |
| | Veillonellaceae | Veillonellaceae_unclassified | Otu0056 | 2.15 | 2.9E-02 |
| | Ruminococcaceae | Faecalibacterium | Otu0257 | 1.20 | 3.4E-02 |
| | Veillonellaceae | Megasphaera | Otu0454 | 0.96 | 4.2E-02 |
| | Family_XI | Gemella | Otu0255 | 0.76 | 4.2E-02 |
| | Ruminococcaceae | Faecalibacterium | Otu0440 | 0.69 | 4.8E-02 |
| Fusobacteria | Fusobacteriaceae | Fusobacterium | Otu0158 | 1.19 | 4.2E-02 |
| Proteobacteria | Enterobacteriaceae | Enterobacteriaceae_unclassified | Otu0357 | 2.22 | 6.8E-07 |
| | | Enterobacteriaceae_unclassified | Otu0365 | 2.03 | 3.2E-06 |
| | | Enterobacteriaceae_unclassified | Otu0348 | 1.93 | 9.8E-06 |
| | | Enterobacteriaceae_unclassified | Otu0128 | 2.20 | 3.8E-05 |
| | Desulfovibrionaceae | Desulfovibrio | Otu0164 | 2.34 | 1.7E-04 |
| | Moraxellaceae | Acinetobacter | Otu0375 | -1.20 | 2.0E-03 |
| | Enterobacteriaceae | Morganella | Otu0097 | 2.31 | 4.5E-03 |
| | Succinivibrionaceae | Succinivibrionaceae_unclassified | Otu0042 | -2.59 | 1.4E-02 |
| | Enterobacteriaceae | Proteus | Otu0113 | 1.10 | 1.4E-02 |
| | | Enterobacteriaceae_unclassified | Otu0268 | 1.37 | 2.1E-02 |
| **Nicaragua Rice bran vs Control at 12 months** | | | | | |
| **Phylum** | **Family** | **Genus** | **OTUs** | **logFC** | **adj.P.Val** |
| Actinobacteria | Bifidobacteriaceae | Bifidobacterium | Otu0332 | 2.61 | 1.4E-05 |
| | Coriobacteriaceae | Atopobium | Otu0234 | 2.14 | 3.0E-05 |
| | Bifidobacteriaceae | Bifidobacteriaceae_unclassified | Otu0314 | 1.35 | 6.5E-04 |

| | | | | | |
|---|---|---|---|---|---|
| | | Bifidobacterium | Otu0394 | 1.30 | 9.6E-03 |
| | | Bifidobacterium | Otu0275 | 1.08 | 2.5E-02 |
| Bacteroidetes | Prevotellaceae | Paraprevotella | Otu0189 | 6.21 | 4.3E-08 |
| | Bacteroidaceae | Bacteroides | Otu0146 | 5.73 | 1.4E-07 |
| | | Bacteroides | Otu0487 | 2.34 | 7.5E-06 |
| | | Bacteroides | Otu0517 | 1.90 | 2.8E-05 |
| | | Bacteroides | Otu0088 | 3.41 | 1.4E-04 |
| | | Bacteroides | Otu0287 | 2.28 | 5.0E-04 |
| | Rikenellaceae | Alistipes | Otu0148 | 1.92 | 5.3E-04 |
| | Prevotellaceae | Prevotella | Otu0426 | 2.04 | 9.3E-04 |
| | Bacteroidaceae | Bacteroides | Otu0433 | 1.77 | 1.1E-03 |
| | Rikenellaceae | Alistipes | Otu0271 | 1.74 | 1.1E-03 |
| | Prevotellaceae | Prevotella_9 | Otu0396 | 1.34 | 4.0E-03 |
| | Bacteroidaceae | Bacteroides | Otu0205 | 2.29 | 7.8E-03 |
| | | Bacteroides | Otu0456 | 1.26 | 8.0E-03 |
| | | Bacteroides | Otu0334 | 1.51 | 8.7E-03 |
| | | Bacteroides | Otu0321 | 1.30 | 1.0E-02 |
| | | Bacteroides | Otu0410 | 1.21 | 1.2E-02 |
| | | Bacteroides | Otu0292 | 1.48 | 1.9E-02 |
| | | Bacteroides | Otu0530 | 1.19 | 1.9E-02 |
| | | Bacteroides | Otu0169 | 1.96 | 2.0E-02 |
| | | Bacteroides | Otu0442 | 1.02 | 3.3E-02 |
| | | Bacteroides | Otu0224 | 1.40 | 3.8E-02 |
| Firmicutes | Veillonellaceae | Allisonella | Otu0209 | -4.07 | 1.6E-08 |
| | Ruminococcaceae | Faecalibacterium | Otu0206 | 4.38 | 1.6E-08 |
| | Acidaminococcaceae | Phascolarctobacterium | Otu0066 | 6.13 | 1.6E-08 |
| | Veillonellaceae | Veillonella | Otu0253 | 3.35 | 3.3E-07 |
| | | Megasphaera | Otu0421 | 2.49 | 3.3E-07 |
| | | Veillonella | Otu0241 | 2.42 | 1.6E-06 |
| | | Megasphaera | Otu0330 | 1.79 | 6.1E-06 |
| | | Megasphaera | Otu0438 | 2.74 | 7.1E-06 |
| | | Megasphaera | Otu0350 | 1.97 | 7.8E-06 |
| | Lachnospiraceae | Lachnospiraceae_unclassified | Otu0280 | 3.26 | 1.4E-05 |

| | | | | | |
|---|---|---|---|---|---|
| | | Eisenbergiella | Otu0156 | -2.72 | 2.4E-05 |
| | Veillonellaceae | Megasphaera | Otu0258 | 1.76 | 8.0E-05 |
| | | Veillonella | Otu0165 | 2.19 | 8.0E-05 |
| | Ruminococcaceae | Ruminococcaceae_unclassified | Otu0374 | -1.54 | 2.8E-04 |
| | Lachnospiraceae | Lachnospiraceae_ND3007_group | Otu0242 | -2.02 | 2.9E-04 |
| | | Lachnospiraceae_unclassified | Otu0353 | -1.91 | 5.6E-04 |
| | Family_XI | Gemella | Otu0255 | 1.95 | 6.5E-04 |
| | Lactobacillaceae | Lactobacillus | Otu0293 | 1.87 | 1.0E-03 |
| | Streptococcaceae | Streptococcus | Otu0201 | 1.26 | 1.5E-03 |
| | Lachnospiraceae | Tyzzerella_4 | Otu0435 | -1.39 | 3.0E-03 |
| | Veillonellaceae | Megasphaera | Otu0340 | 1.57 | 4.2E-03 |
| | | Dialister | Otu0151 | 2.35 | 4.6E-03 |
| | Ruminococcaceae | Faecalibacterium | Otu0514 | 1.06 | 4.8E-03 |
| | | Faecalibacterium | Otu0257 | 1.79 | 5.8E-03 |
| | Carnobacteriaceae | Granulicatella | Otu0364 | 1.49 | 8.7E-03 |
| | Lachnospiraceae | Lachnoclostridium | Otu0162 | 2.33 | 1.2E-02 |
| | | Sellimonas | Otu0098 | 2.13 | 1.4E-02 |
| | Lactobacillaceae | Lactobacillus | Otu0110 | 0.87 | 1.6E-02 |
| | Ruminococcaceae | Butyricicoccus | Otu0250 | 1.50 | 1.9E-02 |
| | Veillonellaceae | Megasphaera | Otu0178 | 1.25 | 1.9E-02 |
| | Clostridiaceae_1 | Clostridium_sensu_stricto_1 | Otu0259 | 1.13 | 2.2E-02 |
| | Erysipelotrichaceae | Erysipelotrichaceae_UCG-003 | Otu0116 | -2.55 | 2.2E-02 |
| | Lachnospiraceae | Dorea | Otu0085 | 1.82 | 2.5E-02 |
| | | Blautia | Otu0168 | -2.00 | 3.6E-02 |
| | Ruminococcaceae | Flavonifractor | Otu0328 | -1.33 | 3.8E-02 |
| | Lachnospiraceae | Lachnospiraceae_unclassified | Otu0108 | 2.20 | 3.8E-02 |
| | Lactobacillaceae | Lactobacillus | Otu0053 | 1.29 | 3.9E-02 |
| | Lachnospiraceae | Lachnospiraceae_unclassified | Otu0157 | -1.48 | 4.1E-02 |
| | Veillonellaceae | Megasphaera | Otu0454 | 1.17 | 4.2E-02 |
| | | Veillonellaceae_unclassified | Otu0056 | 2.29 | 4.9E-02 |
| | Lachnospiraceae | Anaerostipes | Otu0284 | -0.94 | 4.9E-02 |
| | | Lachnospiraceae_unclassified | Otu0297 | 1.07 | 4.9E-02 |

| | | | | | |
|---|---|---|---|---|---|
| Proteobacteria | Pseudomonadaceae | Pseudomonas | Otu0306 | 2.05 | 2.4E-05 |
| | Enterobacteriaceae | Proteus | Otu0113 | -2.77 | 9.1E-05 |
| | | Enterobacteriaceae_unclassified | Otu0268 | 2.09 | 4.8E-03 |
| | Campylobacteraceae | Campylobacter | Otu0064 | 1.87 | 9.7E-03 |

**Table S3.** The differences in gut microbial communities from infants fed rice bran and control. Data presented as log-fold changes in the Mali microbiome at 8 and 12 months of age for rice bran and control groups.

| **Mali Rice bran vs Control at 8 months** | | | | | |
|---|---|---|---|---|---|
| **Phylum** | **Family** | **Genus** | **OTUs** | **logFC** | **adj.P.Val** |
| Actinobacteria | Coriobacteriaceae | Coriobacteriaceae_unclassified | Otu0080 | 5.13 | 4.1E-07 |
| | | Eggerthella | Otu0142 | -2.35 | 1.1E-06 |
| | Bifidobacteriaceae | Bifidobacterium | Otu0275 | -1.83 | 3.9E-06 |
| | Corynebacteriaceae | Corynebacterium_1 | Otu0581 | -1.48 | 5.0E-03 |
| | Bifidobacteriaceae | Bifidobacteriaceae_unclassified | Otu0265 | -1.09 | 1.0E-02 |
| | Actinomycetaceae | Actinomyces | Otu0460 | -1.05 | 2.7E-02 |
| | Coriobacteriaceae | Senegalimassilia | Otu0115 | -1.55 | 2.7E-02 |
| Firmicutes | Lactobacillaceae | Lactobacillus | Otu0356 | 3.26 | 1.4E-09 |
| | Peptostreptococcaceae | Terrisporobacter | Otu0247 | 3.08 | 9.1E-08 |
| | Lactobacillaceae | Lactobacillus | Otu0120 | 5.51 | 2.0E-07 |
| | Veillonellaceae | Megasphaera | Otu0178 | -2.27 | 1.2E-03 |
| | Lactobacillaceae | Lactobacillus | Otu0016 | 4.29 | 1.2E-03 |
| | Ruminococcaceae | Ruminococcaceae_ge | Otu0075 | -2.72 | 6.1E-03 |
| | Lactobacillaceae | Lactobacillus | Otu0024 | 2.78 | 1.3E-02 |
| | Lachnospiraceae | Lachnospiraceae_unclassified | Otu0010 | -2.33 | 1.6E-02 |
| | Clostridiaceae_1 | Clostridium_sensu_stricto_1 | Otu0099 | -1.15 | 4.7E-02 |
| | Family_XI | Gemella | Otu0255 | -0.91 | 4.7E-02 |
| Proteobacteria | Campylobacteraceae | Campylobacter | Otu0123 | -2.22 | 3.1E-02 |
| Verrucomicrobia | Verrucomicrobiaceae | Akkermansia | Otu0018 | 2.25 | 6.1E-03 |
| **Mali Rice bran vs Control at 12 months** | | | | | |
| **Phylum** | **Family** | **Genus** | **OTUs** | **logFC** | **adj.P.Val** |
| Actinobacteria | Dietziaceae | Dietzia | Otu0550 | -1.73 | 4.0E-05 |
| | Bifidobacteriaceae | Bifidobacteriaceae_unclassified | Otu0265 | -2.64 | 4.0E-05 |
| | | Bifidobacteriaceae_unclassified | Otu0278 | 1.54 | 3.2E-04 |

| | | | | | |
|---|---|---|---|---|---|
| | Micrococcaceae | Kocuria | Otu0325 | 1.61 | 2.3E-03 |
| | Bifidobacteriaceae | Bifidobacterium | Otu0275 | -0.88 | 2.2E-02 |
| | | Bifidobacteriaceae_unclassified | Otu0390 | -0.57 | 4.6E-02 |
| Bacteroidetes | Prevotellaceae | Alloprevotella | Otu0068 | 3.63 | 3.5E-04 |
| Firmicutes | Peptostreptococcaceae | Terrisporobacter | Otu0247 | -2.18 | 7.0E-06 |
| | Lachnospiraceae | Lachnospira | Otu0058 | -2.83 | 2.9E-05 |
| | Clostridiaceae_1 | Clostridium_sensu_stricto_1 | Otu0076 | -3.64 | 4.0E-05 |
| | Lachnospiraceae | Tyzzerella_4 | Otu0026 | -3.29 | 6.5E-05 |
| | Veillonellaceae | Megasphaera | Otu0361 | -3.55 | 3.2E-04 |
| | Ruminococcaceae | Faecalibacterium | Otu0173 | -1.63 | 2.1E-03 |
| | Lachnospiraceae | Lachnospiraceae_unclassified | Otu0101 | -1.66 | 7.0E-03 |
| | Lactobacillaceae | Lactobacillus | Otu0053 | 2.70 | 9.8E-03 |
| | Lachnospiraceae | Roseburia | Otu0047 | -1.48 | 1.6E-02 |
| | Veillonellaceae | Veillonella | Otu0152 | 1.99 | 1.6E-02 |
| | Peptostreptococcaceae | Peptostreptococcaceae_unclassified | Otu0078 | -1.51 | 1.6E-02 |
| | Streptococcaceae | Streptococcus | Otu0479 | 1.05 | 1.9E-02 |
| | Lactobacillaceae | Lactobacillus | Otu0024 | 2.18 | 2.0E-02 |
| | Peptococcaceae | Peptococcus | Otu0351 | -1.63 | 2.3E-02 |
| | Ruminococcaceae | Faecalibacterium | Otu0257 | -0.77 | 2.7E-02 |
| Fusobacteria | Fusobacteriaceae | Fusobacterium | Otu0158 | -1.81 | 8.4E-03 |
| Proteobacteria | Enterobacteriaceae | Enterobacteriaceae_unclassified | Otu0304 | -1.61 | 4.0E-05 |
| | | Enterobacteriaceae_unclassified | Otu0128 | -1.62 | 1.2E-03 |
| | Campylobacteraceae | Campylobacter | Otu0123 | 2.55 | 9.1E-03 |
| | Neisseriaceae | Neisseria | Otu0318 | 1.42 | 3.1E-02 |
| | Pasteurellaceae | Haemophilus | Otu0180 | 1.46 | 3.2E-02 |
| Verrucomicrobia | Verrucomicrobiaceae | Akkermansia | Otu0018 | -1.80 | 1.3E-02 |

**Table S4.** The OTUs with overlapping significance for response to rice bran when compared to control in both Mali and Nicaragua**.**

| | | | | mali8m | | mali12m | | nic8m | | nic12m | |
|---|---|---|---|---|---|---|---|---|---|---|---|
| **Phylum** | **Family** | **Genus** | **OTUs** | **logFC** | **adj.P.Val** | **logFC** | **adj.P.Val** | **logFC** | **adj.P.Val** | **logFC** | **adj.P.Val** |
| Actinobacteria | Bifidobacteriaceae | Bifidobacterium | Otu0275 | -1.83 | 3.9E-06 | -0.88 | 0.02 | 1.19 | 0.004 | 1.08 | 0.025 |
| Firmicutes | Veillonellaceae | Megasphaera | Otu0178 | -2.27 | 1.2E-03 | | | | | 1.25 | 0.019 |
| Actinobacteria | Bifidobacteriaceae | Bifidobacteriaceae_unclassified | Otu0265 | -1.09 | 1.0E-02 | -1.09 | 0.01 | 1.09 | 0.009 | | |
| Firmicutes | Family_XI | Gemella | Otu0255 | -0.91 | 4.7E-02 | | | 0.76 | 0.042 | 1.95 | 0.001 |
| Firmicutes | Lactobacillaceae | Lactobacillus | Otu0024 | 2.78 | 1.3E-02 | 2.18 | 0.02 | 1.78 | 0.002 | | |
| Proteobacteria | Enterobacteriaceae | Enterobacteriaceae_unclassified | Otu0128 | | | -1.62 | 0.00 | 2.20 | 0.000 | | |
| Firmicutes | Ruminococcaceae | Faecalibacterium | Otu0173 | | | -1.63 | 0.00 | 1.40 | 0.014 | | |
| Firmicutes | Lachnospiraceae | Lachnospiraceae_unclassified | Otu0101 | | | -1.66 | 0.01 | 2.84 | 0.000 | | |
| Fusobacteria | Fusobacteriaceae | Fusobacterium | Otu0158 | | | -1.81 | 0.01 | 1.19 | 0.042 | | |
| Firmicutes | Lactobacillaceae | Lactobacillus | Otu0053 | | | 2.70 | 0.01 | -3.85 | 0.000 | 1.29 | 0.039 |
| Firmicutes | Ruminococcaceae | Faecalibacterium | Otu0257 | | | -0.77 | 0.03 | 1.20 | 0.034 | | |
| Actinobacteria | Bifidobacteriaceae | Bifidobacteriaceae_unclassified | Otu0390 | | | -0.57 | 0.05 | 0.81 | 0.041 | | |

**Table S5.** All of the stool metabolites identified from the stool metabolome, including those with unknown identity that had fold differences in the relative abundances between rice bran supplementation and control infants. Data are presented for both Nicaraguan and Malian infants at 8 months of age.

| Chemical Class | Metabolic Pathways | Metabolites | HMDB | Nicaragua | | Mali | |
|---|---|---|---|---|---|---|---|
| | | | | Fold Differences | p-value | Fold Differences | p-value |
| Amino Acid | Glycine, Serine and Threonine Metabolism | glycine | HMDB00123 | **0.66** | 0.01 | 1.2 | 0.24 |
| Amino Acid | Glycine, Serine and Threonine Metabolism | N-acetylglycine | HMDB00532 | 0.73 | 0.18 | 1.13 | 0.60 |
| Amino Acid | Glycine, Serine and Threonine Metabolism | sarcosine | HMDB00271 | 0.99 | 0.98 | 0.69 | 0.45 |
| Amino Acid | Glycine, Serine and Threonine Metabolism | dimethylglycine | HMDB00092 | 0.69 | 0.29 | **0.42** | 0.02 |
| Amino Acid | Glycine, Serine and Threonine Metabolism | betaine | HMDB00043 | 0.67 | 0.38 | 0.74 | 0.53 |
| Amino Acid | Glycine, Serine and Threonine Metabolism | serine | HMDB00187 | 0.87 | 0.48 | 1.1 | 0.63 |
| Amino Acid | Glycine, Serine and Threonine Metabolism | N-acetylserine | HMDB02931 | 0.89 | 0.59 | 0.95 | 0.83 |
| Amino Acid | Glycine, Serine and Threonine Metabolism | O-acetylserine | HMDB03011 | 1.05 | 0.34 | **0.89** | 0.06 |
| Amino Acid | Glycine, Serine and Threonine Metabolism | 2-methylserine | | 0.67 | 0.32 | 1.24 | 0.61 |
| Amino Acid | Glycine, Serine and Threonine Metabolism | threonine | HMDB00167 | 0.8 | 0.32 | 1.11 | 0.64 |
| Amino Acid | Glycine, Serine and Threonine Metabolism | N-acetylthreonine | | 0.9 | 0.80 | 0.72 | 0.46 |
| Amino Acid | Glycine, Serine and Threonine Metabolism | allo-threonine | HMDB04041 | 1.13 | 0.77 | 0.86 | 0.73 |
| Amino Acid | Glycine, Serine and Threonine Metabolism | homoserine | HMDB00719 | 0.77 | 0.55 | 0.64 | 0.34 |
| Amino Acid | Glycine, Serine and Threonine | O-acetylhomoserine | | 0.65 | 0.46 | 1.1 | 0.88 |

| | | | | | | | |
|---|---|---|---|---|---|---|---|
| | Metabolism | | | | | | |
| Amino Acid | Alanine and Aspartate Metabolism | alanine | HMDB00161 | 0.84 | 0.12 | 1.08 | 0.49 |
| Amino Acid | Alanine and Aspartate Metabolism | N-acetylalanine | HMDB00766 | 0.88 | 0.49 | 1.07 | 0.75 |
| Amino Acid | Alanine and Aspartate Metabolism | N-methylalanine | HMDB01906 | 1.13 | 0.77 | 0.66 | 0.34 |
| Amino Acid | Alanine and Aspartate Metabolism | N-propionylalanine | | 1.25 | 0.62 | 0.6 | 0.28 |
| Amino Acid | Alanine and Aspartate Metabolism | aspartate | HMDB00191 | 1.18 | 0.53 | **1.66** | 0.07 |
| Amino Acid | Alanine and Aspartate Metabolism | N-acetylaspartate (NAA) | HMDB00812 | 0.95 | 0.90 | 0.96 | 0.92 |
| Amino Acid | Alanine and Aspartate Metabolism | asparagine | HMDB00168 | 1.13 | 0.78 | **2.16** | 0.10 |
| Amino Acid | Alanine and Aspartate Metabolism | N-acetylasparagine | HMDB06028 | 0.92 | 0.72 | 1.05 | 0.84 |
| Amino Acid | Glutamate Metabolism | glutamate | HMDB00148 | 0.88 | 0.60 | 1.46 | 0.15 |
| Amino Acid | Glutamate Metabolism | glutamine | HMDB00641 | 0.72 | 0.52 | 2.19 | 0.13 |
| Amino Acid | Glutamate Metabolism | N-acetylglutamate | HMDB01138 | 0.95 | 0.85 | 1.44 | 0.21 |
| Amino Acid | Glutamate Metabolism | N-acetylglutamine | HMDB06029 | 0.85 | 0.48 | 0.72 | 0.17 |
| Amino Acid | Glutamate Metabolism | 4-hydroxyglutamate | HMDB01344 | 0.8 | 0.57 | 1.04 | 0.91 |
| Amino Acid | Glutamate Metabolism | gamma-carboxyglutamate | HMDB41900 | **1.97** | 0.08 | 0.81 | 0.61 |
| Amino Acid | Glutamate Metabolism | glutamate, gamma-methyl ester | HMDB61715 | 0.99 | 0.98 | 1.3 | 0.29 |
| Amino Acid | Glutamate Metabolism | pyroglutamine* | | 0.78 | 0.46 | 0.92 | 0.82 |
| Amino Acid | Glutamate Metabolism | beta-citrylglutamate | | 0.95 | 0.92 | 0.49 | 0.18 |
| Amino Acid | Glutamate Metabolism | gamma-aminobutyrate (GABA) | HMDB00112 | 0.78 | 0.61 | 1.61 | 0.35 |
| Amino Acid | Glutamate Metabolism | carboxyethyl-GABA | HMDB02201 | 0.8 | 0.47 | 1.3 | 0.40 |
| Amino Acid | Glutamate Metabolism | S-1-pyrroline-5-carboxylate | HMDB01301 | 0.84 | 0.31 | 0.98 | 0.92 |
| Amino Acid | Glutamate Metabolism | propionylglutamine | | 1.18 | 0.58 | 0.75 | 0.35 |
| Amino Acid | Glutamate Metabolism | butyrylglutamine/isobutyrylglutamine | | 1.12 | 0.79 | 0.9 | 0.81 |
| Amino Acid | Glutamate Metabolism | succinylglutamine | | 0.86 | 0.65 | 0.82 | 0.59 |
| Amino Acid | Glutamate Metabolism | valerylglutamine | | 0.75 | 0.54 | 0.96 | 0.93 |
| Amino Acid | Histidine Metabolism | histidine | HMDB00177 | 0.8 | 0.45 | 0.68 | 0.20 |
| Amino Acid | Histidine Metabolism | 1-methylhistidine | HMDB00001 | 1.19 | 0.60 | 0.79 | 0.49 |
| Amino Acid | Histidine Metabolism | 3-methylhistidine | HMDB00479 | 0.64 | 0.25 | **0.48** | 0.07 |
| Amino Acid | Histidine Metabolism | N-acetylhistidine | HMDB32055 | 1.28 | 0.40 | 1.27 | 0.45 |
| Amino Acid | Histidine Metabolism | N-acetyl-1-methylhistidine* | | 0.87 | 0.74 | 0.89 | 0.79 |

| | | | | | | | |
|---|---|---|---|---|---|---|---|
| Amino Acid | Histidine Metabolism | hydantoin-5-propionic acid | HMDB01212 | 0.98 | 0.94 | 1.19 | 0.62 |
| Amino Acid | Histidine Metabolism | trans-urocanate | HMDB00301 | 0.86 | 0.77 | 1.79 | 0.30 |
| Amino Acid | Histidine Metabolism | cis-urocanate | HMDB34174 | 0.73 | 0.28 | 0.91 | 0.75 |
| Amino Acid | Histidine Metabolism | imidazole propionate | HMDB02271 | 0.72 | 0.47 | 1.07 | 0.89 |
| Amino Acid | Histidine Metabolism | formiminoglutamate | HMDB00854 | 0.83 | 0.74 | 1.74 | 0.32 |
| Amino Acid | Histidine Metabolism | imidazole lactate | HMDB02320 | 1.07 | 0.89 | 1.18 | 0.74 |
| Amino Acid | Histidine Metabolism | carnosine | HMDB00033 | **0.4** | 0.08 | 0.72 | 0.55 |
| Amino Acid | Histidine Metabolism | homocarnosine | HMDB00745 | 0.82 | 0.63 | 0.78 | 0.55 |
| Amino Acid | Histidine Metabolism | N-acetylcarnosine | HMDB12881 | 0.75 | 0.44 | 0.87 | 0.73 |
| Amino Acid | Histidine Metabolism | anserine | HMDB00194 | 0.57 | 0.23 | 0.83 | 0.71 |
| Amino Acid | Histidine Metabolism | histamine | HMDB00870 | 1.05 | 0.94 | 2.16 | 0.25 |
| Amino Acid | Histidine Metabolism | 1-methylhistamine | HMDB00898 | 0.94 | 0.89 | 1.13 | 0.77 |
| Amino Acid | Histidine Metabolism | 1-methyl-4-imidazoleacetate | HMDB02820 | 0.69 | 0.23 | 0.79 | 0.47 |
| Amino Acid | Histidine Metabolism | 4-imidazoleacetate | HMDB02024 | 1.04 | 0.92 | 1.4 | 0.40 |
| Amino Acid | Histidine Metabolism | N-acetylhistamine | HMDB13253 | 0.86 | 0.73 | 1.61 | 0.30 |
| Amino Acid | Lysine Metabolism | lysine | HMDB00182 | 0.8 | 0.21 | 1.29 | 0.17 |
| Amino Acid | Lysine Metabolism | N2-acetyllysine | HMDB00446 | 0.92 | 0.79 | 1.08 | 0.81 |
| Amino Acid | Lysine Metabolism | N6-acetyllysine | HMDB00206 | 0.73 | 0.10 | 0.93 | 0.71 |
| Amino Acid | Lysine Metabolism | N2,N6-diacetyllysine | | 1.09 | 0.68 | 0.8 | 0.33 |
| Amino Acid | Lysine Metabolism | N6-formyllysine | | 0.6 | 0.19 | **2.37** | 0.04 |
| Amino Acid | Lysine Metabolism | N6-carboxyethyllysine | | 0.6 | 0.18 | 1.71 | 0.17 |
| Amino Acid | Lysine Metabolism | N6,N6,N6-trimethyllysine | HMDB01325 | **0.56** | 0.08 | 1.03 | 0.94 |
| Amino Acid | Lysine Metabolism | 5-hydroxylysine | HMDB00450 | 0.91 | 0.78 | 1.28 | 0.52 |
| Amino Acid | Lysine Metabolism | saccharopine | HMDB00279 | 1.05 | 0.92 | 1.36 | 0.50 |
| Amino Acid | Lysine Metabolism | 2-aminoadipate | HMDB00510 | 0.73 | 0.29 | 0.84 | 0.56 |
| Amino Acid | Lysine Metabolism | pipecolate | HMDB00070 | 0.92 | 0.81 | 1.1 | 0.79 |
| Amino Acid | Lysine Metabolism | 6-oxopiperidine-2-carboxylate | HMDB61705 | 0.82 | 0.32 | 1.09 | 0.67 |
| Amino Acid | Lysine Metabolism | cadaverine | HMDB02322 | 0.86 | 0.70 | 0.75 | 0.48 |
| Amino Acid | Lysine Metabolism | N-acetyl-cadaverine | HMDB02284 | 0.76 | 0.50 | 0.78 | 0.55 |
| Amino Acid | Lysine Metabolism | 5-aminovalerate | HMDB03355 | 0.74 | 0.35 | 0.69 | 0.28 |
| Amino Acid | Lysine Metabolism | N-trimethyl 5-aminovalerate | | 0.88 | 0.78 | 1.02 | 0.96 |
| Amino Acid | Phenylalanine Metabolism | phenylalanine | HMDB00159 | 0.88 | 0.27 | 1.13 | 0.28 |
| Amino Acid | Phenylalanine Metabolism | N-acetylphenylalanine | HMDB00512 | 1.06 | 0.85 | 1.11 | 0.76 |
| Amino Acid | Phenylalanine Metabolism | phenylpyruvate | HMDB00205 | **0.59** | 0.03 | 1.18 | 0.53 |
| Amino Acid | Phenylalanine Metabolism | phenyllactate (PLA) | HMDB00779 | **0.44** | 0.02 | 1.49 | 0.29 |
| Amino Acid | Phenylalanine | phenethylamine | HMDB02017 | 1.04 | 0.94 | 0.66 | 0.42 |

| | | | | | | | |
|---|---|---|---|---|---|---|---|
| | Metabolism | | | | | | |
| Amino Acid | Phenylalanine Metabolism | phenylacetate | HMDB00209 | 1.47 | 0.48 | 1.11 | 0.85 |
| Amino Acid | Phenylalanine Metabolism | 4-hydroxyphenylacetate | HMDB00020 | 0.95 | 0.93 | 1.65 | 0.37 |
| Amino Acid | Phenylalanine Metabolism | 3-hydroxyphenylacetate | HMDB00440 | 0.87 | 0.65 | 1 | 0.99 |
| Amino Acid | Phenylalanine Metabolism | valerylphenylalanine | | 1.15 | 0.84 | 1.52 | 0.58 |
| Amino Acid | Tyrosine Metabolism | tyrosine | HMDB00158 | 0.94 | 0.75 | 1.36 | 0.15 |
| Amino Acid | Tyrosine Metabolism | N-acetyltyrosine | HMDB00866 | 0.88 | 0.62 | 1.55 | 0.12 |
| Amino Acid | Tyrosine Metabolism | tyramine | HMDB00306 | 0.6 | 0.36 | 1.22 | 0.73 |
| Amino Acid | Tyrosine Metabolism | 4-hydroxyphenylpyruvate | HMDB00707 | **0.55** | 0.02 | 0.92 | 0.75 |
| Amino Acid | Tyrosine Metabolism | 3-(4-hydroxyphenyl)lactate | HMDB00755 | 0.55 | 0.17 | 2.06 | 0.11 |
| Amino Acid | Tyrosine Metabolism | phenol sulfate | HMDB60015 | 0.66 | 0.47 | 1.38 | 0.59 |
| Amino Acid | Tyrosine Metabolism | dopamine | HMDB00073 | 0.63 | 0.34 | 1.21 | 0.70 |
| Amino Acid | Tyrosine Metabolism | vanillactate | HMDB00913 | 0.67 | 0.27 | 1.31 | 0.48 |
| Amino Acid | Tyrosine Metabolism | vanillylmandelate (VMA) | HMDB00291 | 0.71 | 0.38 | 1.38 | 0.43 |
| Amino Acid | Tyrosine Metabolism | 3-methoxytyrosine | HMDB01434 | 0.85 | 0.60 | 1.08 | 0.80 |
| Amino Acid | Tyrosine Metabolism | 3-methoxytyramine sulfate | | 1.04 | 0.79 | 1.14 | 0.42 |
| Amino Acid | Tyrosine Metabolism | (R)-salsolinol | HMDB05199 | 0.75 | 0.52 | 1.39 | 0.49 |
| Amino Acid | Tyrosine Metabolism | o-Tyrosine | HMDB06050 | 0.71 | 0.21 | 1.32 | 0.33 |
| Amino Acid | Tyrosine Metabolism | gentisate | HMDB00152 | 0.91 | 0.83 | 1.81 | 0.23 |
| Amino Acid | Tyrosine Metabolism | 2-hydroxyphenylacetate | HMDB00669 | 1.12 | 0.80 | 0.64 | 0.33 |
| Amino Acid | Tyrosine Metabolism | dopamine 3-O-sulfate | HMDB06275 | 0.92 | 0.87 | 1.59 | 0.37 |
| Amino Acid | Tyrosine Metabolism | tyramine O-sulfate | HMDB06409 | 1.08 | 0.90 | 0.4 | 0.15 |
| Amino Acid | Tyrosine Metabolism | N-formylphenylalanine | | 0.83 | 0.41 | 1.47 | 0.12 |
| Amino Acid | Tyrosine Metabolism | vanillic alcohol sulfate | | 0.7 | 0.26 | **2.01** | 0.04 |
| Amino Acid | Tryptophan Metabolism | tryptophan | HMDB00929 | 0.78 | 0.14 | 1 | 1.00 |
| Amino Acid | Tryptophan Metabolism | N-acetyltryptophan | HMDB13713 | 1.02 | 0.94 | 1.01 | 0.96 |
| Amino Acid | Tryptophan Metabolism | C-glycosyltryptophan | | 0.78 | 0.58 | 0.95 | 0.90 |
| Amino Acid | Tryptophan Metabolism | tryptophan betaine | HMDB61115 | 1.24 | 0.57 | 1.39 | 0.39 |
| Amino Acid | Tryptophan Metabolism | kynurenine | HMDB00684 | 0.84 | 0.39 | 0.91 | 0.68 |
| Amino Acid | Tryptophan Metabolism | N-acetylkynurenine (2) | | 1.05 | 0.80 | 0.98 | 0.91 |
| Amino Acid | Tryptophan Metabolism | kynurenate | HMDB00715 | **0.48** | 0.01 | 1.13 | 0.66 |
| Amino Acid | Tryptophan Metabolism | N-formylanthranilic acid | HMDB04089 | 1.2 | 0.54 | **0.51** | 0.03 |
| Amino Acid | Tryptophan Metabolism | xanthurenate | HMDB00881 | **0.47** | 0.07 | 1.6 | 0.28 |
| Amino Acid | Tryptophan Metabolism | picolinate | HMDB02243 | 0.96 | 0.86 | 0.93 | 0.78 |

| | | | | | | | |
|---|---|---|---|---|---|---|---|
| Amino Acid | Tryptophan Metabolism | serotonin | HMDB00259 | 0.78 | 0.29 | 1.05 | 0.84 |
| Amino Acid | Tryptophan Metabolism | tryptamine | HMDB00303 | 1.46 | 0.57 | 2.4 | 0.21 |
| Amino Acid | Tryptophan Metabolism | indolepyruvate | HMDB60484 | 0.97 | 0.66 | **1.13** | 0.09 |
| Amino Acid | Tryptophan Metabolism | indolelactate | HMDB00671 | 0.51 | 0.21 | 1.5 | 0.46 |
| Amino Acid | Tryptophan Metabolism | indoleacetate | HMDB00197 | 0.7 | 0.31 | 1.28 | 0.50 |
| Amino Acid | Tryptophan Metabolism | indolepropionate | HMDB02302 | **4.67** | 0.02 | 1.33 | 0.67 |
| Amino Acid | Tryptophan Metabolism | indolepropionylglycine | | 0.98 | 0.93 | 1.13 | 0.62 |
| Amino Acid | Tryptophan Metabolism | indoleacetylglutamine | HMDB13240 | 1.03 | 0.96 | 0.78 | 0.61 |
| Amino Acid | Tryptophan Metabolism | skatol | HMDB00466 | 0.84 | 0.28 | 1.03 | 0.86 |
| Amino Acid | Tryptophan Metabolism | indole | HMDB00738 | 1.17 | 0.69 | 1.08 | 0.86 |
| Amino Acid | Tryptophan Metabolism | indole-3-carboxylic acid | HMDB03320 | 0.71 | 0.39 | **0.5** | 0.09 |
| Amino Acid | Tryptophan Metabolism | 3-indoxyl sulfate | HMDB00682 | 0.8 | 0.74 | 0.57 | 0.45 |
| Amino Acid | Tryptophan Metabolism | 5-bromotryptophan | | 1 | 1.00 | 0.94 | 0.50 |
| Amino Acid | Leucine, Isoleucine and Valine Metabolism | leucine | HMDB00687 | 0.9 | 0.36 | 1.11 | 0.37 |
| Amino Acid | Leucine, Isoleucine and Valine Metabolism | N-acetylleucine | HMDB11756 | 0.86 | 0.65 | 1.05 | 0.89 |
| Amino Acid | Leucine, Isoleucine and Valine Metabolism | 4-methyl-2-oxopentanoate | HMDB00695 | **0.63** | 0.09 | 1.22 | 0.47 |
| Amino Acid | Leucine, Isoleucine and Valine Metabolism | alpha-hydroxyisocaproate | HMDB00746 | **0.45** | 0.03 | 1.05 | 0.89 |
| Amino Acid | Leucine, Isoleucine and Valine Metabolism | isovalerate (i5:0) | HMDB00718 | 1.33 | 0.60 | 1.45 | 0.51 |
| Amino Acid | Leucine, Isoleucine and Valine Metabolism | isovalerylglycine | HMDB00678 | 0.85 | 0.63 | 0.99 | 0.97 |
| Amino Acid | Leucine, Isoleucine and Valine Metabolism | isovalerylcarnitine (C5) | HMDB00688 | 1.05 | 0.84 | 1.14 | 0.61 |
| Amino Acid | Leucine, Isoleucine and Valine Metabolism | isovalerylglutamine | | 0.62 | 0.15 | 1.09 | 0.81 |
| Amino Acid | Leucine, Isoleucine and Valine Metabolism | isovalerylhistidine | | 1.01 | 0.98 | 1.16 | 0.76 |
| Amino Acid | Leucine, Isoleucine and Valine Metabolism | isovaleryltryptophan | | 0.85 | 0.72 | 1.45 | 0.44 |
| Amino Acid | Leucine, Isoleucine and Valine Metabolism | beta-hydroxyisovalerate | HMDB00754 | 0.91 | 0.80 | 1.01 | 0.98 |
| Amino Acid | Leucine, Isoleucine and Valine Metabolism | 3-methylglutaconate | HMDB00522 | 0.85 | 0.46 | 0.87 | 0.55 |
| Amino Acid | Leucine, Isoleucine and Valine Metabolism | 5-methylnorleucine | | 0.86 | 0.36 | 1.24 | 0.21 |
| Amino Acid | Leucine, Isoleucine and Valine Metabolism | isoleucine | HMDB00172 | 0.88 | 0.29 | 1.05 | 0.72 |

| | | | | | | | |
|---|---|---|---|---|---|---|---|
| Amino Acid | Leucine, Isoleucine and Valine Metabolism | allo-isoleucine | | 0.67 | 0.27 | 1.43 | 0.34 |
| Amino Acid | Leucine, Isoleucine and Valine Metabolism | N-acetylisoleucine | HMDB61684 | 0.78 | 0.44 | 1.07 | 0.83 |
| Amino Acid | Leucine, Isoleucine and Valine Metabolism | 3-methyl-2-oxovalerate | HMDB03736 | 0.64 | 0.12 | 1.29 | 0.39 |
| Amino Acid | Leucine, Isoleucine and Valine Metabolism | alpha-hydroxyisovalerate | HMDB00407 | **0.47** | 0.03 | 1.08 | 0.83 |
| Amino Acid | Leucine, Isoleucine and Valine Metabolism | 2-methylbutyrylcarnitine (C5) | HMDB00378 | 0.8 | 0.55 | 1.4 | 0.39 |
| Amino Acid | Leucine, Isoleucine and Valine Metabolism | 2-methylbutyrylglycine | HMDB00339 | 0.81 | 0.41 | 1.1 | 0.72 |
| Amino Acid | Leucine, Isoleucine and Valine Metabolism | ethylmalonate | HMDB00622 | 0.77 | 0.37 | 0.74 | 0.34 |
| Amino Acid | Leucine, Isoleucine and Valine Metabolism | methylsuccinate | HMDB01844 | 0.72 | 0.34 | 1.08 | 0.83 |
| Amino Acid | Leucine, Isoleucine and Valine Metabolism | 2,3-dimethylsuccinate | | 1.08 | 0.86 | 1.17 | 0.70 |
| Amino Acid | Leucine, Isoleucine and Valine Metabolism | valine | HMDB00883 | **0.74** | 0.10 | 1.17 | 0.39 |
| Amino Acid | Leucine, Isoleucine and Valine Metabolism | N-acetylvaline | HMDB11757 | 0.68 | 0.20 | 1.19 | 0.58 |
| Amino Acid | Leucine, Isoleucine and Valine Metabolism | 3-methyl-2-oxobutyrate | HMDB00019 | **0.52** | 0.03 | 1.4 | 0.28 |
| Amino Acid | Leucine, Isoleucine and Valine Metabolism | 2-hydroxy-3-methylvalerate | HMDB00317 | **0.46** | 0.02 | 1.14 | 0.71 |
| Amino Acid | Leucine, Isoleucine and Valine Metabolism | isobutyrylcarnitine (C4) | HMDB00736 | 0.91 | 0.81 | **2.08** | 0.07 |
| Amino Acid | Leucine, Isoleucine and Valine Metabolism | isobutyrylglycine | HMDB00730 | 0.67 | 0.24 | 0.62 | 0.19 |
| Amino Acid | Leucine, Isoleucine and Valine Metabolism | 3-hydroxyisobutyrate | HMDB00336 | 1.96 | 0.13 | 1.87 | 0.18 |
| Amino Acid | Leucine, Isoleucine and Valine Metabolism | 2,3-dihydroxy-2-methylbutyrate | HMDB29576 | 2.71 | 0.21 | 1.25 | 0.79 |
| Amino Acid | Methionine, Cysteine, SAM and Taurine Metabolism | methionine | HMDB00696 | 0.79 | 0.47 | **1.96** | 0.05 |
| Amino Acid | Methionine, Cysteine, SAM and Taurine Metabolism | N-acetylmethionine | HMDB11745 | 0.89 | 0.82 | **3.24** | 0.02 |
| Amino Acid | Methionine, Cysteine, SAM and Taurine Metabolism | N-formylmethionine | HMDB01015 | 0.56 | 0.20 | **2.84** | 0.03 |
| Amino Acid | Methionine, Cysteine, SAM and Taurine Metabolism | methionine sulfone | | 0.64 | 0.12 | 1.09 | 0.78 |
| Amino Acid | Methionine, Cysteine, SAM and Taurine Metabolism | methionine sulfoxide | HMDB02005 | 1.04 | 0.82 | 0.82 | 0.28 |
| Amino Acid | Methionine, Cysteine, SAM and Taurine Metabolism | N-acetylmethionine sulfoxide | | 1.07 | 0.78 | 0.86 | 0.55 |

| | | | | | | | |
|---|---|---|---|---|---|---|---|
| Amino Acid | Methionine, Cysteine, SAM and Taurine Metabolism | 4-methylthio-2-oxobutanoate | HMDB01553 | 0.61 | 0.11 | 1.23 | 0.52 |
| Amino Acid | Methionine, Cysteine, SAM and Taurine Metabolism | alpha-ketobutyrate | HMDB00005 | **0.53** | 0.07 | 0.9 | 0.78 |
| Amino Acid | Methionine, Cysteine, SAM and Taurine Metabolism | cysteine | HMDB00574 | **0.66** | 0.05 | 1.29 | 0.24 |
| Amino Acid | Methionine, Cysteine, SAM and Taurine Metabolism | N-acetylcysteine | HMDB01890 | 0.76 | 0.28 | 1.21 | 0.47 |
| Amino Acid | Methionine, Cysteine, SAM and Taurine Metabolism | S-methylcysteine | HMDB02108 | **0.49** | 0.05 | 1.39 | 0.38 |
| Amino Acid | Methionine, Cysteine, SAM and Taurine Metabolism | cysteine s-sulfate | HMDB00731 | **0.61** | 0.08 | 1.18 | 0.57 |
| Amino Acid | Methionine, Cysteine, SAM and Taurine Metabolism | cystine | HMDB00192 | 1.06 | 0.84 | 1.33 | 0.37 |
| Amino Acid | Methionine, Cysteine, SAM and Taurine Metabolism | cysteine sulfinic acid | HMDB00996 | 0.68 | 0.15 | 1.28 | 0.37 |
| Amino Acid | Methionine, Cysteine, SAM and Taurine Metabolism | hypotaurine | HMDB00965 | **0.52** | 0.04 | 0.9 | 0.75 |
| Amino Acid | Methionine, Cysteine, SAM and Taurine Metabolism | taurine | HMDB00251 | 0.57 | 0.18 | 1.16 | 0.74 |
| Amino Acid | Methionine, Cysteine, SAM and Taurine Metabolism | N-acetyltaurine | | 0.62 | 0.23 | 0.92 | 0.83 |
| Amino Acid | Methionine, Cysteine, SAM and Taurine Metabolism | 3-sulfo-L-alanine | HMDB02757 | 1.06 | 0.81 | 1.05 | 0.83 |
| Amino Acid | Urea cycle; Arginine and Proline Metabolism | arginine | HMDB00517 | 0.92 | 0.79 | 1.22 | 0.58 |
| Amino Acid | Urea cycle; Arginine and Proline Metabolism | argininosuccinate | HMDB00052 | 1.19 | 0.52 | 0.92 | 0.76 |
| Amino Acid | Urea cycle; Arginine and Proline Metabolism | ornithine | HMDB03374 | 0.82 | 0.48 | 1.13 | 0.68 |
| Amino Acid | Urea cycle; Arginine and Proline Metabolism | norvaline | HMDB13716 | 0.84 | 0.78 | 0.65 | 0.53 |
| Amino Acid | Urea cycle; Arginine and Proline Metabolism | 2-oxoarginine* | HMDB04225 | 0.65 | 0.35 | 0.95 | 0.91 |
| Amino Acid | Urea cycle; Arginine and Proline Metabolism | citrulline | HMDB00904 | 0.88 | 0.65 | 0.88 | 0.67 |
| Amino Acid | Urea cycle; Arginine and Proline Metabolism | homoarginine | HMDB00670 | 0.74 | 0.47 | 1.53 | 0.32 |
| Amino Acid | Urea cycle; Arginine and Proline Metabolism | homocitrulline | HMDB00679 | 0.84 | 0.64 | 0.97 | 0.94 |
| Amino Acid | Urea cycle; Arginine and Proline Metabolism | proline | HMDB00162 | **0.72** | 0.08 | 1.23 | 0.30 |
| Amino Acid | Urea cycle; Arginine and Proline Metabolism | dimethylarginine (SDMA + ADMA) | HMDB01539 | **0.53** | 0.01 | 0.92 | 0.74 |
| Amino Acid | Urea cycle; Arginine and Proline Metabolism | N-acetylarginine | HMDB04620 | 1.19 | 0.62 | 0.65 | 0.25 |

| | | | | | | | |
|---|---|---|---|---|---|---|---|
| Amino Acid | Urea cycle; Arginine and Proline Metabolism | N-acetylcitrulline | HMDB00856 | 0.94 | 0.84 | **0.57** | 0.09 |
| Amino Acid | Urea cycle; Arginine and Proline Metabolism | N-acetylproline | | 0.65 | 0.15 | 1.09 | 0.78 |
| Amino Acid | Urea cycle; Arginine and Proline Metabolism | N-delta-acetylornithine | | 0.81 | 0.38 | 0.9 | 0.69 |
| Amino Acid | Urea cycle; Arginine and Proline Metabolism | N-alpha-acetylornithine | HMDB03357 | 0.81 | 0.48 | 1.15 | 0.65 |
| Amino Acid | Urea cycle; Arginine and Proline Metabolism | N2,N5-diacetylornithine | | 0.93 | 0.81 | 0.9 | 0.75 |
| Amino Acid | Urea cycle; Arginine and Proline Metabolism | trans-4-hydroxyproline | HMDB00725 | 0.73 | 0.37 | 1.13 | 0.74 |
| Amino Acid | Urea cycle; Arginine and Proline Metabolism | pro-hydroxy-pro | HMDB06695 | 0.6 | 0.14 | 0.84 | 0.64 |
| Amino Acid | Urea cycle; Arginine and Proline Metabolism | N-methylproline | | 1.08 | 0.89 | 1.57 | 0.42 |
| Amino Acid | Urea cycle; Arginine and Proline Metabolism | 1-propanoylproline | | 1 | | 1 | |
| Amino Acid | Urea cycle; Arginine and Proline Metabolism | N-monomethylarginine | HMDB29416 | 0.61 | 0.1188 | 1.14 | 0.6917 |
| Amino Acid | Urea cycle; Arginine and Proline Metabolism | argininate* | HMDB03148 | 0.77 | 0.4317 | 0.91 | 0.7845 |
| Amino Acid | Creatine Metabolism | guanidinoacetate | HMDB00128 | 0.79 | 0.5288 | 0.54 | 0.1254 |
| Amino Acid | Creatine Metabolism | creatine | HMDB00064 | 0.74 | 0.6977 | 1.72 | 0.4937 |
| Amino Acid | Creatine Metabolism | creatinine | HMDB00562 | 0.31 | 0.1835 | 1.12 | 0.9044 |
| Amino Acid | Creatine Metabolism | N-methylhydantoin | HMDB03646 | 0.92 | 0.8558 | 0.71 | 0.4788 |
| Amino Acid | Creatine Metabolism | N-carbamoylsarcosine | HMDB12265 | 1.3 | 0.5825 | **0.38** | 0.0515 |
| Amino Acid | Polyamine Metabolism | agmatine | HMDB01432 | 0.45 | 0.2150 | 0.68 | 0.5571 |
| Amino Acid | Polyamine Metabolism | putrescine | HMDB01414 | 0.75 | 0.4331 | 0.86 | 0.6775 |
| Amino Acid | Polyamine Metabolism | spermidine | HMDB01257 | 0.9 | 0.8593 | 0.67 | 0.4912 |
| Amino Acid | Polyamine Metabolism | acisoga | | 0.62 | 0.1477 | 0.58 | 0.1154 |
| Amino Acid | Polyamine Metabolism | spermine | HMDB01256 | 1 | | 1 | |
| Amino Acid | Polyamine Metabolism | N(1)-acetylspermine | HMDB01186 | 0.73 | 0.5383 | 0.95 | 0.9281 |
| Amino Acid | Polyamine Metabolism | N1,N12-diacetylspermine | HMDB02172 | 0.76 | 0.5298 | 1.1 | 0.8310 |
| Amino Acid | Polyamine Metabolism | 5-methylthioadenosine (MTA) | HMDB01173 | 1.01 | 0.9678 | 0.85 | 0.6179 |
| Amino Acid | Polyamine Metabolism | N-acetylputrescine | HMDB02064 | **0.51** | 0.0718 | 0.82 | 0.6059 |
| Amino Acid | Polyamine Metabolism | 4-acetamidobutanoate | HMDB03681 | 0.75 | 0.2275 | 1.03 | 0.8946 |
| Amino Acid | Polyamine Metabolism | (N(1) + N(8))-acetylspermidine | | 0.62 | 0.1683 | 1.05 | 0.8857 |
| Amino Acid | Guanidino and Acetamido Metabolism | 1-methylguanidine | HMDB01522 | 0.71 | 0.1919 | 1.48 | 0.1603 |

| | | | | | | | |
|---|---|---|---|---|---|---|---|
| Amino Acid | Guanidino and Acetamido Metabolism | 4-guanidinobutanoate | HMDB03464 | 0.83 | 0.4702 | 1.43 | 0.1771 |
| Amino Acid | Guanidino and Acetamido Metabolism | guanidinosuccinate | HMDB03157 | 0.88 | 0.4943 | 1.06 | 0.7449 |
| Amino Acid | Glutathione Metabolism | cysteine-glutathione disulfide | HMDB00656 | 1.04 | 0.7529 | 1.01 | 0.9102 |
| Amino Acid | Glutathione Metabolism | 5-oxoproline | HMDB00267 | **0.5** | 0.0085 | 1.27 | 0.3819 |
| Amino Acid | Glutathione Metabolism | 2-aminobutyrate | HMDB00650 | 0.54 | 0.1996 | 0.46 | 0.1331 |
| Amino Acid | Glutathione Metabolism | 2-hydroxybutyrate/2-hydroxyisobutyrate | | **0.45** | 0.0153 | 0.73 | 0.3554 |
| Amino Acid | Glutathione Metabolism | 4-amino-2-hydroxybutyrate | | 0.91 | 0.8055 | 0.86 | 0.7278 |
| Peptide | Gamma-glutamyl Amino Acid | gamma-glutamylalanine | HMDB29142 | 0.7 | 0.3539 | 1.39 | 0.4058 |
| Peptide | Gamma-glutamyl Amino Acid | gamma-glutamylglutamate | HMDB11737 | 1.63 | 0.3485 | 1.05 | 0.9258 |
| Peptide | Gamma-glutamyl Amino Acid | gamma-glutamylglutamine | HMDB11738 | **0.49** | 0.0278 | 1.05 | 0.8844 |
| Peptide | Gamma-glutamyl Amino Acid | gamma-glutamylglycine | HMDB11667 | 0.55 | 0.1578 | **2.21** | 0.0724 |
| Peptide | Gamma-glutamyl Amino Acid | gamma-glutamylhistidine | | 0.76 | 0.5057 | 0.96 | 0.9228 |
| Peptide | Gamma-glutamyl Amino Acid | gamma-glutamylisoleucine* | HMDB11170 | 0.91 | 0.7904 | 1.54 | 0.2570 |
| Peptide | Gamma-glutamyl Amino Acid | gamma-glutamylleucine | HMDB11171 | 0.86 | 0.6597 | 1.57 | 0.1996 |
| Peptide | Gamma-glutamyl Amino Acid | gamma-glutamyl-alpha-lysine | | 0.98 | 0.9304 | 1.38 | 0.2266 |
| Peptide | Gamma-glutamyl Amino Acid | gamma-glutamyl-epsilon-lysine | HMDB03869 | **0.49** | 0.0137 | 0.84 | 0.5553 |
| Peptide | Gamma-glutamyl Amino Acid | gamma-glutamylmethionine | HMDB29155 | 1.04 | 0.9377 | 2.11 | 0.1285 |
| Peptide | Gamma-glutamyl Amino Acid | gamma-glutamylphenylalanine | HMDB00594 | 0.7 | 0.3271 | 1.6 | 0.2115 |
| Peptide | Gamma-glutamyl Amino Acid | gamma-glutamylthreonine | HMDB29159 | 0.95 | 0.8991 | 1.1 | 0.8194 |
| Peptide | Gamma-glutamyl Amino Acid | gamma-glutamyltryptophan | HMDB29160 | 0.82 | 0.6082 | 1.4 | 0.3963 |
| Peptide | Gamma-glutamyl Amino Acid | gamma-glutamyltyrosine | HMDB11741 | 0.7 | 0.3668 | 1.59 | 0.2464 |
| Peptide | Gamma-glutamyl Amino Acid | gamma-glutamylvaline | HMDB11172 | 0.67 | 0.4551 | 2.08 | 0.1882 |
| Peptide | Gamma-glutamyl Amino Acid | gamma-glutamyl-2-aminobutyrate | | 1.28 | 0.5706 | 1 | 0.9981 |
| Peptide | Dipeptide | alanylleucine | HMDB28691 | 1.03 | 0.9480 | 0.9 | 0.7961 |
| Peptide | Dipeptide | glycylisoleucine | HMDB28844 | 0.97 | 0.9235 | 0.74 | 0.3017 |
| Peptide | Dipeptide | glycylleucine | HMDB00759 | 0.97 | 0.9184 | 0.77 | 0.3515 |
| Peptide | Dipeptide | glycylvaline | HMDB28854 | 0.89 | 0.6428 | 0.8 | 0.3933 |
| Peptide | Dipeptide | isoleucylglycine | HMDB28907 | 0.93 | 0.8767 | 0.87 | 0.7558 |
| Peptide | Dipeptide | leucylalanine | HMDB28922 | 1.85 | 0.2104 | 1.1 | 0.8464 |
| Peptide | Dipeptide | leucylglycine | HMDB28929 | 1.23 | 0.6105 | 0.81 | 0.6274 |
| Peptide | Dipeptide | lysylleucine | HMDB28955 | 0.71 | 0.4994 | 1.42 | 0.5058 |
| Peptide | Dipeptide | phenylalanylalanine | | 1.22 | 0.5802 | 0.81 | 0.5787 |
| Peptide | Dipeptide | phenylalanylglycine | HMDB28995 | 0.98 | 0.9603 | 0.85 | 0.7296 |

| | | | | | | | |
|---|---|---|---|---|---|---|---|
| Peptide | Dipeptide | prolylglycine | HMDB11178 | 0.52 | 0.1240 | 1.12 | 0.7908 |
| Peptide | Dipeptide | threonylphenylalanine | HMDB29068 | 1.04 | 0.9246 | 0.97 | 0.9504 |
| Peptide | Dipeptide | tryptophylglycine | HMDB29083 | 1.03 | 0.9535 | **0.42** | 0.0957 |
| Peptide | Dipeptide | tyrosylglycine | HMDB29105 | 0.93 | 0.9027 | 0.89 | 0.8454 |
| Peptide | Dipeptide | valylglutamine | HMDB29125 | 0.88 | 0.7982 | 0.75 | 0.5783 |
| Peptide | Dipeptide | valylglycine | HMDB29127 | 0.89 | 0.8232 | 0.96 | 0.9490 |
| Peptide | Dipeptide | valylleucine | HMDB29131 | 1.36 | 0.5069 | 1.04 | 0.9342 |
| Peptide | Dipeptide | leucylglutamine* | HMDB28927 | 1.22 | 0.6496 | 0.88 | 0.7917 |
| Peptide | Polypeptide | alanyl-glutamyl-meso-diaminopimelate | | 0.95 | 0.9094 | 0.49 | 0.1544 |
| Peptide | Acetylated Peptides | phenylacetylalanine | | 1.09 | 0.8203 | 0.89 | 0.7699 |
| Peptide | Acetylated Peptides | phenylacetylhistidine | | 0.89 | 0.6945 | 0.87 | 0.6500 |
| Peptide | Acetylated Peptides | phenylacetylglutamate | HMDB59772 | 1.25 | 0.6201 | 0.97 | 0.9415 |
| Peptide | Acetylated Peptides | phenylacetylglutamine | HMDB06344 | 0.55 | 0.4249 | 0.34 | 0.1717 |
| Peptide | Acetylated Peptides | phenylacetylglycine | HMDB00821 | 0.75 | 0.5903 | 1.11 | 0.8502 |
| Peptide | Acetylated Peptides | 4-hydroxyphenylacetylglycine | | 0.63 | 0.2054 | 1.59 | 0.2175 |
| Peptide | Acetylated Peptides | phenylacetylmethionine | | 0.99 | 0.9875 | 1.42 | 0.3911 |
| Peptide | Acetylated Peptides | phenylacetylphenylalanine | | 1.08 | 0.8611 | 1 | 0.9992 |
| Carbohydrate | Glycolysis, Gluconeogenesis, and Pyruvate Metabolism | 1,5-anhydroglucitol (1,5-AG) | HMDB02712 | 0.91 | 0.5487 | 0.83 | 0.2867 |
| Carbohydrate | Glycolysis, Gluconeogenesis, and Pyruvate Metabolism | glucose | HMDB00122 | 0.92 | 0.8142 | 1.64 | 0.1717 |
| Carbohydrate | Glycolysis, Gluconeogenesis, and Pyruvate Metabolism | pyruvate | HMDB00243 | **0.46** | 0.0237 | 1.26 | 0.5202 |
| Carbohydrate | Glycolysis, Gluconeogenesis, and Pyruvate Metabolism | lactate | HMDB00190 | 0.48 | 0.1483 | 1.04 | 0.9411 |
| Carbohydrate | Glycolysis, Gluconeogenesis, and Pyruvate Metabolism | glycerate | HMDB00139 | 0.84 | 0.5264 | 0.89 | 0.6648 |
| Carbohydrate | Glycolysis, Gluconeogenesis, and Pyruvate Metabolism | glucosylglycerol | | 0.52 | 0.2177 | **0.39** | 0.0941 |
| Carbohydrate | Pentose Metabolism | ribose | HMDB00283 | 1.12 | 0.7537 | **1.9** | 0.0810 |
| Carbohydrate | Pentose Metabolism | ribitol | HMDB00508 | 0.9 | 0.8277 | 2.15 | 0.1384 |
| Carbohydrate | Pentose Metabolism | ribonate | HMDB00867 | 0.63 | 0.4202 | **0.36** | 0.0882 |
| Carbohydrate | Pentose Metabolism | xylose | HMDB00098 | 1.8 | 0.3714 | 2.26 | 0.2409 |
| Carbohydrate | Pentose Metabolism | arabinose | HMDB00646 | 1.66 | 0.3040 | 0.82 | 0.6996 |

| | | | | | | | |
|---|---|---|---|---|---|---|---|
| Carbohydr ate | Pentose Metabolism | fucose | HMDB001 74 | 0.71 | 0.359 2 | 0.85 | 0.663 0 |
| Carbohydr ate | Pentose Metabolism | arabitol/xylitol | | 1.05 | 0.915 3 | 1.01 | 0.976 7 |
| Carbohydr ate | Pentose Metabolism | arabonate/xylonate | | 1.03 | 0.949 0 | 1.24 | 0.633 4 |
| Carbohydr ate | Pentose Metabolism | sedoheptulose | HMDB032 19 | 0.99 | 0.980 5 | 0.48 | 0.236 5 |
| Carbohydr ate | Glycogen Metabolism | maltotetraose | HMDB012 96 | 0.75 | 0.582 2 | 1.62 | 0.365 7 |
| Carbohydr ate | Glycogen Metabolism | maltotriose | HMDB012 62 | 1.17 | 0.743 9 | 0.7 | 0.473 8 |
| Carbohydr ate | Glycogen Metabolism | maltose | HMDB001 63 | 1.11 | 0.830 0 | 0.56 | 0.267 0 |
| Carbohydr ate | Disaccharides and Oligosaccharides | lactose | HMDB001 86 | 1.09 | 0.877 0 | 1.99 | 0.212 4 |
| Carbohydr ate | Disaccharides and Oligosaccharides | lacto-N-fucopentaose I | | 0.97 | 0.866 6 | 0.99 | 0.960 3 |
| Carbohydr ate | Disaccharides and Oligosaccharides | lacto-N-fucopentaose II | | 0.99 | 0.952 8 | 1.05 | 0.859 1 |
| Carbohydr ate | Disaccharides and Oligosaccharides | lacto-N-tetraose | HMDB065 66 | 1.09 | 0.759 9 | 1.06 | 0.825 5 |
| Carbohydr ate | Disaccharides and Oligosaccharides | lacto-N-neotetraose | | 0.63 | 0.285 5 | 1.36 | 0.490 7 |
| Carbohydr ate | Disaccharides and Oligosaccharides | 3-sialyllactose | HMDB008 25 | **0.8** | 0.048 4 | 1 | 1.000 0 |
| Carbohydr ate | Disaccharides and Oligosaccharides | 6'-sialyllactose | HMDB065 69 | 0.94 | 0.797 4 | 0.99 | 0.980 9 |
| Carbohydr ate | Disaccharides and Oligosaccharides | 2-fucosyllactose | HMDB020 98 | 0.8 | 0.664 0 | 2.17 | 0.133 1 |
| Carbohydr ate | Disaccharides and Oligosaccharides | 3-fucosyllactose | HMDB020 94 | 1.03 | 0.948 3 | 1.68 | 0.317 1 |
| Carbohydr ate | Disaccharides and Oligosaccharides | Lewis a trisaccharide | | **1.36** | 0.044 3 | 0.99 | 0.952 0 |
| Carbohydr ate | Disaccharides and Oligosaccharides | Lewis X trisaccharide | HMDB065 68 | 1 | 0.931 3 | 1 | 1.000 0 |
| Carbohydr ate | Disaccharides and Oligosaccharides | sucrose | HMDB002 58 | 1.52 | 0.248 1 | 0.67 | 0.282 2 |
| Carbohydr ate | Fructose, Mannose and Galactose Metabolism | fructose | HMDB006 60 | 1.06 | 0.911 5 | 1.02 | 0.974 8 |
| Carbohydr ate | Fructose, Mannose and Galactose Metabolism | mannitol/sorbitol | HMDB002 47 | 0.94 | 0.891 8 | 1.57 | 0.381 8 |
| Carbohydr ate | Fructose, Mannose and Galactose Metabolism | mannose | HMDB001 69 | 0.83 | 0.677 2 | 2.04 | 0.125 9 |
| Carbohydr ate | Fructose, Mannose and Galactose Metabolism | galactose | HMDB001 43 | 0.8 | 0.449 2 | 0.84 | 0.554 5 |
| Carbohydr ate | Fructose, Mannose and Galactose Metabolism | galactitol (dulcitol) | HMDB001 07 | 0.59 | 0.450 8 | 1.12 | 0.874 7 |
| Carbohydr ate | Fructose, Mannose and Galactose Metabolism | galactonate | HMDB005 65 | 0.85 | 0.771 5 | 1.15 | 0.817 3 |
| Carbohydr ate | Aminosugar Metabolism | glucosamine | HMDB015 14 | 0.68 | 0.207 6 | 1.21 | 0.552 1 |
| Carbohydr ate | Aminosugar Metabolism | glucosamine 6-sulfate | HMDB005 92 | 0.61 | 0.546 1 | 2.1 | 0.378 6 |
| Carbohydr ate | Aminosugar Metabolism | glucuronate | HMDB001 27 | 0.84 | 0.715 6 | 0.82 | 0.685 6 |
| Carbohydr ate | Aminosugar Metabolism | N-acetylglucosamine 6-sulfate | HMDB008 41 | 0.66 | 0.480 6 | 2.53 | 0.131 3 |
| Carbohydr ate | Aminosugar Metabolism | N-acetylneuraminate | HMDB002 30 | 0.69 | 0.390 8 | 1.07 | 0.876 9 |
| Carbohydr ate | Aminosugar Metabolism | N-acetyl-beta-glucosaminylamine | HMDB011 04 | 0.88 | 0.804 7 | 0.83 | 0.720 1 |

| | | | | | | | |
|---|---|---|---|---|---|---|---|
| Carbohydrate | Aminosugar Metabolism | N-acetylmuramate | HMDB60493 | 1.17 | 0.6353 | 1.6 | 0.1651 |
| Carbohydrate | Aminosugar Metabolism | 6-sialyl-N-acetyllactosamine | HMDB06584 | 1.05 | 0.8548 | 1.57 | 0.1345 |
| Carbohydrate | Aminosugar Metabolism | N-acetylglucosaminylasparagine | HMDB00489 | 0.65 | 0.2608 | 1.17 | 0.6864 |
| Carbohydrate | Aminosugar Metabolism | erythronate* | HMDB00613 | 0.75 | 0.4044 | 1.3 | 0.4606 |
| Carbohydrate | Aminosugar Metabolism | N-acetylglucosamine/N-acetylgalactosamine | HMDB00215 | 0.9 | 0.7447 | 1.15 | 0.6902 |
| Carbohydrate | Advanced Glycation End-product | N6-carboxymethyllysine | | 0.68 | 0.1001 | 1.36 | 0.2126 |
| Energy | TCA Cycle | citrate | HMDB00094 | 0.56 | 0.3396 | 0.46 | 0.2271 |
| Energy | TCA Cycle | aconitate [cis or trans] | | 1.44 | 0.5431 | 1.03 | 0.9567 |
| Energy | TCA Cycle | isocitric lactone | | 0.53 | 0.3503 | 1.55 | 0.5414 |
| Energy | TCA Cycle | alpha-ketoglutarate | HMDB00208 | 0.65 | 0.1785 | **1.96** | 0.0484 |
| Energy | TCA Cycle | succinylcarnitine (C4-DC) | HMDB61717 | 0.68 | 0.3075 | 0.99 | 0.9777 |
| Energy | TCA Cycle | succinate | HMDB00254 | 1.18 | 0.6810 | 0.87 | 0.7303 |
| Energy | TCA Cycle | fumarate | HMDB00134 | 1.04 | 0.9307 | 1.06 | 0.8993 |
| Energy | TCA Cycle | malate | HMDB00156 | 1.19 | 0.5626 | 0.88 | 0.6846 |
| Energy | TCA Cycle | tricarballylate | HMDB31193 | 0.76 | 0.4707 | 0.75 | 0.4546 |
| Energy | TCA Cycle | citraconate/glutaconate | | 1.06 | 0.7325 | 1.12 | 0.5480 |
| Energy | TCA Cycle | 2-methylcitrate/homocitrate | | 1.12 | 0.7784 | 0.8 | 0.6043 |
| Energy | Oxidative Phosphorylation | phosphate | HMDB01429 | 2.04 | 0.2073 | 1.2 | 0.7520 |
| Lipid | Fatty Acid Synthesis | malonylcarnitine | HMDB02095 | **0.62** | 0.0871 | 1 | 0.9969 |
| Lipid | Fatty Acid Synthesis | malonate | HMDB00691 | 1.19 | 0.5826 | 1.01 | 0.9820 |
| Lipid | Short Chain Fatty Acid | valerate (5:0) | HMDB00892 | 1.19 | 0.7583 | 1.95 | 0.2713 |
| Lipid | Medium Chain Fatty Acid | caproate (6:0) | HMDB00535 | 0.85 | 0.7455 | 1.21 | 0.7182 |
| Lipid | Medium Chain Fatty Acid | heptanoate (7:0) | HMDB00666 | 0.64 | 0.2881 | 0.87 | 0.7561 |
| Lipid | Medium Chain Fatty Acid | caprylate (8:0) | HMDB00482 | 1.06 | 0.7535 | 0.94 | 0.7415 |
| Lipid | Medium Chain Fatty Acid | caprate (10:0) | HMDB00511 | 0.87 | 0.5483 | 1.03 | 0.9026 |
| Lipid | Medium Chain Fatty Acid | undecanoate (11:0) | HMDB00947 | 0.95 | 0.7658 | 0.9 | 0.5437 |
| Lipid | Medium Chain Fatty Acid | laurate (12:0) | HMDB00638 | 0.65 | 0.2049 | 0.78 | 0.4782 |
| Lipid | Medium Chain Fatty Acid | 5-dodecenoate (12:1n7) | HMDB00529 | 1 | 0.9978 | 0.87 | 0.6846 |
| Lipid | Long Chain Fatty Acid | myristate (14:0) | HMDB00806 | 0.75 | 0.3197 | 0.86 | 0.6189 |
| Lipid | Long Chain Fatty Acid | myristoleate (14:1n5) | HMDB02000 | 1.03 | 0.8923 | 1.1 | 0.7136 |
| Lipid | Long Chain Fatty Acid | pentadecanoate (15:0) | HMDB00826 | 1.15 | 0.6403 | 1.06 | 0.8358 |
| Lipid | Long Chain Fatty Acid | palmitate (16:0) | HMDB00220 | 0.88 | 0.5893 | 1.17 | 0.5041 |
| Lipid | Long Chain Fatty Acid | palmitoleate (16:1n7) | HMDB03229 | 0.94 | 0.8218 | 0.97 | 0.9154 |
| Lipid | Long Chain Fatty | margarate (17:0) | HMDB022 | 0.87 | 0.631 | 1.19 | 0.569 |

| | | | | | | | |
|---|---|---|---|---|---|---|---|
| | Acid | | 59 | | 6 | | 0 |
| Lipid | Long Chain Fatty Acid | 10-heptadecenoate (17:1n7) | HMDB60038 | 0.89 | 0.7284 | 0.87 | 0.6666 |
| Lipid | Long Chain Fatty Acid | trans-heptadecenoate (tr 17:1)* | | 0.98 | 0.9629 | 1.17 | 0.6612 |
| Lipid | Long Chain Fatty Acid | stearate (18:0) | HMDB00827 | 0.94 | 0.8217 | 1.13 | 0.6366 |
| Lipid | Long Chain Fatty Acid | oleate/vaccenate (18:1) | | 0.89 | 0.6700 | 1.34 | 0.3059 |
| Lipid | Long Chain Fatty Acid | nonadecanoate (19:0) | HMDB00772 | 0.9 | 0.7441 | 1.55 | 0.1783 |
| Lipid | Long Chain Fatty Acid | 10-nonadecenoate (19:1n9) | HMDB13622 | 0.65 | 0.1860 | 0.94 | 0.8616 |
| Lipid | Long Chain Fatty Acid | trans-nonadecenoate (tr 19:1)* | | 0.97 | 0.9194 | 1.06 | 0.8608 |
| Lipid | Long Chain Fatty Acid | arachidate (20:0) | HMDB02212 | 0.98 | 0.9479 | **1.72** | 0.0772 |
| Lipid | Long Chain Fatty Acid | eicosenoate (20:1) | HMDB02231 | 0.73 | 0.2877 | 1.38 | 0.2895 |
| Lipid | Long Chain Fatty Acid | erucate (22:1n9) | HMDB02068 | 0.8 | 0.5187 | 1.26 | 0.5072 |
| Lipid | Polyunsaturated Fatty Acid (n3 and n6) | heneicosapentaenoate (21:5n3) | | 0.96 | 0.8612 | 0.84 | 0.4439 |
| Lipid | Polyunsaturated Fatty Acid (n3 and n6) | hexadecadienoate (16:2n6) | HMDB00477 | 0.85 | 0.5093 | 1.01 | 0.9713 |
| Lipid | Polyunsaturated Fatty Acid (n3 and n6) | hexadecatrienoate (16:3n3) | | 0.98 | 0.9430 | 0.81 | 0.5486 |
| Lipid | Polyunsaturated Fatty Acid (n3 and n6) | stearidonate (18:4n3) | HMDB06547 | **0.43** | 0.0511 | 1.37 | 0.4840 |
| Lipid | Polyunsaturated Fatty Acid (n3 and n6) | eicosapentaenoate (EPA; 20:5n3) | HMDB01999 | 0.62 | 0.1691 | 0.88 | 0.7232 |
| Lipid | Polyunsaturated Fatty Acid (n3 and n6) | docosapentaenoate (n3 DPA; 22:5n3) | HMDB06528 | 0.91 | 0.7939 | 0.95 | 0.9003 |
| Lipid | Polyunsaturated Fatty Acid (n3 and n6) | docosahexaenoate (DHA; 22:6n3) | HMDB02183 | **0.49** | 0.0635 | 0.96 | 0.9172 |
| Lipid | Polyunsaturated Fatty Acid (n3 and n6) | docosatrienoate (22:3n3) | HMDB02823 | 0.77 | 0.5258 | 0.94 | 0.8822 |
| Lipid | Polyunsaturated Fatty Acid (n3 and n6) | omega-3 arachidonate (20:4n3) | HMDB02177 | 1.04 | 0.7929 | 1 | 1.0000 |
| Lipid | Polyunsaturated Fatty Acid (n3 and n6) | nisinate (24:6n3) | HMDB02007 | 0.83 | 0.6598 | 0.78 | 0.5691 |
| Lipid | Polyunsaturated Fatty Acid (n3 and n6) | linoleate (18:2n6) | HMDB00673 | 1.12 | 0.7118 | **1.69** | 0.0921 |
| Lipid | Polyunsaturated Fatty Acid (n3 and n6) | linolenate [alpha or gamma; (18:3n3 or 6)] | HMDB03073 | 1.28 | 0.4446 | 1.69 | 0.1224 |
| Lipid | Polyunsaturated Fatty Acid (n3 and n6) | dihomo-linolenate (20:3n3 or n6) | HMDB02925 | 1.01 | 0.9706 | 1.01 | 0.9831 |
| Lipid | Polyunsaturated Fatty Acid (n3 and n6) | arachidonate (20:4n6) | HMDB01043 | 0.7 | 0.2967 | 0.92 | 0.8166 |
| Lipid | Polyunsaturated Fatty Acid (n3 and n6) | adrenate (22:4n6) | HMDB02226 | 1 | 0.9901 | 0.78 | 0.5377 |
| Lipid | Polyunsaturated Fatty Acid (n3 and | docosapentaenoate (n6 DPA; 22:5n6) | HMDB01976 | 0.81 | 0.6475 | 0.91 | 0.8450 |

| | | | | | | | |
|---|---|---|---|---|---|---|---|
| | n6) | | | | | | |
| Lipid | Polyunsaturated Fatty Acid (n3 and n6) | docosadienoate (22:2n6) | HMDB61714 | 0.74 | 0.4620 | 0.87 | 0.7358 |
| Lipid | Polyunsaturated Fatty Acid (n3 and n6) | dihomo-linoleate (20:2n6) | HMDB05060 | 0.69 | 0.3007 | 0.93 | 0.8487 |
| Lipid | Polyunsaturated Fatty Acid (n3 and n6) | mead acid (20:3n9) | HMDB10378 | 0.45 | 0.2616 | 1 | 0.9950 |
| Lipid | Polyunsaturated Fatty Acid (n3 and n6) | docosatrienoate (22:3n6)* | | 0.88 | 0.7963 | 0.98 | 0.9635 |
| Lipid | Fatty Acid Hydroxyl Fatty Acid | PAHSA (16:0/OH-18:0) | | 0.61 | 0.1742 | 0.96 | 0.9202 |
| Lipid | Fatty Acid Hydroxyl Fatty Acid | LAHSA (18:2/OH-18:0)* | | 0.94 | 0.8699 | 1.17 | 0.7061 |
| Lipid | Fatty Acid Hydroxyl Fatty Acid | OAHSA (18:1/OH-18:0) | | 0.81 | 0.5961 | 1.1 | 0.8296 |
| Lipid | Fatty Acid, Branched | isocaproate (i6:0) | HMDB00689 | 0.94 | 0.9162 | 1.71 | 0.3463 |
| Lipid | Fatty Acid, Branched | 13-methylmyristate (i15:0) | | 0.92 | 0.8472 | 0.94 | 0.8866 |
| Lipid | Fatty Acid, Branched | 15-methylpalmitate (i17:0) | | 0.92 | 0.7691 | 1.15 | 0.6476 |
| Lipid | Fatty Acid, Branched | 15-methylmargarate (a18:0) | | 1.31 | 0.4471 | 1.15 | 0.7031 |
| Lipid | Fatty Acid, Branched | 17-methylstearate (i19:0) | HMDB37397 | 0.88 | 0.6744 | 1.25 | 0.4533 |
| Lipid | Fatty Acid, Branched | pristanate | HMDB00795 | 1.32 | 0.6127 | 1.47 | 0.4875 |
| Lipid | Fatty Acid, Dicarboxylate | dimethylmalonic acid | HMDB02001 | 0.88 | 0.6576 | 1.45 | 0.2145 |
| Lipid | Fatty Acid, Dicarboxylate | glutarate (C5-DC) | HMDB00661 | 1.02 | 0.9634 | 0.75 | 0.4547 |
| Lipid | Fatty Acid, Dicarboxylate | 3-methylglutarate/2-methylglutarate | HMDB00752 | 0.86 | 0.6382 | 0.93 | 0.8216 |
| Lipid | Fatty Acid, Dicarboxylate | 2-hydroxyglutarate | HMDB00606 | 0.67 | 0.2314 | 1.69 | 0.1328 |
| Lipid | Fatty Acid, Dicarboxylate | adipate (C6-DC) | HMDB00448 | 0.7 | 0.1357 | 1.18 | 0.5019 |
| Lipid | Fatty Acid, Dicarboxylate | 2-hydroxyadipate | HMDB00321 | 0.87 | 0.6449 | 1.12 | 0.7298 |
| Lipid | Fatty Acid, Dicarboxylate | 3-methyladipate | HMDB00555 | 0.64 | 0.2408 | 0.63 | 0.2441 |
| Lipid | Fatty Acid, Dicarboxylate | maleate | HMDB00176 | 0.84 | 0.3419 | 1.14 | 0.4991 |
| Lipid | Fatty Acid, Dicarboxylate | pimelate (C7-DC) | HMDB00857 | 0.93 | 0.7811 | **2.13** | 0.0078 |
| Lipid | Fatty Acid, Dicarboxylate | suberate (C8-DC) | HMDB00893 | 0.95 | 0.7889 | 1 | 0.9837 |
| Lipid | Fatty Acid, Dicarboxylate | azelate (C9-DC) | HMDB00784 | 1 | 0.9925 | 1.18 | 0.3261 |
| Lipid | Fatty Acid, Dicarboxylate | sebacate (C10-DC) | HMDB00792 | 1.01 | 0.9484 | 0.94 | 0.7680 |
| Lipid | Fatty Acid, Dicarboxylate | undecanedioate (C11-DC) | HMDB00888 | 0.93 | 0.6061 | 0.99 | 0.9272 |
| Lipid | Fatty Acid, Dicarboxylate | dodecanedioate (C12-DC) | HMDB00623 | 1.02 | 0.9012 | 1.08 | 0.6024 |
| Lipid | Fatty Acid, Dicarboxylate | tetradecanedioate (C14-DC) | HMDB00872 | 1.24 | 0.1242 | 0.98 | 0.8885 |
| Lipid | Fatty Acid, Dicarboxylate | hexadecanedioate (C16-DC) | HMDB00672 | 1.32 | 0.1153 | 0.91 | 0.6133 |
| Lipid | Fatty Acid, Amide | oleamide | HMDB02117 | 1.28 | 0.5222 | 0.62 | 0.2324 |
| Lipid | Fatty Acid, Amide | linoleamide (18:2n6) | | 1.57 | 0.3321 | 0.67 | 0.4009 |

| | | | | | | | |
|---|---|---|---|---|---|---|---|
| Lipid | Fatty Acid, Amino | 2-aminoheptanoate | | **0.55** | 0.0859 | 1.03 | 0.9379 |
| Lipid | Fatty Acid Metabolism (also BCAA Metabolism) | butyrylglycine | HMDB00808 | 0.99 | 0.9822 | 2.3 | 0.1120 |
| Lipid | Fatty Acid Metabolism (also BCAA Metabolism) | propionylcarnitine (C3) | HMDB00824 | 0.64 | 0.3322 | 1.4 | 0.4834 |
| Lipid | Fatty Acid Metabolism (also BCAA Metabolism) | propionylglycine | HMDB00783 | 0.85 | 0.7333 | 1.12 | 0.8216 |
| Lipid | Fatty Acid Metabolism (also BCAA Metabolism) | methylmalonate (MMA) | HMDB00202 | 0.92 | 0.8832 | 1.41 | 0.5425 |
| Lipid | Fatty Acid Metabolism (Acyl Glutamine) | hexanoylglutamine | | 0.64 | 0.3880 | 0.44 | 0.1206 |
| Lipid | Fatty Acid Metabolism(Acyl Glycine) | valerylglycine | HMDB00927 | 0.63 | 0.3034 | 1.73 | 0.2463 |
| Lipid | Fatty Acid Metabolism(Acyl Glycine) | hexanoylglycine | HMDB00701 | 0.81 | 0.6930 | 1.33 | 0.6114 |
| Lipid | Fatty Acid Metabolism(Acyl Glycine) | heptanoyl glycine | | 1.18 | 0.6975 | 1.27 | 0.5871 |
| Lipid | Fatty Acid Metabolism(Acyl Glycine) | N-octanoylglycine | HMDB00832 | 1 | 1.0000 | 0.88 | 0.2990 |
| Lipid | Fatty Acid Metabolism(Acyl Glycine) | N-palmitoylglycine | HMDB13034 | 1.03 | 0.9342 | 1.15 | 0.6711 |
| Lipid | Fatty Acid Metabolism(Acyl Glycine) | N-oleoylglycine | HMDB13631 | 1.5 | 0.4704 | 1.16 | 0.7939 |
| Lipid | Fatty Acid Metabolism(Acyl Glycine) | N-linoleoylglycine | | 0.85 | 0.6966 | 1.98 | 0.1290 |
| Lipid | Fatty Acid Metabolism(Acyl Carnitine) | acetylcarnitine (C2) | HMDB00201 | 1.79 | 0.4574 | 0.79 | 0.7780 |
| Lipid | Fatty Acid Metabolism(Acyl Carnitine) | 3-hydroxybutyrylcarnitine (1) | HMDB13127 | 1.08 | 0.7806 | 1.25 | 0.4618 |
| Lipid | Fatty Acid Metabolism(Acyl Carnitine) | 3-hydroxybutyrylcarnitine (2) | HMDB13127 | 0.78 | 0.5784 | 1.95 | 0.1543 |
| Lipid | Fatty Acid Metabolism(Acyl Carnitine) | hexanoylcarnitine (C6) | HMDB00705 | 0.76 | 0.3067 | 1.32 | 0.3367 |
| Lipid | Fatty Acid Metabolism(Acyl Carnitine) | octanoylcarnitine (C8) | HMDB00791 | 0.82 | 0.6618 | 1.79 | 0.2048 |
| Lipid | Fatty Acid Metabolism(Acyl Carnitine) | decanoylcarnitine (C10) | HMDB00651 | 0.85 | 0.7380 | 0.86 | 0.7769 |
| Lipid | Fatty Acid Metabolism(Acyl Carnitine) | cis-4-decenoylcarnitine (C10:1) | | 1.18 | 0.7960 | 1.62 | 0.4526 |
| Lipid | Fatty Acid Metabolism(Acyl Carnitine) | laurylcarnitine (C12) | HMDB02250 | 0.84 | 0.7964 | 0.9 | 0.8777 |
| Lipid | Fatty Acid Metabolism(Acyl Carnitine) | myristoylcarnitine (C14) | HMDB05066 | 1.02 | 0.9666 | 0.61 | 0.4021 |
| Lipid | Fatty Acid Metabolism(Acyl Carnitine) | palmitoylcarnitine (C16) | HMDB00222 | 0.84 | 0.7123 | **0.45** | 0.0985 |

| | | | | | | | |
|---|---|---|---|---|---|---|---|
| Lipid | Fatty Acid Metabolism(Acyl Carnitine) | palmitoleoylcarnitine (C16:1)* | | 1 | 0.9981 | 1.14 | 0.7913 |
| Lipid | Fatty Acid Metabolism(Acyl Carnitine) | stearoylcarnitine (C18) | HMDB00848 | 0.75 | 0.2644 | 0.7 | 0.1879 |
| Lipid | Fatty Acid Metabolism(Acyl Carnitine) | linoleoylcarnitine (C18:2)* | HMDB06469 | 1.18 | 0.7219 | 0.6 | 0.2937 |
| Lipid | Fatty Acid Metabolism(Acyl Carnitine) | linolenoylcarnitine (C18:3)* | | 0.99 | 0.9648 | 0.71 | 0.2768 |
| Lipid | Fatty Acid Metabolism(Acyl Carnitine) | oleoylcarnitine (C18:1) | HMDB05065 | 0.88 | 0.7615 | 0.48 | 0.1032 |
| Lipid | Fatty Acid Metabolism(Acyl Carnitine) | myristoleoylcarnitine (C14:1)* | | 0.9 | 0.8642 | 2.06 | 0.2538 |
| Lipid | Fatty Acid Metabolism(Acyl Carnitine) | suberoylcarnitine (C8-DC) | | 0.88 | 0.7835 | 1.65 | 0.3025 |
| Lipid | Fatty Acid Metabolism(Acyl Carnitine) | adipoylcarnitine (C6-DC) | HMDB61677 | 0.61 | 0.1821 | 1.5 | 0.2980 |
| Lipid | Fatty Acid Metabolism(Acyl Carnitine) | arachidoylcarnitine (C20)* | HMDB06460 | 0.88 | 0.6527 | 0.93 | 0.8083 |
| Lipid | Fatty Acid Metabolism(Acyl Carnitine) | arachidonoylcarnitine (C20:4) | | 0.69 | 0.1752 | 0.75 | 0.3075 |
| Lipid | Fatty Acid Metabolism(Acyl Carnitine) | behenoylcarnitine (C22)* | | 0.81 | 0.4088 | 1.27 | 0.3748 |
| Lipid | Fatty Acid Metabolism(Acyl Carnitine) | dihomo-linolenoylcarnitine (20:3n3 or 6)* | | 0.81 | 0.4679 | 0.72 | 0.2645 |
| Lipid | Fatty Acid Metabolism(Acyl Carnitine) | dihomo-linoleoylcarnitine (C20:2)* | | 0.89 | 0.6805 | 0.79 | 0.4223 |
| Lipid | Fatty Acid Metabolism(Acyl Carnitine) | eicosenoylcarnitine (C20:1)* | | 0.82 | 0.4790 | 0.85 | 0.5950 |
| Lipid | Fatty Acid Metabolism(Acyl Carnitine) | erucoylcarnitine (C22:1)* | | 0.74 | 0.3224 | 0.88 | 0.6794 |
| Lipid | Fatty Acid Metabolism(Acyl Carnitine) | docosahexaenoylcarnitine (C22:6)* | | 0.7 | 0.3946 | 0.89 | 0.7902 |
| Lipid | Fatty Acid Metabolism(Acyl Carnitine) | lignoceroylcarnitine (C24)* | | 0.83 | 0.4163 | 1.19 | 0.4605 |
| Lipid | Fatty Acid Metabolism(Acyl Carnitine) | margaroylcarnitine (C17)* | HMDB06210 | 0.82 | 0.5110 | 0.89 | 0.6912 |
| Lipid | Fatty Acid Metabolism(Acyl Carnitine) | nervonoylcarnitine (C24:1)* | | 0.92 | 0.7505 | 0.93 | 0.8089 |
| Lipid | Fatty Acid Metabolism(Acyl Carnitine) | cerotoylcarnitine (C26)* | HMDB06347 | 1.01 | 0.9696 | 0.76 | 0.2926 |
| Lipid | Fatty Acid Metabolism(Acyl Carnitine) | ximenoylcarnitine (C26:1)* | | 0.93 | 0.7411 | 0.85 | 0.4903 |
| Lipid | Fatty Acid Metabolism(Acyl Carnitine) | pentadecanoylcarnitine (C15)* | | 0.84 | 0.6347 | 0.92 | 0.8269 |
| Lipid | Carnitine Metabolism | deoxycarnitine | HMDB01161 | 0.7 | 0.1769 | 0.67 | 0.1374 |

| | | | | | | | |
|---|---|---|---|---|---|---|---|
| Lipid | Carnitine Metabolism | carnitine | HMDB00062 | 0.95 | 0.8271 | 1 | 0.9957 |
| Lipid | Ketone Bodies | 3-hydroxybutyrate (BHBA) | HMDB00357 | 1.75 | 0.1445 | 0.83 | 0.6402 |
| Lipid | Neurotransmitter | acetylcholine | HMDB00895 | 0.67 | 0.5419 | 2.66 | 0.1516 |
| Lipid | Fatty Acid Metabolism (Acyl Choline) | palmitoylcholine | | 1.5 | 0.2667 | 1.38 | 0.3921 |
| Lipid | Fatty Acid Metabolism (Acyl Choline) | oleoylcholine | | **2.07** | 0.0634 | **2.17** | 0.0568 |
| Lipid | Fatty Acid Metabolism (Acyl Choline) | palmitoloelycholine | | 1.17 | 0.1301 | **0.74** | 0.0070 |
| Lipid | Fatty Acid Metabolism (Acyl Choline) | linoleoylcholine* | | **1.85** | 0.0459 | 1.5 | 0.2118 |
| Lipid | Fatty Acid Metabolism (Acyl Choline) | stearoylcholine* | | 1.18 | 0.2793 | 0.92 | 0.5906 |
| Lipid | Fatty Acid, Monohydroxy | 4-hydroxybutyrate (GHB) | HMDB00710 | 1.31 | 0.3641 | 1.32 | 0.3721 |
| Lipid | Fatty Acid, Monohydroxy | 2-hydroxydecanoate | | 0.92 | 0.6194 | 1 | 0.9799 |
| Lipid | Fatty Acid, Monohydroxy | 2-hydroxypalmitate | HMDB31057 | 1.06 | 0.8682 | 0.88 | 0.7384 |
| Lipid | Fatty Acid, Monohydroxy | 2-hydroxystearate | | 1.19 | 0.5186 | 0.9 | 0.6972 |
| Lipid | Fatty Acid, Monohydroxy | 3-hydroxypropanoate | HMDB00700 | 1 | 1.0000 | 0.91 | 0.5915 |
| Lipid | Fatty Acid, Monohydroxy | 3-hydroxysuberate | HMDB00325 | 0.81 | 0.3547 | 0.83 | 0.4350 |
| Lipid | Fatty Acid, Monohydroxy | 3-hydroxyhexanoate | | 1.28 | 0.4908 | 0.9 | 0.7714 |
| Lipid | Fatty Acid, Monohydroxy | 3-hydroxyoctanoate | HMDB01954 | 0.97 | 0.8562 | 1.01 | 0.9580 |
| Lipid | Fatty Acid, Monohydroxy | 3-hydroxydecanoate | HMDB02203 | 0.96 | 0.8622 | 1.05 | 0.8125 |
| Lipid | Fatty Acid, Monohydroxy | 3-hydroxysebacate | HMDB00350 | 0.91 | 0.7490 | 0.94 | 0.8516 |
| Lipid | Fatty Acid, Monohydroxy | 3-hydroxylaurate | HMDB00387 | 0.94 | 0.7528 | 1.14 | 0.5134 |
| Lipid | Fatty Acid, Monohydroxy | 3-hydroxymyristate | | 0.94 | 0.7735 | 0.96 | 0.8341 |
| Lipid | Fatty Acid, Monohydroxy | 3-hydroxypalmitate | HMDB10734 | 1.19 | 0.6388 | 1.01 | 0.9723 |
| Lipid | Fatty Acid, Monohydroxy | 5-hydroxyhexanoate | HMDB00525 | **0.55** | 0.0523 | 0.92 | 0.8074 |
| Lipid | Fatty Acid, Monohydroxy | 5-hydroxydecanoate | HMDB40329 | 0.95 | 0.7920 | 1.09 | 0.6620 |
| Lipid | Fatty Acid, Monohydroxy | 8-hydroxyoctanoate | HMDB61914 | 0.75 | 0.1576 | **1.61** | 0.0224 |
| Lipid | Fatty Acid, Monohydroxy | 13-HODE + 9-HODE | | 1.32 | 0.4913 | 1.36 | 0.4661 |
| Lipid | Fatty Acid, Monohydroxy | 14-HDoHE/17-HDoHE | | 0.87 | 0.7837 | 1.85 | 0.2342 |
| Lipid | Fatty Acid, Monohydroxy | 10-hydroxystearate | HMDB37396 | 0.82 | 0.4929 | 1.41 | 0.2483 |
| Lipid | Fatty Acid, Monohydroxy | 3-hydroxystearate | | 1.36 | 0.3525 | 1.24 | 0.5362 |
| Lipid | Fatty Acid, Monohydroxy | 2-hydroxylaurate | | 1 | 0.9882 | 0.77 | 0.1940 |
| Lipid | Fatty Acid, Monohydroxy | 3-hydroxyvalerate | HMDB00531 | 2.11 | 0.1131 | 1.33 | 0.5630 |
| Lipid | Fatty Acid, Dihydroxy | 12,13-DiHOME | HMDB04705 | 0.89 | 0.7839 | **2.78** | 0.0246 |
| Lipid | Fatty Acid, | 9,10-DiHOME | HMDB04704 | 0.9 | 0.781 | **3.75** | 0.001 |

| | | | | | | | |
|---|---|---|---|---|---|---|---|
| | Dihydroxy | | | | 6 | | 2 |
| Lipid | Endocannabinoid | oleoyl ethanolamide | HMDB02088 | 0.84 | 0.5659 | **1.68** | 0.0881 |
| Lipid | Endocannabinoid | myristoyl ethanolamide | | 0.88 | 0.6254 | 0.92 | 0.7693 |
| Lipid | Endocannabinoid | palmitoyl ethanolamide | HMDB02100 | 0.89 | 0.6443 | 1.08 | 0.7486 |
| Lipid | Endocannabinoid | stearoyl ethanolamide | HMDB13078 | 0.84 | 0.4626 | 1.08 | 0.7485 |
| Lipid | Endocannabinoid | dihomo-linolenoyl ethanolamide | HMDB13625 | 1.1 | 0.7301 | 0.81 | 0.4638 |
| Lipid | Endocannabinoid | arachidonoyl ethanolamide | HMDB04080 | 0.96 | 0.9331 | 0.46 | 0.1109 |
| Lipid | Endocannabinoid | N-oleoyltaurine | | 0.63 | 0.2938 | 1.41 | 0.4566 |
| Lipid | Endocannabinoid | N-stearoyltaurine | | 1.16 | 0.6978 | 0.98 | 0.9637 |
| Lipid | Endocannabinoid | N-palmitoyltaurine | | 0.84 | 0.7077 | 1.09 | 0.8682 |
| Lipid | Endocannabinoid | linoleoyl ethanolamide | HMDB12252 | 0.86 | 0.6820 | 1.22 | 0.6049 |
| Lipid | Endocannabinoid | arachidoyl ethanolamide (20:0)* | | 1.03 | 0.9219 | 1.29 | 0.3824 |
| Lipid | Endocannabinoid | behenoyl ethanolamide (22:0)* | | 1.15 | 0.6814 | 1.01 | 0.9767 |
| Lipid | Endocannabinoid | lignoceroyl ethanolamide (24:0)* | | 1.24 | 0.5184 | 0.88 | 0.7195 |
| Lipid | Endocannabinoid | palmitoleoyl ethanolamide* | HMDB13648 | 0.71 | 0.1952 | 1 | 0.9959 |
| Lipid | Endocannabinoid | margaroyl ethanolamide* | | 0.9 | 0.6750 | 1.07 | 0.8087 |
| Lipid | Endocannabinoid | N-oleoylserine | | 0.74 | 0.3353 | **1.8** | 0.0733 |
| Lipid | Endocannabinoid | N-palmitoylserine | | 1.79 | 0.2014 | 0.65 | 0.3592 |
| Lipid | Inositol Metabolism | myo-inositol | HMDB00211 | 0.81 | 0.6592 | **2.69** | 0.0526 |
| Lipid | Inositol Metabolism | chiro-inositol | HMDB34220 | 0.66 | 0.4460 | 2.4 | 0.1245 |
| Lipid | Inositol Metabolism | pinitol | HMDB34219 | 0.58 | 0.3614 | 0.82 | 0.7475 |
| Lipid | Inositol Metabolism | inositol 1-phosphate (I1P) | HMDB00213 | **2.15** | 0.0554 | 0.84 | 0.6651 |
| Lipid | Phospholipid Metabolism | choline | HMDB00097 | **0.64** | 0.0858 | 1.02 | 0.9402 |
| Lipid | Phospholipid Metabolism | choline phosphate | HMDB01565 | 1.03 | 0.9378 | 1.04 | 0.9389 |
| Lipid | Phospholipid Metabolism | glycerophosphorylcholine (GPC) | HMDB00086 | 0.72 | 0.6045 | 1.08 | 0.9117 |
| Lipid | Phospholipid Metabolism | glycerophosphoethanolamine | HMDB00114 | 1.3 | 0.6092 | 0.87 | 0.8008 |
| Lipid | Phospholipid Metabolism | glycerophosphoserine* | | 1.22 | 0.5868 | 1.23 | 0.6048 |
| Lipid | Phospholipid Metabolism | glycerophosphoinositol* | | 1.08 | 0.8440 | 0.76 | 0.5102 |
| Lipid | Phospholipid Metabolism | trimethylamine N-oxide | HMDB00925 | 1.11 | 0.7502 | 0.78 | 0.4701 |
| Lipid | Phosphatidylcholine (PC) | 1,2-dipalmitoyl-GPC (16:0/16:0) | HMDB00564 | 0.64 | 0.4054 | 0.73 | 0.5839 |
| Lipid | Phosphatidylcholine (PC) | 1-palmitoyl-2-stearoyl-GPC (16:0/18:0) | HMDB07970 | 1.3 | 0.4483 | 0.94 | 0.8672 |
| Lipid | Phosphatidylcholine (PC) | 1-palmitoyl-2-oleoyl-GPC (16:0/18:1) | HMDB07972 | 0.62 | 0.2562 | 0.89 | 0.7825 |
| Lipid | Phosphatidylcholine (PC) | 1-palmitoyl-2-linoleoyl-GPC (16:0/18:2) | HMDB07973 | 0.75 | 0.4039 | 1.04 | 0.9223 |
| Lipid | Phosphatidylcholine (PC) | 1-stearoyl-2-oleoyl-GPC (18:0/18:1) | HMDB08038 | 1 | 0.9993 | 1.12 | 0.6255 |

| | | | | | | | |
|---|---|---|---|---|---|---|---|
| Lipid | Lysophospholipid | 1-palmitoyl-GPC (16:0) | HMDB10382 | 0.63 | 0.4555 | 0.73 | 0.6209 |
| Lipid | Lysophospholipid | 2-palmitoyl-GPC (16:0)* | HMDB61702 | 0.93 | 0.8383 | 0.78 | 0.5354 |
| Lipid | Lysophospholipid | 1-stearoyl-GPC (18:0) | HMDB10384 | 0.62 | 0.4491 | 0.9 | 0.8662 |
| Lipid | Lysophospholipid | 1-oleoyl-GPC (18:1) | HMDB02815 | 1.03 | 0.9559 | 1 | 0.9996 |
| Lipid | Lysophospholipid | 1-linoleoyl-GPC (18:2) | HMDB10386 | 1.07 | 0.8771 | 1.17 | 0.7419 |
| Lipid | Lysophospholipid | 1-palmitoyl-GPE (16:0) | HMDB11503 | 1.75 | 0.1966 | 1.02 | 0.9689 |
| Lipid | Lysophospholipid | 1-stearoyl-GPE (18:0) | HMDB11130 | 0.92 | 0.8812 | 1.01 | 0.9846 |
| Lipid | Lysophospholipid | 2-stearoyl-GPE (18:0)* | HMDB11129 | 1.1 | 0.8268 | 1.12 | 0.8074 |
| Lipid | Lysophospholipid | 1-oleoyl-GPE (18:1) | HMDB11506 | 0.85 | 0.6745 | 0.98 | 0.9524 |
| Lipid | Lysophospholipid | 1-linoleoyl-GPE (18:2)* | HMDB11507 | 1.26 | 0.3214 | 0.98 | 0.9309 |
| Lipid | Lysophospholipid | 1-palmitoyl-GPS (16:0)* | | 1.41 | 0.2138 | 1.09 | 0.7760 |
| Lipid | Lysophospholipid | 1-stearoyl-GPS (18:0)* | | 0.99 | 0.9836 | 0.88 | 0.8183 |
| Lipid | Lysophospholipid | 1-palmitoyl-GPG (16:0)* | | 0.89 | 0.7767 | 0.82 | 0.6487 |
| Lipid | Lysophospholipid | 1-stearoyl-GPG (18:0) | | 1.1 | 0.8316 | 1 | 0.9933 |
| Lipid | Lysophospholipid | 1-oleoyl-GPG (18:1)* | | 1.21 | 0.5622 | 0.98 | 0.9417 |
| Lipid | Lysophospholipid | 1-linoleoyl-GPG (18:2)* | | 0.98 | 0.9236 | 1.08 | 0.7553 |
| Lipid | Lysophospholipid | 1-palmitoyl-GPI (16:0) | HMDB61695 | 1.41 | 0.4974 | 0.65 | 0.4037 |
| Lipid | Lysophospholipid | 1-stearoyl-GPI (18:0) | HMDB61696 | 1.09 | 0.8863 | 0.67 | 0.5095 |
| Lipid | Lysophospholipid | 1-oleoyl-GPI (18:1)* | | 0.91 | 0.7468 | 0.99 | 0.9736 |
| Lipid | Lysophospholipid | 1-linoleoyl-GPI (18:2)* | | **1.36** | 0.0724 | 0.99 | 0.9419 |
| Lipid | Glycolipid Metabolism | galactosylglycerol* | HMDB06790 | 1.42 | 0.5564 | 0.77 | 0.6726 |
| Lipid | Glycolipid Metabolism | 1-palmitoyl-2-linoleoyl-digalactosylglycerol (16:0/18:2)* | | 0.72 | 0.3390 | 1.46 | 0.2829 |
| Lipid | Glycolipid Metabolism | 1-palmitoyl-2-linoleoyl-galactosylglycerol (16:0/18:2)* | | 1.07 | 0.8597 | 1.43 | 0.3687 |
| Lipid | Glycolipid Metabolism | 1-palmitoyl-2-linolenoyl-galactosylglycerol (16:0/18:3)* | | 1.28 | 0.2591 | 0.99 | 0.9737 |
| Lipid | Glycolipid Metabolism | 1,2-dilinoleoyl-digalactosylglycerol (18:2/18:2)* | | 0.77 | 0.4201 | 0.94 | 0.8551 |
| Lipid | Glycolipid Metabolism | 1,2-dilinoleoyl-galactosylglycerol (18:2/18:2)* | | 1.43 | 0.2438 | 1.27 | 0.4606 |
| Lipid | Glycolipid Metabolism | 1-linoleoyl-2-linolenoyl-galactosylglycerol (18:2/18:3)* | | 0.93 | 0.7558 | 1.09 | 0.7050 |
| Lipid | Glycolipid Metabolism | 1-linoleoyl-2-linolenoyl-digalactosylglycerol (18:2/18:3)* | | 0.97 | 0.8943 | 1.01 | 0.9561 |
| Lipid | Glycolipid Metabolism | 1,2-dilinolenoyl-galactosylglycerol (18:3/18:3)* | | 0.91 | 0.7955 | 0.7 | 0.3216 |
| Lipid | Lysoplasmalogen | 1-(1-enyl-palmitoyl)-GPC (P-16:0)* | HMDB10407 | 0.8 | 0.3343 | 1.03 | 0.8877 |
| Lipid | Lysoplasmalogen | 1-(1-enyl-palmitoyl)-GPE (P-16:0)* | | 0.95 | 0.8687 | 1.13 | 0.6952 |
| Lipid | Lysoplasmalogen | 1-(1-enyl-oleoyl)-GPE (P-18:1)* | | 1.15 | 0.6979 | 1.28 | 0.5233 |
| Lipid | Lysoplasmalogen | 1-(1-enyl-stearoyl)-GPE (P- | | 0.83 | 0.496 | 1.14 | 0.621 |

| | | | | | | | |
|---|---|---|---|---|---|---|---|
| | | 18:0)* | | | 4 | | 2 |
| Lipid | Glycerolipid Metabolism | glycerol | HMDB00131 | 1.08 | 0.8403 | 0.96 | 0.9258 |
| Lipid | Glycerolipid Metabolism | glycerol 3-phosphate | HMDB00126 | 1.04 | 0.9245 | 0.8 | 0.5743 |
| Lipid | Glycerolipid Metabolism | glycerophosphoglycerol | | 0.97 | 0.9462 | 0.91 | 0.8620 |
| Lipid | Monoacylglycerol | 1-myristoylglycerol (14:0) | HMDB11561 | 1.08 | 0.8363 | **0.51** | 0.0894 |
| Lipid | Monoacylglycerol | 1-pentadecanoylglycerol (15:0) | | 1.54 | 0.1293 | 1.02 | 0.9447 |
| Lipid | Monoacylglycerol | 1-palmitoylglycerol (16:0) | HMDB31074 | 1.21 | 0.5590 | 1.21 | 0.5765 |
| Lipid | Monoacylglycerol | 1-palmitoleoylglycerol (16:1)* | HMDB11565 | 1.02 | 0.9362 | 0.79 | 0.3335 |
| Lipid | Monoacylglycerol | 1-oleoylglycerol (18:1) | HMDB11567 | 1.33 | 0.5450 | 1.64 | 0.3102 |
| Lipid | Monoacylglycerol | 1-linoleoylglycerol (18:2) | | 1.6 | 0.3964 | 2.35 | 0.1443 |
| Lipid | Monoacylglycerol | 1-linolenoylglycerol (18:3) | HMDB11569 | **2.36** | 0.0457 | 1.01 | 0.9871 |
| Lipid | Monoacylglycerol | 2-palmitoylglycerol (16:0) | HMDB11533 | 1.02 | 0.9524 | 1.22 | 0.6155 |
| Lipid | Monoacylglycerol | 2-oleoylglycerol (18:1) | HMDB11537 | 1.01 | 0.9848 | 0.96 | 0.9377 |
| Lipid | Monoacylglycerol | 2-linoleoylglycerol (18:2) | HMDB11538 | 1.01 | 0.9779 | 1.75 | 0.2884 |
| Lipid | Diacylglycerol | palmitoyl-oleoyl-glycerol (16:0/18:1) [2]* | HMDB07102 | **1.3** | 0.0912 | 0.87 | 0.4012 |
| Lipid | Diacylglycerol | palmitoyl-linoleoyl-glycerol (16:0/18:2) [1]* | HMDB07103 | 1.19 | 0.4401 | 1.12 | 0.6269 |
| Lipid | Diacylglycerol | palmitoyl-linoleoyl-glycerol (16:0/18:2) [2]* | HMDB07103 | 1.62 | 0.1274 | 1.19 | 0.6012 |
| Lipid | Diacylglycerol | oleoyl-oleoyl-glycerol (18:1/18:1) [2]* | HMDB07218 | **1.55** | 0.0901 | 0.71 | 0.2113 |
| Lipid | Diacylglycerol | oleoyl-linoleoyl-glycerol (18:1/18:2) [1] | HMDB07219 | 1.54 | 0.3074 | **2.21** | 0.0778 |
| Lipid | Diacylglycerol | oleoyl-linoleoyl-glycerol (18:1/18:2) [2] | HMDB07219 | **2.43** | 0.0932 | 2.1 | 0.1811 |
| Lipid | Diacylglycerol | linoleoyl-linoleoyl-glycerol (18:2/18:2) [1]* | HMDB07248 | 2.42 | 0.1319 | 2.12 | 0.2236 |
| Lipid | Diacylglycerol | linoleoyl-linoleoyl-glycerol (18:2/18:2) [2]* | HMDB07248 | 1.9 | 0.1856 | 1.35 | 0.5510 |
| Lipid | Diacylglycerol | linoleoyl-linolenoyl-glycerol (18:2/18:3) [1]* | HMDB07249 | **2.04** | 0.0285 | 1.18 | 0.6201 |
| Lipid | Diacylglycerol | linoleoyl-linolenoyl-glycerol (18:2/18:3) [2]* | HMDB07250 | **2.33** | 0.0665 | 1.31 | 0.5719 |
| Lipid | Diacylglycerol | linolenoyl-linolenoyl-glycerol (18:3/18:3) [1]* | HMDB07278 | 1.31 | 0.1037 | 0.93 | 0.7007 |
| Lipid | Diacylglycerol | linolenoyl-linolenoyl-glycerol (18:3/18:3) [2]* | HMDB07278 | **2.29** | 0.0404 | 0.76 | 0.5121 |
| Lipid | Diacylglycerol | oleoyl-arachidonoyl-glycerol (18:1/20:4) [2]* | HMDB07228 | 1.21 | 0.1904 | 0.82 | 0.1850 |
| Lipid | Diacylglycerol | linoleoyl-arachidonoyl-glycerol (18:2/20:4) [1]* | HMDB07257 | 1.27 | 0.1361 | **0.73** | 0.0596 |
| Lipid | Diacylglycerol | linoleoyl-arachidonoyl-glycerol (18:2/20:4) [2]* | HMDB07257 | 1.22 | 0.4137 | 0.73 | 0.2139 |
| Lipid | Diacylglycerol | linoleoyl-docosahexaenoyl-glycerol (18:2/22:6) [1]* | | 1.03 | 0.8045 | **0.66** | 0.0006 |
| Lipid | Diacylglycerol | linoleoyl-docosahexaenoyl-glycerol (18:2/22:6) [2]* | HMDB07266 | 0.98 | 0.9291 | **0.59** | 0.0165 |
| Lipid | Sphingolipid Metabolism | sphinganine | HMDB00269 | 0.92 | 0.7653 | 0.84 | 0.5424 |
| Lipid | Sphingolipid Metabolism | 3-ketosphinganine | HMDB01480 | 1.9 | 0.1885 | 0.63 | 0.3693 |
| Lipid | Sphingolipid Metabolism | N-palmitoyl-sphinganine (d18:0/16:0) | HMDB11760 | 1.04 | 0.8507 | 0.87 | 0.5506 |

| | | | | | | | |
|---|---|---|---|---|---|---|---|
| Lipid | Sphingolipid Metabolism | N-behenoyl-sphingadienine (d18:2/22:0)* | | 0.39 | 0.125 1 | 1.57 | 0.480 1 |
| Lipid | Sphingolipid Metabolism | galactosylsphingosine | HMDB006 48 | 0.59 | 0.261 7 | 1.3 | 0.580 1 |
| Lipid | Sphingolipid Metabolism | N-butyroyl-sphingosine (d18:1/4:0) | | 1.16 | 0.731 8 | 1.85 | 0.174 3 |
| Lipid | Sphingolipid Metabolism | myristoyl dihydrosphingomyelin (d18:0/14:0)* | HMDB120 85 | 0.73 | 0.466 8 | 0.64 | 0.324 8 |
| Lipid | Sphingolipid Metabolism | palmitoyl dihydrosphingomyelin (d18:0/16:0)* | | 0.67 | 0.231 9 | 0.9 | 0.748 0 |
| Lipid | Sphingolipid Metabolism | behenoyl dihydrosphingomyelin (d18:0/22:0)* | HMDB120 91 | 0.52 | 0.149 2 | 1.21 | 0.686 9 |
| Lipid | Sphingolipid Metabolism | palmitoyl sphingomyelin (d18:1/16:0) | | 0.63 | 0.326 2 | 0.77 | 0.575 9 |
| Lipid | Sphingolipid Metabolism | stearoyl sphingomyelin (d18:1/18:0) | HMDB013 48 | 0.45 | 0.152 2 | 0.84 | 0.760 6 |
| Lipid | Sphingolipid Metabolism | behenoyl sphingomyelin (d18:1/22:0)* | HMDB121 03 | 0.4 | 0.117 1 | 0.71 | 0.565 9 |
| Lipid | Sphingolipid Metabolism | tricosanoyl sphingomyelin (d18:1/23:0)* | HMDB121 05 | 0.58 | 0.278 6 | 1.21 | 0.710 8 |
| Lipid | Sphingolipid Metabolism | lignoceroyl sphingomyelin (d18:1/24:0) | | 0.65 | 0.419 7 | 0.83 | 0.736 5 |
| Lipid | Sphingolipid Metabolism | sphingomyelin (d18:1/14:0, d16:1/16:0)* | HMDB120 97 | 0.61 | 0.312 9 | 0.83 | 0.701 9 |
| Lipid | Sphingolipid Metabolism | sphingomyelin (d17:1/16:0, d18:1/15:0, d16:1/17:0)* | | 0.82 | 0.638 1 | **2.17** | 0.077 9 |
| Lipid | Sphingolipid Metabolism | sphingomyelin (d18:2/16:0, d18:1/16:1)* | | 0.77 | 0.447 1 | 1.01 | 0.969 3 |
| Lipid | Sphingolipid Metabolism | sphingomyelin (d18:1/17:0, d17:1/18:0, d19:1/16:0) | | 0.77 | 0.500 9 | 1.61 | 0.224 7 |
| Lipid | Sphingolipid Metabolism | sphingomyelin (d18:1/18:1, d18:2/18:0) | HMDB121 01 | 0.66 | 0.161 7 | 1.07 | 0.818 2 |
| Lipid | Sphingolipid Metabolism | sphingomyelin (d18:1/20:0, d16:1/22:0)* | HMDB121 02 | 0.55 | 0.281 4 | 0.76 | 0.629 5 |
| Lipid | Sphingolipid Metabolism | sphingomyelin (d18:1/20:1, d18:2/20:0)* | | **0.62** | 0.093 0 | 0.77 | 0.364 0 |
| Lipid | Sphingolipid Metabolism | sphingomyelin (d18:1/21:0, d17:1/22:0, d16:1/23:0)* | | 0.59 | 0.107 8 | 0.86 | 0.652 2 |
| Lipid | Sphingolipid Metabolism | sphingomyelin (d18:1/22:1, d18:2/22:0, d16:1/24:1)* | HMDB121 04 | 0.76 | 0.547 4 | 1.02 | 0.969 4 |
| Lipid | Sphingolipid Metabolism | sphingomyelin (d18:2/23:0, d18:1/23:1, d17:1/24:1)* | | 0.73 | 0.127 6 | 0.94 | 0.757 9 |
| Lipid | Sphingolipid Metabolism | sphingomyelin (d18:1/24:1, d18:2/24:0)* | HMDB121 07 | 0.49 | 0.233 8 | 0.86 | 0.797 7 |
| Lipid | Sphingolipid Metabolism | sphingomyelin (d18:2/24:1, d18:1/24:2)* | | 0.87 | 0.766 5 | 0.92 | 0.862 4 |
| Lipid | Sphingolipid Metabolism | sphingosine | HMDB002 52 | 0.81 | 0.343 7 | 1 | 0.988 2 |
| Lipid | Sphingolipid Metabolism | N-acetylsphingosine | HMDB049 50 | 1.11 | 0.798 7 | **2.44** | 0.038 7 |
| Lipid | Sphingolipid Metabolism | phytosphingosine | HMDB046 10 | 0.78 | 0.263 0 | **1.51** | 0.065 0 |
| Lipid | Sphingolipid Metabolism | sphingomyelin (d18:0/20:0, d16:0/22:0)* | | 0.7 | 0.296 9 | 0.74 | 0.377 0 |
| Lipid | Sphingolipid Metabolism | sphingomyelin (d18:0/18:0, d19:0/17:0)* | HMDB120 87 | **0.48** | 0.075 1 | 0.87 | 0.738 1 |
| Lipid | Sphingolipid Metabolism | sphingomyelin (d18:1/19:0, d19:1/18:0)* | | 0.73 | 0.157 0 | 0.9 | 0.640 3 |
| Lipid | Sphingolipid Metabolism | heptadecasphingosine (d17:1) | | 0.76 | 0.422 1 | 1.07 | 0.841 6 |
| Lipid | Sphingolipid Metabolism | hexadecasphingosine (d16:1)* | | 0.86 | 0.648 1 | 1.08 | 0.813 5 |
| Lipid | Sphingolipid Metabolism | N-palmitoyl-heptadecasphingosine (d17:1/16:0)* | | 1.02 | 0.947 4 | 0.88 | 0.639 7 |

| | | | | | | | |
|---|---|---|---|---|---|---|---|
| Lipid | Sphingolipid Metabolism | N-stearoyl-sphinganine (d18:0/18:0)* | | 0.95 | 0.8354 | 0.89 | 0.6923 |
| Lipid | Sphingolipid Metabolism | lactosyl-N-behenoyl-sphingosine (d18:1/22:0)* | | 0.91 | 0.8299 | 0.7 | 0.4428 |
| Lipid | Sphingolipid Metabolism | hexadecasphinganine (d16:0)* | | 1.16 | 0.7215 | 0.7 | 0.4117 |
| Lipid | Sphingolipid Metabolism | lactosyl-N-arachidoyl-sphingosine (d18:1/20:0)* | | 0.77 | 0.5216 | 0.81 | 0.5996 |
| Lipid | Sphingolipid Metabolism | N-(2-hydroxypalmitoyl)-sphingosine (d18:1/16:0(2OH)) | | 0.98 | 0.9540 | 1.11 | 0.7528 |
| Lipid | Sphingolipid Metabolism | N-oleoyl-sphingosine (d18:1/18:1)* | HMDB04948 | 0.66 | 0.2016 | 1.07 | 0.8468 |
| Lipid | Sphingolipid Metabolism | eicosanoylsphingosine (d20:1)* | | 0.87 | 0.5485 | 0.98 | 0.9359 |
| Lipid | Ceramides | N-palmitoyl-sphingosine (d18:1/16:0) | HMDB04949 | 1 | 0.9973 | 0.82 | 0.3655 |
| Lipid | Ceramides | N-stearoyl-sphingosine (d18:1/18:0)* | HMDB04950 | 0.79 | 0.3811 | 0.89 | 0.6705 |
| Lipid | Ceramides | ceramide (d16:1/24:1, d18:1/22:1)* | | 1.03 | 0.9520 | 0.54 | 0.2239 |
| Lipid | Ceramides | ceramide (d18:1/14:0, d16:1/16:0)* | | 1.03 | 0.9195 | 0.76 | 0.2937 |
| Lipid | Ceramides | ceramide (d18:1/17:0, d17:1/18:0)* | | 1 | 0.9929 | 0.89 | 0.6459 |
| Lipid | Ceramides | ceramide (d18:2/24:1, d18:1/24:2)* | | 0.93 | 0.8008 | 0.82 | 0.5114 |
| Lipid | Ceramides | glycosyl-N-palmitoyl-sphingosine (d18:1/16:0) | | 0.88 | 0.6698 | 0.95 | 0.8727 |
| Lipid | Ceramides | glycosyl-N-stearoyl-sphingosine (d18:1/18:0) | | 0.58 | 0.1520 | 1.26 | 0.5561 |
| Lipid | Ceramides | glycosyl-N-behenoyl-sphingadienine (d18:2/22:0)* | | 0.6 | 0.2354 | 1.18 | 0.7004 |
| Lipid | Ceramides | glycosyl-N-(2-hydroxynervonoyl)-sphingosine (d18:1/24:1(2OH))* | | 0.91 | 0.7439 | 1.48 | 0.1962 |
| Lipid | Ceramides | lactosyl-N-palmitoyl-sphingosine (d18:1/16:0) | | 0.9 | 0.7968 | 0.64 | 0.2856 |
| Lipid | Ceramides | lactosyl-N-stearoyl-sphingosine (d18:1/18:0)* | HMDB11591 | 0.58 | 0.2668 | 0.73 | 0.5342 |
| Lipid | Ceramides | lactosyl-N-nervonoyl-sphingosine (d18:1/24:1)* | | 0.92 | 0.8640 | 0.5 | 0.1555 |
| Lipid | Ceramides | glycosyl ceramide (d16:1/24:1, d18:1/22:1)* | | 0.76 | 0.4292 | 0.72 | 0.3617 |
| Lipid | Ceramides | glycosyl ceramide (d18:1/20:0, d16:1/22:0)* | | 0.65 | 0.2589 | 1.27 | 0.5299 |
| Lipid | Ceramides | glycosyl ceramide (d18:2/24:1, d18:1/24:2)* | | 0.75 | 0.4919 | 0.92 | 0.8346 |
| Lipid | Mevalonate Metabolism | 3-hydroxy-3-methylglutarate | HMDB00355 | 0.74 | 0.4206 | **2.77** | 0.0098 |
| Lipid | Mevalonate Metabolism | mevalonate | HMDB00227 | 1.46 | 0.3525 | 0.54 | 0.1372 |
| Lipid | Mevalonate Metabolism | mevalonolactone | HMDB06024 | 1.08 | 0.7964 | 0.65 | 0.1611 |
| Lipid | Sterol | lanosterol | HMDB01251 | 0.74 | 0.3366 | 1.19 | 0.5842 |
| Lipid | Sterol | desmosterol | HMDB02719 | **0.48** | 0.0780 | 1 | 0.9945 |
| Lipid | Sterol | cholesterol | HMDB00067 | 0.73 | 0.2720 | 1.03 | 0.9326 |
| Lipid | Sterol | 3beta-hydroxy-5-cholestenoate | | 1.13 | 0.7122 | 1.05 | 0.8872 |
| Lipid | Sterol | coprostanol | HMDB00577 | 1.08 | 0.7510 | 0.76 | 0.2509 |
| Lipid | Sterol | 4-cholesten-3-one | HMDB00921 | 0.71 | 0.3356 | 0.79 | 0.5203 |
| Lipid | Sterol | beta-sitosterol | HMDB00852 | 0.77 | 0.369 | **2.89** | 0.000 |

| | | | | | | | |
|---|---|---|---|---|---|---|---|
| | | | | | 9 | | 6 |
| Lipid | Sterol | stigmasterol | HMDB00937 | 1.14 | 0.6191 | **2.01** | 0.0159 |
| Lipid | Sterol | campesterol | HMDB02869 | 0.86 | 0.6689 | **2.1** | 0.0415 |
| Lipid | Sterol | 7-hydroxycholesterol (alpha or beta) | HMDB06119 | 0.81 | 0.4562 | 0.85 | 0.5864 |
| Lipid | Pregnenolone Steroids | pregnenolone sulfate | HMDB00774 | 0.8 | 0.5656 | 0.86 | 0.7158 |
| Lipid | Pregnenolone Steroids | 17alpha-hydroxypregnenolone 3-sulfate | HMDB00416 | 1.15 | 0.6368 | 0.67 | 0.1965 |
| Lipid | Pregnenolone Steroids | 21-hydroxypregnenolone monosulfate (1) | | 0.78 | 0.1765 | 0.76 | 0.1549 |
| Lipid | Pregnenolone Steroids | 21-hydroxypregnenolone disulfate | | 0.79 | 0.6159 | 1.12 | 0.8150 |
| Lipid | Pregnenolone Steroids | 21-hydroxypregnanolone disulfate | | 0.91 | 0.8199 | 1.47 | 0.3710 |
| Lipid | Pregnenolone Steroids | pregnenediol disulfate (C21H34O8S2)* | | 0.75 | 0.6166 | 0.88 | 0.8293 |
| Lipid | Pregnenolone Steroids | pregnenediol sulfate (C21H34O5S)* | | 0.96 | 0.9322 | 0.87 | 0.7832 |
| Lipid | Progestin Steroids | 5alpha-pregnan-3beta-ol,20-one sulfate | | 1.36 | 0.2800 | 0.91 | 0.7620 |
| Lipid | Progestin Steroids | 5alpha-pregnan-3beta,20beta-diol monosulfate (1) | | 0.56 | 0.1447 | 0.59 | 0.1764 |
| Lipid | Progestin Steroids | 5alpha-pregnan-3beta,20alpha-diol monosulfate (2) | | 0.89 | 0.6558 | 0.79 | 0.3978 |
| Lipid | Progestin Steroids | 5alpha-pregnan-3beta,20alpha-diol disulfate | | 0.67 | 0.2903 | 0.83 | 0.6302 |
| Lipid | Progestin Steroids | 5alpha-pregnan-diol disulfate | | 0.61 | 0.2676 | 0.57 | 0.2174 |
| Lipid | Progestin Steroids | pregnanolone/allopregnanolone sulfate | | **0.38** | 0.0640 | **0.36** | 0.0564 |
| Lipid | Corticosteroids | tetrahydrocorticosterone | HMDB00268 | 0.65 | 0.1965 | 0.71 | 0.3195 |
| Lipid | Corticosteroids | cortisone 21-sulfate | HMDB02802 | 0.74 | 0.1949 | 0.81 | 0.3731 |
| Lipid | Corticosteroids | cortolone | HMDB03128 | 0.81 | 0.4222 | 0.96 | 0.8866 |
| Lipid | Androgenic Steroids | 11-ketoetiocholanolone sulfate | | **0.57** | 0.0625 | 0.61 | 0.1135 |
| Lipid | Androgenic Steroids | dehydroisoandrosterone sulfate (DHEA-S) | HMDB01032 | 0.88 | 0.7183 | 0.83 | 0.5930 |
| Lipid | Androgenic Steroids | 16a-hydroxy DHEA 3-sulfate | | 0.93 | 0.8511 | 0.83 | 0.6574 |
| Lipid | Androgenic Steroids | androsterone sulfate | HMDB02759 | 0.72 | 0.2540 | 0.75 | 0.3111 |
| Lipid | Androgenic Steroids | 5alpha-androstan-3alpha,17alpha-diol monosulfate | | 0.92 | 0.7385 | 0.77 | 0.2800 |
| Lipid | Androgenic Steroids | 5alpha-androstan-3alpha,17alpha-diol disulfate | | **0.56** | 0.0199 | **0.63** | 0.0625 |
| Lipid | Androgenic Steroids | androstenediol (3beta,17beta) monosulfate (1) | HMDB03818 | 0.95 | 0.8934 | 0.88 | 0.7215 |
| Lipid | Androgenic Steroids | androstenediol (3beta,17beta) disulfate (1) | HMDB03818 | 0.83 | 0.5938 | 0.78 | 0.4899 |
| Lipid | Androgenic Steroids | androstenediol (3beta,17beta) disulfate (2) | HMDB03818 | 0.71 | 0.2185 | **0.54** | 0.0324 |
| Lipid | Androgenic Steroids | androstenediol (3alpha, 17alpha) monosulfate (2) | | 0.74 | 0.1809 | 0.91 | 0.6905 |
| Lipid | Androgenic Steroids | 5alpha-androstan-3alpha,17beta-diol monosulfate (1) | | 0.78 | 0.1059 | 1.06 | 0.7224 |
| Lipid | Androgenic Steroids | andro steroid monosulfate C19H28O6S (1)* | HMDB02759 | 0.92 | 0.8125 | 0.63 | 0.1893 |
| Lipid | Primary Bile Acid Metabolism | cholate | HMDB00619 | 0.66 | 0.3826 | 2.1 | 0.1315 |

| | | | | | | | |
|---|---|---|---|---|---|---|---|
| Lipid | Primary Bile Acid Metabolism | glycocholate | HMDB001<br>38 | **0.39** | 0.048<br>3 | 1.37 | 0.523<br>7 |
| Lipid | Primary Bile Acid Metabolism | taurocholate | HMDB000<br>36 | 0.46 | 0.160<br>7 | 1.03 | 0.955<br>5 |
| Lipid | Primary Bile Acid Metabolism | chenodeoxycholate | HMDB005<br>18 | 0.66 | 0.420<br>3 | 2.37 | 0.110<br>6 |
| Lipid | Primary Bile Acid Metabolism | glycochenodeoxycholate | HMDB006<br>37 | **0.43** | 0.043<br>4 | 0.64 | 0.305<br>2 |
| Lipid | Primary Bile Acid Metabolism | taurochenodeoxycholate | HMDB009<br>51 | **0.39** | 0.072<br>3 | 1.15 | 0.797<br>8 |
| Lipid | Primary Bile Acid Metabolism | cholate sulfate | | **3.53** | 0.082<br>6 | 0.3 | 0.115<br>1 |
| Lipid | Primary Bile Acid Metabolism | glycochenodeoxycholate glucuronide (2) | | **0.37** | 0.019<br>5 | 0.78 | 0.571<br>7 |
| Lipid | Primary Bile Acid Metabolism | glycochenodeoxycholate sulfate | | 0.57 | 0.228<br>3 | **0.35** | 0.028<br>4 |
| Lipid | Primary Bile Acid Metabolism | glycocholate sulfate | | 0.79 | 0.374<br>6 | 1.09 | 0.754<br>5 |
| Lipid | Secondary Bile Acid Metabolism | deoxycholate | HMDB006<br>26 | 0.54 | 0.227<br>4 | 0.58 | 0.290<br>7 |
| Lipid | Secondary Bile Acid Metabolism | deoxycholic acid sulfate | | 0.86 | 0.746<br>7 | 0.97 | 0.950<br>0 |
| Lipid | Secondary Bile Acid Metabolism | glycodeoxycholate | HMDB006<br>31 | 0.89 | 0.647<br>6 | 0.99 | 0.974<br>2 |
| Lipid | Secondary Bile Acid Metabolism | lithocholate | HMDB007<br>61 | 0.38 | 0.277<br>0 | 0.37 | 0.265<br>4 |
| Lipid | Secondary Bile Acid Metabolism | taurolithocholate | HMDB007<br>22 | 0.8 | 0.108<br>8 | 0.95 | 0.705<br>5 |
| Lipid | Secondary Bile Acid Metabolism | taurolithocholate 3-sulfate | HMDB025<br>80 | 0.53 | 0.156<br>8 | 1.18 | 0.719<br>3 |
| Lipid | Secondary Bile Acid Metabolism | ursodeoxycholate | HMDB009<br>46 | 1.04 | 0.933<br>5 | 2.12 | 0.119<br>7 |
| Lipid | Secondary Bile Acid Metabolism | isoursodeoxycholate | HMDB006<br>86 | 0.97 | 0.942<br>0 | 1.36 | 0.520<br>4 |
| Lipid | Secondary Bile Acid Metabolism | glycoursodeoxycholate | HMDB007<br>08 | 0.63 | 0.324<br>0 | 0.76 | 0.566<br>2 |
| Lipid | Secondary Bile Acid Metabolism | tauroursodeoxycholate | HMDB008<br>74 | 0.76 | 0.533<br>7 | 1.01 | 0.988<br>8 |
| Lipid | Secondary Bile Acid Metabolism | dehydrolithocholate | HMDB005<br>02 | 0.73 | 0.604<br>5 | 0.58 | 0.374<br>0 |
| Lipid | Secondary Bile Acid Metabolism | 7,12-diketolithocholate | | 0.82 | 0.712<br>4 | 0.95 | 0.933<br>7 |
| Lipid | Secondary Bile Acid Metabolism | 6-oxolithocholate | | 0.72 | 0.619<br>8 | 0.77 | 0.693<br>7 |
| Lipid | Secondary Bile Acid Metabolism | 7-ketolithocholate | HMDB004<br>67 | 0.88 | 0.768<br>9 | 1.41 | 0.455<br>0 |
| Lipid | Secondary Bile Acid Metabolism | hyocholate | HMDB007<br>60 | 0.68 | 0.282<br>6 | 1.59 | 0.206<br>3 |
| Lipid | Secondary Bile Acid Metabolism | dehydrocholic acid | | 0.96 | 0.918<br>5 | 0.63 | 0.304<br>5 |
| Lipid | Secondary Bile Acid Metabolism | 3-dehydrocholate | HMDB005<br>02 | 0.49 | 0.194<br>9 | **2.84** | 0.065<br>6 |
| Lipid | Secondary Bile Acid Metabolism | 12-dehydrocholate | HMDB004<br>00 | **0.39** | 0.050<br>8 | 2.25 | 0.112<br>1 |
| Lipid | Secondary Bile Acid Metabolism | glycocholenate sulfate* | | 0.79 | 0.698<br>8 | 1.28 | 0.702<br>4 |
| Lipid | Secondary Bile Acid Metabolism | taurocholenate sulfate | | 0.62 | 0.485<br>6 | 2.2 | 0.263<br>9 |
| Lipid | Secondary Bile Acid Metabolism | 7-ketodeoxycholate | HMDB003<br>91 | 0.81 | 0.653<br>6 | 1.5 | 0.402<br>6 |
| Lipid | Secondary Bile Acid Metabolism | 7alpha-hydroxycholestenone | HMDB019<br>93 | **0.64** | 0.041<br>1 | 0.88 | 0.571<br>6 |
| Lipid | Secondary Bile Acid Metabolism | 3b-hydroxy-5-cholenoic acid | HMDB003<br>08 | 1.11 | 0.713<br>9 | 0.78 | 0.401<br>8 |
| Lipid | Secondary Bile Acid Metabolism | ursodeoxycholate sulfate (1) | | 0.98 | 0.976<br>3 | 0.37 | 0.158<br>7 |
| Lipid | Secondary Bile Acid Metabolism | ursocholate | | 1.17 | 0.786<br>7 | 2.3 | 0.165<br>2 |

| | | | | | | | |
|---|---|---|---|---|---|---|---|
| Nucleotide | Purine Metabolism, (Hypo)Xanthine/Inosine containing | inosine | HMDB00195 | 0.94 | 0.9058 | 0.84 | 0.7682 |
| Nucleotide | Purine Metabolism, (Hypo)Xanthine/Inosine containing | hypoxanthine | HMDB00157 | 0.73 | 0.2458 | 1.22 | 0.4738 |
| Nucleotide | Purine Metabolism, (Hypo)Xanthine/Inosine containing | xanthine | HMDB00292 | 0.9 | 0.5782 | 0.94 | 0.7652 |
| Nucleotide | Purine Metabolism, (Hypo)Xanthine/Inosine containing | xanthosine | HMDB00299 | 0.68 | 0.3536 | 1.46 | 0.3641 |
| Nucleotide | Purine Metabolism, (Hypo)Xanthine/Inosine containing | 2'-deoxyinosine | HMDB00071 | 0.53 | 0.3082 | 2.76 | 0.1210 |
| Nucleotide | Purine Metabolism, (Hypo)Xanthine/Inosine containing | urate | HMDB00289 | 0.78 | 0.6366 | 1.2 | 0.7409 |
| Nucleotide | Purine Metabolism, (Hypo)Xanthine/Inosine containing | allantoin | HMDB00462 | 0.64 | 0.2900 | 1.49 | 0.3523 |
| Nucleotide | Purine Metabolism, (Hypo)Xanthine/Inosine containing | allantoic acid | HMDB01209 | 0.66 | 0.2929 | 1.07 | 0.8709 |
| Nucleotide | Purine Metabolism, Adenine containing | adenosine | HMDB00050 | 0.98 | 0.9572 | 1.51 | 0.2550 |
| Nucleotide | Purine Metabolism, Adenine containing | adenine | HMDB00034 | 0.86 | 0.6977 | 1.68 | 0.2080 |
| Nucleotide | Purine Metabolism, Adenine containing | 1-methyladenine | HMDB11599 | 1.05 | 0.8425 | 0.81 | 0.4123 |
| Nucleotide | Purine Metabolism, Adenine containing | N6-dimethylallyladenine | | 0.94 | 0.8249 | **0.28** | 0.0000 |
| Nucleotide | Purine Metabolism, Adenine containing | N6-carbamoylthreonyladenosine | HMDB41623 | 1.07 | 0.7292 | 1 | 1.0000 |
| Nucleotide | Purine Metabolism, Adenine containing | 2'-deoxyadenosine | HMDB00101 | 1.19 | 0.7249 | 2.21 | 0.1266 |
| Nucleotide | Purine Metabolism, Guanine containing | guanosine-2',3'-cyclic monophosphate | HMDB11629 | 1.03 | 0.8840 | **1.51** | 0.0655 |
| Nucleotide | Purine Metabolism, Guanine containing | guanosine | HMDB00133 | 0.81 | 0.7435 | **3.02** | 0.0886 |
| Nucleotide | Purine Metabolism, Guanine containing | guanine | HMDB00132 | **0.47** | 0.0876 | **3.22** | 0.0112 |
| Nucleotide | Purine Metabolism, Guanine containing | 1-methylguanine | HMDB03282 | **0.54** | 0.0748 | 1.51 | 0.2487 |
| Nucleotide | Purine Metabolism, Guanine containing | 7-methylguanine | HMDB00897 | 0.9 | 0.7969 | 1.55 | 0.2797 |
| Nucleotide | Purine Metabolism, Guanine containing | 2'-O-methylguanosine | | 0.84 | 0.2864 | 0.97 | 0.8539 |
| Nucleotide | Purine Metabolism, Guanine containing | N2-methylguanosine | HMDB05862 | 0.82 | 0.2140 | 1 | 1.0000 |
| Nucleotide | Purine Metabolism, Guanine containing | 8-hydroxyguanine | HMDB02032 | 0.7 | 0.5426 | 1.75 | 0.3501 |
| Nucleotide | Purine Metabolism, Guanine containing | 8-hydroxy-2'-deoxyguanosine | HMDB03333 | 0.96 | 0.9033 | 0.99 | 0.9820 |
| Nucleotide | Purine Metabolism, Guanine containing | 2'-deoxyguanosine | HMDB00085 | 0.64 | 0.2531 | 1.25 | 0.5809 |
| Nucleotide | Pyrimidine Metabolism, Orotate containing | N-carbamoylaspartate | HMDB00828 | 1.14 | 0.5814 | **0.59** | 0.0326 |
| Nucleotide | Pyrimidine Metabolism, Orotate containing | dihydroorotate | HMDB03349 | 1.89 | 0.2513 | 1.4 | 0.5595 |
| Nucleotide | Pyrimidine Metabolism, Orotate containing | orotate | HMDB00226 | 1.27 | 0.4799 | 0.93 | 0.8333 |
| Nucleotide | Pyrimidine Metabolism, Orotate containing | orotidine | HMDB00788 | 0.55 | 0.4108 | 2.98 | 0.1450 |

| | | | | | | | |
|---|---|---|---|---|---|---|---|
| Nucleotide | Pyrimidine Metabolism, Uracil containing | uridine-2',3'-cyclic monophosphate | HMDB11640 | 1.27 | 0.3081 | 1.75 | 0.0243 |
| Nucleotide | Pyrimidine Metabolism, Uracil containing | uridine | HMDB00296 | 1.13 | 0.7916 | 1.93 | 0.1647 |
| Nucleotide | Pyrimidine Metabolism, Uracil containing | uracil | HMDB00300 | 0.96 | 0.8453 | 0.9 | 0.6448 |
| Nucleotide | Pyrimidine Metabolism, Uracil containing | pseudouridine | HMDB00767 | 0.81 | 0.4740 | 0.87 | 0.6483 |
| Nucleotide | Pyrimidine Metabolism, Uracil containing | 2'-O-methyluridine | | 0.69 | 0.1875 | 0.9 | 0.7055 |
| Nucleotide | Pyrimidine Metabolism, Uracil containing | 5-methyluridine (ribothymidine) | HMDB00884 | 0.84 | 0.6936 | 1.07 | 0.8784 |
| Nucleotide | Pyrimidine Metabolism, Uracil containing | 5,6-dihydrouracil | HMDB00076 | 1.05 | 0.9167 | 1.39 | 0.5116 |
| Nucleotide | Pyrimidine Metabolism, Uracil containing | 2'-deoxyuridine | HMDB00012 | 1 | 0.9899 | 0.75 | 0.4019 |
| Nucleotide | Pyrimidine Metabolism, Uracil containing | 4-ureidobutyrate | | 0.6 | 0.2206 | 1.07 | 0.8824 |
| Nucleotide | Pyrimidine Metabolism, Uracil containing | 3-ureidoisobutyrate | HMDB02031 | 0.98 | 0.9574 | 0.54 | 0.1939 |
| Nucleotide | Pyrimidine Metabolism, Uracil containing | 3-ureidopropionate | HMDB00026 | 1.14 | 0.7789 | 0.64 | 0.3526 |
| Nucleotide | Pyrimidine Metabolism, Uracil containing | beta-alanine | HMDB00056 | 1.26 | 0.5585 | 0.94 | 0.8729 |
| Nucleotide | Pyrimidine Metabolism, Uracil containing | N-acetyl-beta-alanine | | 0.77 | 0.5149 | 1.38 | 0.4493 |
| Nucleotide | Pyrimidine Metabolism, Cytidine containing | cytidine 2',3'-cyclic monophosphate | HMDB11691 | 1.43 | 0.4104 | 1.93 | 0.1594 |
| Nucleotide | Pyrimidine Metabolism, Cytidine containing | cytidine | HMDB00089 | 0.85 | 0.6812 | 1.06 | 0.8888 |
| Nucleotide | Pyrimidine Metabolism, Cytidine containing | cytosine | HMDB00630 | 0.76 | 0.5880 | 1.38 | 0.5243 |
| Nucleotide | Pyrimidine Metabolism, Cytidine containing | 2'-deoxycytidine | HMDB00014 | 0.71 | 0.4040 | 0.86 | 0.7239 |
| Nucleotide | Pyrimidine Metabolism, Thymine containing | thymidine | HMDB00273 | 0.96 | 0.9088 | 1.13 | 0.7010 |
| Nucleotide | Pyrimidine Metabolism, Thymine containing | thymine | HMDB00262 | 0.9 | 0.6576 | 0.93 | 0.7613 |
| Nucleotide | Pyrimidine Metabolism, Thymine containing | 5,6-dihydrothymine | HMDB00079 | 0.99 | 0.9825 | 1.08 | 0.8696 |
| Nucleotide | Pyrimidine Metabolism, Thymine containing | 3-aminoisobutyrate | HMDB03911 | 1.23 | 0.6961 | 0.27 | 0.0175 |
| Nucleotide | Purine and Pyrimidine Metabolism | methylphosphate | HMDB61711 | 1.52 | 0.3528 | 1.41 | 0.4602 |
| Cofactors and Vitamins | Nicotinate and Nicotinamide Metabolism | quinolinate | HMDB00232 | 0.88 | 0.7272 | 0.44 | 0.0313 |

| | | | | | | | |
|---|---|---|---|---|---|---|---|
| Cofactors and Vitamins | Nicotinate and Nicotinamide Metabolism | nicotinate | HMDB01488 | 1.15 | 0.3958 | **1.6** | 0.0053 |
| Cofactors and Vitamins | Nicotinate and Nicotinamide Metabolism | nicotinate ribonucleoside | HMDB06809 | 1.22 | 0.5553 | 1.19 | 0.6008 |
| Cofactors and Vitamins | Nicotinate and Nicotinamide Metabolism | nicotinamide | HMDB01406 | 0.96 | 0.8468 | 0.88 | 0.5517 |
| Cofactors and Vitamins | Nicotinate and Nicotinamide Metabolism | nicotinamide riboside | HMDB00855 | 1.22 | 0.7072 | 0.59 | 0.3186 |
| Cofactors and Vitamins | Nicotinate and Nicotinamide Metabolism | 1-methylnicotinamide | HMDB00699 | 1 | 0.9885 | 1.09 | 0.8317 |
| Cofactors and Vitamins | Nicotinate and Nicotinamide Metabolism | 6-hydroxynicotinate | HMDB02658 | 0.81 | 0.6362 | 1.98 | 0.1440 |
| Cofactors and Vitamins | Nicotinate and Nicotinamide Metabolism | trigonelline (N'-methylnicotinate) | HMDB00875 | 0.86 | 0.6859 | 0.61 | 0.1842 |
| Cofactors and Vitamins | Nicotinate and Nicotinamide Metabolism | N1-Methyl-2-pyridone-5-carboxamide | HMDB04193 | 0.84 | 0.6932 | 0.8 | 0.6201 |
| Cofactors and Vitamins | Riboflavin Metabolism | riboflavin (Vitamin B2) | HMDB00244 | 1.12 | 0.6689 | 1.3 | 0.3478 |
| Cofactors and Vitamins | Pantothenate and CoA Metabolism | pantothenate | HMDB00210 | 1.02 | 0.9262 | 1.06 | 0.7748 |
| Cofactors and Vitamins | Ascorbate and Aldarate Metabolism | threonate | HMDB00943 | 0.58 | 0.1748 | 1.41 | 0.4148 |
| Cofactors and Vitamins | Ascorbate and Aldarate Metabolism | oxalate (ethanedioate) | HMDB02329 | 1.59 | 0.1511 | 1.2 | 0.5830 |
| Cofactors and Vitamins | Ascorbate and Aldarate Metabolism | gulonate* | HMDB03290 | 1.28 | 0.7074 | **0.3** | 0.0747 |
| Cofactors and Vitamins | Tocopherol Metabolism | alpha-tocopherol | HMDB01893 | 0.67 | 0.2817 | 1.25 | 0.5562 |
| Cofactors and Vitamins | Tocopherol Metabolism | alpha-tocopherol acetate | HMDB34227 | 1.1 | 0.7661 | 0.95 | 0.8804 |
| Cofactors and Vitamins | Tocopherol Metabolism | delta-tocopherol | HMDB02902 | 0.66 | 0.3030 | 1.74 | 0.1867 |
| Cofactors and Vitamins | Tocopherol Metabolism | alpha-tocotrienol | HMDB06327 | 0.69 | 0.3922 | **3.01** | 0.0168 |
| Cofactors and Vitamins | Tocopherol Metabolism | gamma-tocotrienol | HMDB12958 | 0.67 | 0.1909 | **2.53** | 0.0044 |
| Cofactors and Vitamins | Tocopherol Metabolism | gamma-CEHC | HMDB01931 | 0.84 | 0.5742 | 1.45 | 0.2479 |
| Cofactors and Vitamins | Tocopherol Metabolism | gamma-CEHC glucuronide* | | **0.41** | 0.0035 | 0.96 | 0.8862 |
| Cofactors and Vitamins | Tocopherol Metabolism | alpha-CEHC sulfate | | 0.85 | 0.7430 | 1.16 | 0.7833 |
| Cofactors and Vitamins | Tocopherol Metabolism | alpha-CEHC | HMDB01518 | 1.17 | 0.4215 | 0.95 | 0.7928 |
| Cofactors and Vitamins | Tocopherol Metabolism | gamma-tocopherol/beta-tocopherol | | 0.6 | 0.1370 | 1.58 | 0.1987 |

| | | | | | | | |
|---|---|---|---|---|---|---|---|
| Cofactors and Vitamins | Biotin Metabolism | biotin | HMDB00030 | 1.15 | 0.6334 | 1.17 | 0.6031 |
| Cofactors and Vitamins | Biotin Metabolism | biocytin | | 0.92 | 0.7880 | 1.12 | 0.7250 |
| Cofactors and Vitamins | Tetrahydrobiopterin Metabolism | biopterin | HMDB00468 | 0.98 | 0.9630 | 0.6 | 0.2419 |
| Cofactors and Vitamins | Pterin Metabolism | isoxanthopterin | HMDB00704 | 0.64 | 0.3937 | 0.95 | 0.9262 |
| Cofactors and Vitamins | Pterin Metabolism | pterin | HMDB00802 | 0.92 | 0.7667 | 0.82 | 0.5110 |
| Cofactors and Vitamins | Hemoglobin and Porphyrin Metabolism | protoporphyrin IX | HMDB00241 | 2.11 | 0.1471 | 1.2 | 0.7309 |
| Cofactors and Vitamins | Hemoglobin and Porphyrin Metabolism | bilirubin (Z,Z) | HMDB00054 | 1.19 | 0.7312 | 1.26 | 0.6548 |
| Cofactors and Vitamins | Hemoglobin and Porphyrin Metabolism | bilirubin (E,E)* | | 0.87 | 0.7219 | 1.36 | 0.4458 |
| Cofactors and Vitamins | Hemoglobin and Porphyrin Metabolism | bilirubin (E,Z or Z,E)* | HMDB00488 | 1.07 | 0.8111 | 1.63 | 0.1077 |
| Cofactors and Vitamins | Hemoglobin and Porphyrin Metabolism | biliverdin | HMDB01008 | 0.9 | 0.7848 | 0.9 | 0.7772 |
| Cofactors and Vitamins | Hemoglobin and Porphyrin Metabolism | I-urobilinogen | HMDB04157 | 0.96 | 0.9348 | 1.07 | 0.8931 |
| Cofactors and Vitamins | Hemoglobin and Porphyrin Metabolism | D-urobilin | HMDB04161 | 1.28 | 0.6174 | 1.02 | 0.9689 |
| Cofactors and Vitamins | Hemoglobin and Porphyrin Metabolism | L-urobilin | HMDB04159 | 0.86 | 0.7429 | 0.73 | 0.5097 |
| Cofactors and Vitamins | Hemoglobin and Porphyrin Metabolism | mesobilirubin | | 1.23 | 0.6486 | 1.58 | 0.3229 |
| Cofactors and Vitamins | Thiamine Metabolism | thiamin (Vitamin B1) | HMDB00235 | 1.17 | 0.7732 | 0.76 | 0.6255 |
| Cofactors and Vitamins | Thiamine Metabolism | 5-(2-Hydroxyethyl)-4-methylthiazole | | 1.14 | 0.8126 | 1.16 | 0.8009 |
| Cofactors and Vitamins | Thiamine Metabolism | hydroxymethylpyrimidine | | 0.83 | 0.6784 | 1.18 | 0.7248 |
| Cofactors and Vitamins | Vitamin A Metabolism | retinol (Vitamin A) | HMDB00305 | 0.86 | 0.4848 | 0.95 | 0.8181 |
| Cofactors and Vitamins | Vitamin A Metabolism | carotene diol (1) | | 0.67 | 0.3085 | 0.84 | 0.6577 |
| Cofactors and Vitamins | Vitamin A Metabolism | carotene diol (2) | | 0.79 | 0.4863 | 0.69 | 0.2912 |
| Cofactors and Vitamins | Vitamin A Metabolism | carotene diol (3) | | 1.02 | 0.9420 | 0.82 | 0.4935 |
| Cofactors and Vitamins | Vitamin A Metabolism | beta-cryptoxanthin | HMDB33844 | 0.99 | 0.9720 | 1.09 | 0.7715 |
| Cofactors and Vitamins | Vitamin A Metabolism | retinal | HMDB01358 | 0.93 | 0.5920 | 1 | 0.9823 |

| | | | | | | | |
|---|---|---|---|---|---|---|---|
| Cofactors and Vitamins | Vitamin B6 Metabolism | pyridoxine (Vitamin B6) | HMDB02075 | 1.58 | 0.3051 | 4.65 | 0.0011 |
| Cofactors and Vitamins | Vitamin B6 Metabolism | pyridoxamine | HMDB01431 | 1.06 | 0.8465 | 0.97 | 0.9033 |
| Cofactors and Vitamins | Vitamin B6 Metabolism | pyridoxal | HMDB01545 | 0.84 | 0.5038 | 0.92 | 0.7545 |
| Cofactors and Vitamins | Vitamin B6 Metabolism | pyridoxate | HMDB00017 | 0.86 | 0.5441 | 2.34 | 0.0014 |
| Xenobiotics | Benzoate Metabolism | hippurate | HMDB00714 | 0.4 | 0.1330 | 1.23 | 0.7491 |
| Xenobiotics | Benzoate Metabolism | 2-hydroxyhippurate (salicylurate) | HMDB00840 | 0.57 | 0.1420 | 1.23 | 0.6105 |
| Xenobiotics | Benzoate Metabolism | 3-hydroxyhippurate | HMDB06116 | 0.48 | 0.0940 | 1.25 | 0.6159 |
| Xenobiotics | Benzoate Metabolism | 4-hydroxyhippurate | HMDB13678 | 0.35 | 0.0570 | 1.36 | 0.5851 |
| Xenobiotics | Benzoate Metabolism | 4-hydroxymandelate | HMDB00822 | 0.61 | 0.2176 | 1.31 | 0.5274 |
| Xenobiotics | Benzoate Metabolism | benzoate | HMDB01870 | 0.89 | 0.6220 | 0.96 | 0.8821 |
| Xenobiotics | Benzoate Metabolism | 4-hydroxybenzoate | HMDB00500 | 0.88 | 0.6260 | 1.8 | 0.0272 |
| Xenobiotics | Benzoate Metabolism | 3-hydroxybenzoate | HMDB02466 | 1.24 | 0.5385 | 1.08 | 0.8367 |
| Xenobiotics | Benzoate Metabolism | 2,4,6-trihydroxybenzoate | HMDB29649 | 0.92 | 0.8619 | 1.43 | 0.4928 |
| Xenobiotics | Benzoate Metabolism | 3,4-dihydroxybenzoate | HMDB01856 | 0.83 | 0.6844 | 1.96 | 0.1685 |
| Xenobiotics | Benzoate Metabolism | catechol sulfate | HMDB59724 | 1.2 | 0.7046 | 1.64 | 0.3322 |
| Xenobiotics | Benzoate Metabolism | 4-methylcatechol sulfate | | 1.13 | 0.7115 | 1.3 | 0.4690 |
| Xenobiotics | Benzoate Metabolism | p-hydroxybenzaldehyde | HMDB11718 | 0.79 | 0.6014 | 1.95 | 0.1528 |
| Xenobiotics | Benzoate Metabolism | methyl-4-hydroxybenzoate | HMDB32572 | 0.57 | 0.0299 | 0.9 | 0.7061 |
| Xenobiotics | Benzoate Metabolism | 4-ethylphenylsulfate | | 1 | 1.0000 | 1.19 | 0.1087 |
| Xenobiotics | Benzoate Metabolism | 4-vinylphenol sulfate | HMDB04072 | 0.97 | 0.8993 | 1.11 | 0.6532 |
| Xenobiotics | Benzoate Metabolism | 3-methoxycatechol sulfate (1) | | 0.98 | 0.9277 | 1.34 | 0.2239 |
| Xenobiotics | Benzoate Metabolism | methyl-4-hydroxybenzoate sulfate | | 0.92 | 0.7914 | 1.43 | 0.2899 |
| Xenobiotics | Benzoate Metabolism | propyl 4-hydroxybenzoate | HMDB32574 | 0.95 | 0.1981 | 0.96 | 0.3164 |
| Xenobiotics | Benzoate Metabolism | propyl 4-hydroxybenzoate sulfate | | 1.23 | 0.2524 | 1 | 0.9802 |
| Xenobiotics | Benzoate Metabolism | p-cresol | HMDB01858 | 0.96 | 0.9482 | 1.36 | 0.6217 |
| Xenobiotics | Benzoate Metabolism | p-cresol sulfate | HMDB11635 | 0.73 | 0.6344 | 0.67 | 0.5550 |
| Xenobiotics | Benzoate Metabolism | o-cresol sulfate | | 1 | 0.9679 | 1.05 | 0.5902 |
| Xenobiotics | Benzoate Metabolism | phenylpropionylglycine | HMDB00860 | 1.6 | 0.2274 | 1.09 | 0.8282 |
| Xenobiotics | Benzoate Metabolism | 2-methylbutyrylphenylalanine | | 0.72 | 0.4425 | 1.52 | 0.3361 |
| Xenobiotics | Benzoate Metabolism | 3-(3-hydroxyphenyl)propionate sulfate | | 0.9 | 0.8253 | 1.55 | 0.3685 |
| Xenobiotics | Benzoate Metabolism | 3-(3-hydroxyphenyl)propionate | HMDB00375 | 1.03 | 0.9652 | 1.28 | 0.7287 |
| Xenobiotics | Benzoate Metabolism | 3-(4-hydroxyphenyl)propionate | HMDB02199 | 1.08 | 0.8898 | 6.94 | 0.0007 |

| | | | | | | | |
|---|---|---|---|---|---|---|---|
| Xenobiotics | Benzoate Metabolism | 3-phenylpropionate (hydrocinnamate) | HMDB00764 | 2.01 | 0.2463 | 0.91 | 0.8777 |
| Xenobiotics | Xanthine Metabolism | paraxanthine | HMDB01860 | 1.27 | 0.4562 | 1.17 | 0.6330 |
| Xenobiotics | Xanthine Metabolism | theobromine | HMDB02825 | 1 | 0.9980 | 0.68 | 0.3008 |
| Xenobiotics | Xanthine Metabolism | theophylline | HMDB01889 | 0.86 | 0.6266 | **0.44** | 0.0091 |
| Xenobiotics | Xanthine Metabolism | 1-methylurate | HMDB03099 | 1.06 | 0.8976 | 1.05 | 0.9228 |
| Xenobiotics | Xanthine Metabolism | 7-methylurate | HMDB11107 | 0.62 | 0.2774 | 0.92 | 0.8609 |
| Xenobiotics | Xanthine Metabolism | 1,3-dimethylurate | HMDB01857 | 0.74 | 0.4749 | 1.11 | 0.8098 |
| Xenobiotics | Xanthine Metabolism | 1,7-dimethylurate | HMDB11103 | 0.74 | 0.5596 | 1.47 | 0.4542 |
| Xenobiotics | Xanthine Metabolism | 3,7-dimethylurate | HMDB01982 | 1.04 | 0.9384 | 1.08 | 0.8762 |
| Xenobiotics | Xanthine Metabolism | 1,3,7-trimethylurate | HMDB02123 | 0.79 | 0.5763 | 1.06 | 0.8940 |
| Xenobiotics | Xanthine Metabolism | 1-methylxanthine | HMDB10738 | 0.98 | 0.9426 | 0.86 | 0.6562 |
| Xenobiotics | Xanthine Metabolism | 3-methylxanthine | HMDB01886 | 0.93 | 0.8408 | **0.55** | 0.0916 |
| Xenobiotics | Xanthine Metabolism | 5-acetylamino-6-amino-3-methyluracil | HMDB04400 | 0.79 | 0.6607 | 1.01 | 0.9838 |
| Xenobiotics | Xanthine Metabolism | 5-acetylamino-6-formylamino-3-methyluracil | HMDB11105 | 1.05 | 0.9049 | 1.04 | 0.9215 |
| Xenobiotics | Xanthine Metabolism | caffeic acid sulfate | HMDB41708 | 0.91 | 0.8037 | 1.12 | 0.7600 |
| Xenobiotics | Food Component/Plant | maltol | HMDB30776 | 0.9 | 0.5747 | 0.95 | 0.8153 |
| Xenobiotics | Food Component/Plant | piperidine | HMDB34301 | 1.41 | 0.3320 | 0.86 | 0.6818 |
| Xenobiotics | Food Component/Plant | 2-piperidinone | HMDB11749 | 0.94 | 0.8768 | 0.96 | 0.9299 |
| Xenobiotics | Food Component/Plant | kaempferol | HMDB05801 | 1.27 | 0.6236 | 1.52 | 0.4146 |
| Xenobiotics | Food Component/Plant | kaempferol-3-rhamnoside | | 1 | | 1 | |
| Xenobiotics | Food Component/Plant | dihydrokaempferol | HMDB30847 | 0.92 | 0.8667 | 1.09 | 0.8751 |
| Xenobiotics | Food Component/Plant | quercitrin | HMDB33751 | 1.09 | 0.8248 | 1.17 | 0.6879 |
| Xenobiotics | Food Component/Plant | indoleacetylaspartate | HMDB38666 | 1.1 | 0.7543 | **2.14** | 0.0134 |
| Xenobiotics | Food Component/Plant | erythrose | HMDB02649 | 0.55 | 0.2569 | 0.98 | 0.9725 |
| Xenobiotics | Food Component/Plant | sucralose | HMDB31554 | 2.51 | 0.1693 | 0.79 | 0.7311 |
| Xenobiotics | Food Component/Plant | genistein | HMDB03217 | **0.34** | 0.0769 | 0.81 | 0.7389 |
| Xenobiotics | Food Component/Plant | 3-dehydroshikimate | | 0.82 | 0.6965 | 0.89 | 0.8272 |
| Xenobiotics | Food Component/Plant | catechin | HMDB02780 | 0.89 | 0.7170 | 0.96 | 0.9077 |
| Xenobiotics | Food Component/Plant | epicatechin | HMDB01871 | 1.12 | 0.6533 | 0.89 | 0.6721 |
| Xenobiotics | Food Component/Plant | 1-methyl-beta-carboline-3-carboxylic acid | | 0.85 | 0.6275 | 1.06 | 0.8564 |
| Xenobiotics | Food Component/Plant | diaminopimelate | HMDB01370 | 1.28 | 0.5148 | 1.24 | 0.5802 |
| Xenobiotics | Food Component/Plant | 1-kestose | HMDB11729 | 0.57 | 0.2077 | 1.43 | 0.4366 |
| Xenobiotics | Food Component/Plant | apigenin | HMDB02124 | 0.7 | 0.4986 | 1.16 | 0.7830 |
| Xenobiotics | Food Component/Plant | luteolin | HMDB05800 | 0.55 | 0.2694 | 1.07 | 0.9083 |

| | | | | | | | |
|---|---|---|---|---|---|---|---|
| Xenobiotics | Food Component/Plant | levulinate (4-oxovalerate) | HMDB00720 | 1.14 | 0.6335 | 0.71 | 0.2171 |
| Xenobiotics | Food Component/Plant | vanillate | HMDB00484 | 0.82 | 0.5775 | **2.27** | 0.0246 |
| Xenobiotics | Food Component/Plant | 2,3-dihydroxyisovalerate | HMDB12141 | 0.65 | 0.2899 | 0.7 | 0.3880 |
| Xenobiotics | Food Component/Plant | 2,8-quinolinediol | | 0.94 | 0.8968 | 1.17 | 0.7464 |
| Xenobiotics | Food Component/Plant | 2,8-quinolinediol sulfate | | 1.44 | 0.3805 | **2.14** | 0.0846 |
| Xenobiotics | Food Component/Plant | 20-hydroxyecdysone | HMDB30180 | 0.96 | 0.8670 | 0.86 | 0.5683 |
| Xenobiotics | Food Component/Plant | 2-isopropylmalate | HMDB00402 | 1.2 | 0.7211 | 0.88 | 0.8076 |
| Xenobiotics | Food Component/Plant | 2-oxindole-3-acetate | HMDB35514 | 1.25 | 0.6189 | 1.83 | 0.1879 |
| Xenobiotics | Food Component/Plant | 3,5-dihydroxybenzoic acid | HMDB13677 | **1.5** | 0.0546 | 1.06 | 0.7754 |
| Xenobiotics | Food Component/Plant | 4-hydroxybenzyl alcohol | HMDB11724 | 1.06 | 0.8650 | 1.68 | 0.1742 |
| Xenobiotics | Food Component/Plant | gluconate | HMDB00625 | 1.16 | 0.8017 | 2.44 | 0.1445 |
| Xenobiotics | Food Component/Plant | 2-acetolactate | HMDB06833 | 1.01 | 0.9810 | 1.09 | 0.7308 |
| Xenobiotics | Food Component/Plant | apiin | | **0.8** | 0.0667 | 0.88 | 0.3489 |
| Xenobiotics | Food Component/Plant | beta-guanidinopropanoate | HMDB13222 | 0.73 | 0.3985 | 1.15 | 0.7159 |
| Xenobiotics | Food Component/Plant | caffeate | HMDB01964 | 0.46 | 0.2835 | 2.64 | 0.1980 |
| Xenobiotics | Food Component/Plant | chrysoeriol | HMDB30667 | 0.69 | 0.4838 | 1.54 | 0.4264 |
| Xenobiotics | Food Component/Plant | coumaroylquinate (1) | | 0.89 | 0.7935 | 1.54 | 0.3380 |
| Xenobiotics | Food Component/Plant | coumaroylquinate (2) | | 1.02 | 0.9191 | 0.97 | 0.8920 |
| Xenobiotics | Food Component/Plant | coumaroylquinate (3) | | 1 | 0.9930 | 1.28 | 0.2698 |
| Xenobiotics | Food Component/Plant | coumaroylquinate (4) | | 0.92 | 0.5933 | 1.01 | 0.9683 |
| Xenobiotics | Food Component/Plant | coumaroylquinate (5) | | 1.06 | 0.7593 | 1 | 0.9976 |
| Xenobiotics | Food Component/Plant | cryptochlorogenic acid | | 0.69 | 0.2018 | 0.72 | 0.2820 |
| Xenobiotics | Food Component/Plant | daidzein | HMDB03312 | 0.41 | 0.1369 | 0.53 | 0.3065 |
| Xenobiotics | Food Component/Plant | deoxymugineic acid | | 0.6 | 0.2733 | **4.5** | 0.0021 |
| Xenobiotics | Food Component/Plant | digalacturonic acid | HMDB39721 | 1.34 | 0.5550 | 0.62 | 0.3537 |
| Xenobiotics | Food Component/Plant | dihydroferulic acid | | 0.63 | 0.4522 | **6.99** | 0.0032 |
| Xenobiotics | Food Component/Plant | dihydroquercetin | | 0.95 | 0.8679 | 0.85 | 0.6083 |
| Xenobiotics | Food Component/Plant | enterolactone | | 1.91 | 0.1988 | 1.14 | 0.7973 |
| Xenobiotics | Food Component/Plant | equol sulfate | | 0.82 | 0.3446 | 1.04 | 0.8524 |
| Xenobiotics | Food Component/Plant | eriodictyol | HMDB05810 | 0.62 | 0.4002 | 1.31 | 0.6467 |
| Xenobiotics | Food Component/Plant | erythritol | HMDB02994 | 0.49 | 0.1772 | 0.95 | 0.9294 |
| Xenobiotics | Food Component/Plant | ferulate | HMDB00954 | 1.24 | 0.6403 | **3.5** | 0.0100 |
| Xenobiotics | Food Component/Plant | ferulic acid 4-sulfate | HMDB29200 | 1.14 | 0.8494 | **4.88** | 0.0274 |
| Xenobiotics | Food Component/Plant | ferulylglycine (1) | | **0.4** | 0.0256 | **2.16** | 0.0665 |

| | | | | | | | |
|---|---|---|---|---|---|---|---|
| Xenobiotics | Food Component/Plant | fucitol | | 0.57 | 0.1015 | 1.32 | 0.4324 |
| Xenobiotics | Food Component/Plant | glycitein | HMDB05781 | 0.54 | 0.1195 | 0.7 | 0.3797 |
| Xenobiotics | Food Component/Plant | glycyrrhetinate | HMDB11628 | 1 | | 1 | |
| Xenobiotics | Food Component/Plant | hesperetin | HMDB05782 | **0.28** | 0.0647 | 0.65 | 0.5465 |
| Xenobiotics | Food Component/Plant | indoleacrylate | HMDB00734 | 1.85 | 0.1317 | 0.78 | 0.5663 |
| Xenobiotics | Food Component/Plant | indolin-2-one | | 1.17 | 0.7388 | 0.62 | 0.3251 |
| Xenobiotics | Food Component/Plant | isovitexin | | 1.05 | 0.9119 | 1.27 | 0.5695 |
| Xenobiotics | Food Component/Plant | methyl indole-3-acetate | HMDB29738 | 0.69 | 0.1778 | 1.26 | 0.4184 |
| Xenobiotics | Food Component/Plant | N-(2-furoyl)glycine | HMDB00439 | **0.62** | 0.0932 | 1.01 | 0.9776 |
| Xenobiotics | Food Component/Plant | naringenin | HMDB02670 | 0.64 | 0.4783 | 0.84 | 0.7920 |
| Xenobiotics | Food Component/Plant | N-glycolylneuraminate | HMDB00833 | 0.77 | 0.2565 | 1.02 | 0.9366 |
| Xenobiotics | Food Component/Plant | nicotianamine | | 0.37 | 0.1351 | 2.99 | 0.1092 |
| Xenobiotics | Food Component/Plant | pheophorbide A | | 1.05 | 0.9323 | 0.46 | 0.2056 |
| Xenobiotics | Food Component/Plant | phytanate | HMDB00801 | 1.01 | 0.9768 | 1.12 | 0.8120 |
| Xenobiotics | Food Component/Plant | piperine | HMDB29377 | 1.11 | 0.7986 | 0.94 | 0.8930 |
| Xenobiotics | Food Component/Plant | ponciretin | | 0.61 | 0.3313 | 1.88 | 0.2317 |
| Xenobiotics | Food Component/Plant | quinate | HMDB03072 | 1.19 | 0.7376 | 1.12 | 0.8341 |
| Xenobiotics | Food Component/Plant | rosmarinate | HMDB03572 | **0.57** | 0.0206 | 1.28 | 0.3008 |
| Xenobiotics | Food Component/Plant | saccharin | HMDB29723 | 1.78 | 0.4042 | 1.16 | 0.8351 |
| Xenobiotics | Food Component/Plant | acesulfame | HMDB33585 | 0.49 | 0.3540 | 1.97 | 0.3945 |
| Xenobiotics | Food Component/Plant | secoisolariciresinol | HMDB13692 | 0.76 | 0.4516 | 1.51 | 0.2763 |
| Xenobiotics | Food Component/Plant | sinapate | HMDB32616 | 0.91 | 0.8010 | **1.9** | 0.0882 |
| Xenobiotics | Food Component/Plant | solanidine | HMDB03236 | 0.79 | 0.7392 | 0.89 | 0.8705 |
| Xenobiotics | Food Component/Plant | soyasaponin I | HMDB34649 | 0.79 | 0.6918 | 2.18 | 0.2059 |
| Xenobiotics | Food Component/Plant | soyasaponin II | HMDB34650 | 0.52 | 0.1238 | 0.65 | 0.3384 |
| Xenobiotics | Food Component/Plant | soyasaponin III | | 0.87 | 0.7942 | 0.8 | 0.6928 |
| Xenobiotics | Food Component/Plant | stachydrine | HMDB04827 | 0.65 | 0.3577 | 0.75 | 0.5442 |
| Xenobiotics | Food Component/Plant | syringic acid | HMDB02085 | 0.64 | 0.3822 | 2.08 | 0.1708 |
| Xenobiotics | Food Component/Plant | tartarate | HMDB00956 | 1.47 | 0.3922 | 0.95 | 0.9210 |
| Xenobiotics | Food Component/Plant | tribuloside | | 1 | 1.0000 | **1.13** | 0.0616 |
| Xenobiotics | Food Component/Plant | tyrosol | HMDB04284 | 1.01 | 0.9832 | **1.98** | 0.0412 |
| Xenobiotics | Food Component/Plant | vitexin | | 0.48 | 0.1505 | 2.21 | 0.1271 |
| Xenobiotics | Food Component/Plant | methyl glucopyranoside (alpha + beta) | | 0.73 | 0.3141 | 0.9 | 0.7565 |
| Xenobiotics | Food Component/Plant | capsaicin | HMDB02227 | 0.98 | 0.9412 | 0.86 | 0.5188 |

| | | | | | | | |
|---|---|---|---|---|---|---|---|
| Xenobiotics | Food Component/Plant | hesperidin | HMDB03265 | 0.65 | 0.4507 | 0.6 | 0.4026 |
| Xenobiotics | Food Component/Plant | sinigrin | HMDB34070 | 1 | 0.9796 | 1 | 0.9744 |
| Xenobiotics | Food Component/Plant | enterodiol | | 1.53 | 0.3754 | 1.1 | 0.8449 |
| Xenobiotics | Food Component/Plant | limonin | | 0.69 | 0.1313 | 1.01 | 0.9653 |
| Xenobiotics | Food Component/Plant | narirutin | | 0.98 | 0.9781 | 0.62 | 0.2312 |
| Xenobiotics | Food Component/Plant | neoponcirin | | 0.99 | 0.9754 | 0.91 | 0.7390 |
| Xenobiotics | Food Component/Plant | harmane | HMDB35196 | 1 | 1.0000 | 1.04 | 0.9076 |
| Xenobiotics | Food Component/Plant | diosmetin | HMDB29676 | **0.29** | 0.0192 | 1.25 | 0.6872 |
| Xenobiotics | Food Component/Plant | synephrine | HMDB04826 | 0.75 | 0.1673 | 1.01 | 0.9795 |
| Xenobiotics | Food Component/Plant | nomilin | | 0.83 | 0.5771 | 0.9 | 0.7641 |
| Xenobiotics | Food Component/Plant | pyrraline | HMDB33143 | 0.56 | 0.1115 | 1.56 | 0.2327 |
| Xenobiotics | Food Component/Plant | umbelliferone sulfate | | 1.06 | 0.8178 | 1.27 | 0.3639 |
| Xenobiotics | Food Component/Plant | daidzein sulfate (2) | | **0.39** | 0.0123 | 1.38 | 0.4129 |
| Xenobiotics | Food Component/Plant | N-acetylpyrraline | | 0.93 | 0.7928 | 1.05 | 0.8678 |
| Xenobiotics | Food Component/Plant | daidzein sulfate (1) | | **0.35** | 0.0023 | 1.04 | 0.9083 |
| Xenobiotics | Food Component/Plant | 2-keto-3-deoxy-gluconate | HMDB01353 | 0.76 | 0.5048 | 1 | 0.9966 |
| Xenobiotics | Food Component/Plant | 3-methylsulfinylpropyl-glucosinolate | HMDB38406 | 1 | | 1 | |
| Xenobiotics | Food Component/Plant | kaempferol 3-O-glucoside/galactoside | | 1 | | 1 | |
| Xenobiotics | Food Component/Plant | 1,2-dilinolenoyl-digalactosylglycerol (18:3/18:3) | | 0.97 | 0.8932 | 0.64 | 0.1021 |
| Xenobiotics | Food Component/Plant | 1-palmitoyl-2-linolenoyl-digalactosylglycerol (16:0/18:3) | | 0.9 | 0.7860 | 0.67 | 0.3250 |
| Xenobiotics | Food Component/Plant | dihydrocaffeate | HMDB00423 | 2.17 | 0.2086 | 0.97 | 0.9646 |
| Xenobiotics | Food Component/Plant | 4-hydroxycinnamate | HMDB02035 | 1.67 | 0.2786 | 2.01 | 0.1566 |
| Xenobiotics | Food Component/Plant | DIMBOA | | 1.01 | 0.9574 | 1.13 | 0.5366 |
| Xenobiotics | Food Component/Plant | Urolithin A | HMDB13695 | 1.33 | 0.2855 | 1.18 | 0.5365 |
| Xenobiotics | Food Component/Plant | luteolin-8-C-glucoside | | 0.87 | 0.7626 | 2.04 | 0.1241 |
| Xenobiotics | Food Component/Plant | malonylgenistin | | **0.51** | 0.0023 | 0.99 | 0.9670 |
| Xenobiotics | Bacterial/Fungal | 2-acetamidobutanoate | | 0.95 | 0.9003 | 0.92 | 0.8331 |
| Xenobiotics | Bacterial/Fungal | glutamyl-meso-diaminopimelate | | 1.31 | 0.5884 | 0.45 | 0.1286 |
| Xenobiotics | Bacterial/Fungal | N-acetylmuramyl-alanyl-isoglutamine | | 0.64 | 0.2656 | 1.77 | 0.1734 |
| Xenobiotics | Bacterial/Fungal | N-methylpipecolate | | 0.66 | 0.1234 | 0.96 | 0.8696 |
| Xenobiotics | Bacterial/Fungal | Urolithin B | HMDB13696 | 0.96 | 0.8379 | 0.97 | 0.9008 |
| Xenobiotics | Drug - Analgesics, Anesthetics | 4-acetamidophenol | HMDB01859 | 0.61 | 0.4231 | 0.6 | 0.4287 |
| Xenobiotics | Drug - Analgesics, Anesthetics | 3-(N-acetyl-L-cystein-S-yl) acetaminophen | | 0.68 | 0.5408 | 0.83 | 0.7780 |

| | | | | | | | |
|---|---|---|---|---|---|---|---|
| Xenobiotics | Drug - Analgesics, Anesthetics | 4-acetaminophen sulfate | HMDB59911 | 0.87 | 0.8338 | 0.57 | 0.4359 |
| Xenobiotics | Drug - Analgesics, Anesthetics | 4-acetamidophenylglucuronide | HMDB10316 | 0.99 | 0.0484 | 1 | 1.0000 |
| Xenobiotics | Drug - Analgesics, Anesthetics | 2-hydroxyacetaminophen sulfate* | | 0.84 | 0.6788 | 1.01 | 0.9829 |
| Xenobiotics | Drug - Analgesics, Anesthetics | 2-methoxyacetaminophen sulfate* | | 0.75 | 0.5363 | 0.85 | 0.7320 |
| Xenobiotics | Drug - Analgesics, Anesthetics | 2-methoxyacetaminophen glucuronide* | | 0.77 | 0.0049 | 1 | 1.0000 |
| Xenobiotics | Drug - Analgesics, Anesthetics | 3-(cystein-S-yl)acetaminophen* | | 0.91 | 0.7472 | 0.96 | 0.8792 |
| Xenobiotics | Drug - Analgesics, Anesthetics | 2-acetamidophenol sulfate | | 0.93 | 0.7815 | 1.25 | 0.4052 |
| Xenobiotics | Drug - Analgesics, Anesthetics | 4-aminophenol sulfate (2) | | 1.14 | 0.3558 | 0.95 | 0.7482 |
| Xenobiotics | Drug - Analgesics, Anesthetics | carboxyibuprofen | HMDB60564 | 1 | 1.0000 | 0.97 | 0.1620 |
| Xenobiotics | Drug - Analgesics, Anesthetics | diclofenac | HMDB14724 | 1 | | 1 | |
| Xenobiotics | Drug - Antibiotic | azithromycin | HMDB14352 | 0.55 | 0.1555 | 1 | 1.0000 |
| Xenobiotics | Drug - Antibiotic | amoxicillin | HMDB15193 | 0.8 | 0.2421 | 0.96 | 0.8232 |
| Xenobiotics | Drug - Antibiotic | sulfamethoxazole | HMDB15150 | 0.99 | 0.0803 | 0.99 | 0.3564 |
| Xenobiotics | Drug - Antibiotic | metronidazole | HMDB15052 | 1 | 0.9518 | 1 | 0.8696 |
| Xenobiotics | Drug - Antibiotic | clotrimazole | HMDB01922 | 0.88 | 0.6602 | 0.94 | 0.8433 |
| Xenobiotics | Drug - Antibiotic | fluconazole | HMDB14342 | 1 | | 1 | |
| Xenobiotics | Drug - Antibiotic | 3-hydroxyquinine | HMDB01091 | 1 | 0.9788 | 1 | 0.9186 |
| Xenobiotics | Drug - Antibiotic | quinine | | 1 | 1.0000 | 0.99 | 0.8757 |
| Xenobiotics | Drug - Gastrointestinal | promethazine | HMDB15202 | 1 | 1.0000 | 0.84 | 0.2494 |
| Xenobiotics | Drug - Gastrointestinal | ranitidine | HMDB01930 | 1 | | 1 | |
| Xenobiotics | Drug - Respiratory | diphenhydramine | HMDB01927 | 1 | 0.9313 | 1 | 1.0000 |
| Xenobiotics | Drug - Respiratory | loratadine | HMDB05000 | 1.05 | 0.5450 | 1 | 1.0000 |
| Xenobiotics | Drug - Topical Agents | salicylate | HMDB01895 | 1.78 | 0.0970 | 4.67 | 0.0000 |
| Xenobiotics | Drug - Topical Agents | hydroquinone sulfate | HMDB02434 | 1.03 | 0.9321 | 1.94 | 0.0898 |
| Xenobiotics | Drug - Other | S-carboxymethyl-L-cysteine | HMDB29415 | 0.46 | 0.0857 | 0.98 | 0.9662 |
| Xenobiotics | Chemical | 1,3-propanediol | | 1.09 | 0.8425 | 1.77 | 0.2279 |
| Xenobiotics | Chemical | 3-hydroxypyridine | | 1.54 | 0.1970 | 1.05 | 0.8788 |
| Xenobiotics | Chemical | sulfate* | HMDB01448 | 0.89 | 0.6110 | 0.89 | 0.6485 |
| Xenobiotics | Chemical | O-sulfo-L-tyrosine | | 0.43 | 0.1445 | 1.15 | 0.8168 |
| Xenobiotics | Chemical | 2-aminophenol sulfate | HMDB61116 | 0.87 | 0.6264 | 1 | 1.0000 |
| Xenobiotics | Chemical | 1,4-butanediol | | 1 | 1.0000 | 1.08 | 0.7333 |
| Xenobiotics | Chemical | S-(3-hydroxypropyl)mercapturic acid (HPMA) | | 0.87 | 0.7359 | 1.28 | 0.5669 |
| Xenobiotics | Chemical | 5-aminoimidazole-4-carboxamide | HMDB03192 | 0.52 | 0.1253 | 0.91 | 0.8232 |
| Xenobiotic | Chemical | dexpanthenol | HMDB042 | 0.9 | 0.503 | 1 | 1.000 |

| | | | | | | | |
|---|---|---|---|---|---|---|---|
| s | | | 31 | | 1 | | 0 |
| Xenobiotics | Chemical | ectoine | | 0.9 | 0.7470 | 0.85 | 0.6109 |
| Xenobiotics | Chemical | HEPES | | 1 | 0.9419 | 1 | 0.9732 |
| Xenobiotics | Chemical | N-propionylmethionine | | 1.09 | 0.8631 | **3.6** | 0.0111 |
| Xenobiotics | Chemical | azeloylcarnitine (C9-DC) | | 0.84 | 0.4583 | 0.97 | 0.8935 |
| Xenobiotics | Chemical | succinimide | | 0.84 | 0.3722 | 0.84 | 0.3862 |
| Xenobiotics | Chemical | triethanolamine | HMDB32538 | 0.7 | 0.2556 | 0.95 | 0.8765 |
| Xenobiotics | Chemical | trizma acetate | | 1 | | 1 | |
| Xenobiotics | Chemical | 2,4-dihydroxyhydrocinnamate | | 0.96 | 0.8770 | 1 | 0.9923 |
| Xenobiotics | Chemical | 4-hydroxychlorothalonil | | 1.35 | 0.2771 | 1 | 0.9876 |
| Xenobiotics | Chemical | 4-thiouracil | | 0.92 | 0.7698 | 1.21 | 0.4962 |
| Xenobiotics | Chemical | 1,2,3-benzenetriol sulfate (2) | | 0.73 | 0.4073 | **2.01** | 0.0820 |
| Xenobiotics | Chemical | 1,2,3-benzenetriol sulfate (1) | | 0.91 | 0.6512 | 1.21 | 0.3785 |
| Xenobiotics | Chemical | 4-acetamidobenzoate | | 0.97 | 0.9339 | 1.21 | 0.5847 |
| Xenobiotics | Chemical | thioproline | | 0.59 | 0.2682 | 1.53 | 0.3952 |
| N/A | N/A | amodiaquine | HMDB14751 | 1 | 0.9262 | 1 | 0.9533 |
| N/A | N/A | X - 02249 - retired for 3-Cmpfp** | | 0.68 | 0.2019 | 1.63 | 0.1223 |
| N/A | N/A | X - 09789 | | 0.85 | 0.6316 | **2.63** | 0.0056 |
| N/A | N/A | X - 10457 | | 0.83 | 0.4117 | 1.17 | 0.4845 |
| N/A | N/A | X - 10458 | | 1 | 0.9591 | 0.97 | 0.5751 |
| N/A | N/A | X - 11261 | | 0.88 | 0.6734 | 1.15 | 0.6702 |
| N/A | N/A | X - 11308 | | 0.66 | 0.2841 | 0.95 | 0.8969 |
| N/A | N/A | X - 11357 | | 0.75 | 0.5199 | 1.04 | 0.9281 |
| N/A | N/A | X - 11407 | | 0.8 | 0.2742 | **2.03** | 0.0016 |
| N/A | N/A | X - 11429 - retired for 5,6-dihydrouridine | | 0.59 | 0.2394 | 1.81 | 0.2054 |
| N/A | N/A | X - 11440 | | 0.91 | 0.8549 | 0.93 | 0.8832 |
| N/A | N/A | X - 11441 | | 0.89 | 0.7432 | 0.75 | 0.4288 |
| N/A | N/A | X - 11442 | | 1 | 0.9931 | 0.74 | 0.4539 |
| N/A | N/A | X - 11444 | | **0.61** | 0.0260 | 1.16 | 0.5270 |
| N/A | N/A | X - 11452 - retired for sulfate of piperine metabolite C16H19NO3 (2)* | | 0.54 | 0.1618 | 1.25 | 0.6269 |
| N/A | N/A | X - 11470 | | **0.65** | 0.0426 | 1.11 | 0.6496 |
| N/A | N/A | X - 11491 | | **0.39** | 0.0859 | 1.02 | 0.9670 |
| N/A | N/A | X - 11522 | | 0.82 | 0.5580 | 0.83 | 0.6015 |
| N/A | N/A | X - 11530 | | 1.04 | 0.9272 | 0.78 | 0.6179 |

| | | | | | | | |
|---|---|---|---|---|---|---|---|
| N/A | N/A | X - 11538 - retired for octadecenedioate (C18:1-DC) | | 0.99 | 0.965 9 | 1.66 | 0.120 9 |
| N/A | N/A | X - 11540 - retired for 5-dodecenoylcarnitine | | 0.89 | 0.830 5 | 2.16 | 0.190 6 |
| N/A | N/A | X - 11564 | | 0.72 | 0.336 8 | 0.84 | 0.617 6 |
| N/A | N/A | X - 11612 | | 0.77 | 0.586 0 | 1 | 1.000 0 |
| N/A | N/A | X - 11632 | | 0.95 | 0.859 0 | 1.11 | 0.695 6 |
| N/A | N/A | X - 11640 | | 1.65 | 0.185 4 | 1.57 | 0.247 6 |
| N/A | N/A | X - 11648 | | 0.94 | 0.902 2 | 0.75 | 0.585 4 |
| N/A | N/A | X - 11787 | | 0.86 | 0.576 5 | 1.3 | 0.334 8 |
| N/A | N/A | X - 11795 | | 0.95 | 0.911 7 | 0.9 | 0.836 0 |
| N/A | N/A | X - 11838 | | 1.94 | 0.107 7 | 0.78 | 0.569 2 |
| N/A | N/A | X - 11905 - retired for hexadecenedioate (C16:1-DC)* | | 1.31 | 0.193 6 | 1.19 | 0.410 1 |
| N/A | N/A | X - 11979 | | 0.78 | 0.185 6 | 1 | 1.000 0 |
| N/A | N/A | X - 12007 | | 0.76 | 0.509 4 | 1.22 | 0.653 6 |
| N/A | N/A | X - 12013 | | 1.01 | 0.983 4 | 1.49 | 0.406 3 |
| N/A | N/A | X - 12015 | | 1.42 | 0.481 5 | 1.3 | 0.608 9 |
| N/A | N/A | X - 12026 | | 0.8 | 0.498 2 | 1.4 | 0.333 5 |
| N/A | N/A | X - 12027 | | 1.09 | 0.763 8 | **1.93** | 0.034 1 |
| N/A | N/A | X - 12093 | | 0.85 | 0.689 2 | 0.93 | 0.852 5 |
| N/A | N/A | X - 12096 | | 0.77 | 0.492 4 | 1.28 | 0.519 0 |
| N/A | N/A | X - 12100 | | **0.66** | 0.072 9 | 1.07 | 0.772 2 |
| N/A | N/A | X - 12101 | | 0.73 | 0.463 8 | 0.76 | 0.531 9 |
| N/A | N/A | X - 12117 | | **0.57** | 0.054 2 | 0.91 | 0.751 1 |
| N/A | N/A | X - 12125 | | 1.34 | 0.388 2 | 0.77 | 0.471 6 |
| N/A | N/A | X - 12127 | | 0.54 | 0.236 7 | 1.21 | 0.722 6 |
| N/A | N/A | X - 12199 | | **1.66** | 0.037 3 | 0.69 | 0.153 8 |
| N/A | N/A | X - 12206 | | 0.51 | 0.165 8 | 1.14 | 0.791 0 |
| N/A | N/A | X - 12214 | | 1.22 | 0.616 6 | **2.29** | 0.045 2 |
| N/A | N/A | X - 12216 | | 0.69 | 0.527 7 | 2.02 | 0.253 8 |
| N/A | N/A | X - 12230 | | 0.94 | 0.454 2 | 1.1 | 0.295 1 |
| N/A | N/A | X - 12231 - retired for sulfate of piperine metabolite C16H19NO3 (3)* | | 0.74 | 0.217 9 | 0.95 | 0.847 5 |
| N/A | N/A | X - 12261 | | 0.8 | 0.406 4 | **1.76** | 0.040 6 |
| N/A | N/A | X - 12263 | | 0.56 | 0.116 9 | **2.24** | 0.040 6 |
| N/A | N/A | X - 12398 | | 0.69 | 0.502 3 | 1.26 | 0.682 9 |

| | | | | | | | |
|---|---|---|---|---|---|---|---|
| N/A | N/A | X - 12410 | | 0.74 | 0.6510 | 1.95 | 0.3403 |
| N/A | N/A | X - 12411 | | 1.05 | 0.9014 | 1.24 | 0.6289 |
| N/A | N/A | X - 12442 | | 0.75 | 0.2680 | 0.94 | 0.8170 |
| N/A | N/A | X - 12456 | | 1.05 | 0.8815 | 0.66 | 0.1976 |
| N/A | N/A | X - 12462 | | 1.05 | 0.8960 | 0.55 | 0.1346 |
| N/A | N/A | X - 12472 | | 0.66 | 0.4420 | 0.86 | 0.7882 |
| N/A | N/A | X - 12511 - retired for N-acetyl-2-aminoctanoate | | 1.04 | 0.8763 | 0.85 | 0.5667 |
| N/A | N/A | X - 12565 | | 2.28 | 0.1893 | 1.12 | 0.8641 |
| N/A | N/A | X - 12680 | | **0.46** | 0.0305 | 1.01 | 0.9751 |
| N/A | N/A | X - 12688 - retired for N-acetyl-isoputrenine* | | 0.73 | 0.2086 | 0.84 | 0.5009 |
| N/A | N/A | X - 12695 - retired for alpha-keto-glutaramate** | | 0.61 | 0.1805 | 1.09 | 0.8250 |
| N/A | N/A | X - 12721 | | 0.98 | 0.9082 | 1.12 | 0.5003 |
| N/A | N/A | X - 12733 | | 0.68 | 0.2983 | 1.38 | 0.4012 |
| N/A | N/A | X - 12739 | | 0.6 | 0.1513 | 0.69 | 0.3132 |
| N/A | N/A | X - 12740 | | 1.34 | 0.5365 | 0.71 | 0.4818 |
| N/A | N/A | X - 12753 | | **0.69** | 0.0596 | **0.64** | 0.0360 |
| N/A | N/A | X - 12813 | | 1.39 | 0.5568 | 0.98 | 0.9774 |
| N/A | N/A | X - 12814 - retired for glucuronide of C12H22O4 (1) | | 0.99 | 0.9392 | 1.22 | 0.3282 |
| N/A | N/A | X - 12815 | | 1.06 | 0.7569 | 1.06 | 0.7581 |
| N/A | N/A | X - 12818 | | 0.79 | 0.5143 | 0.76 | 0.4510 |
| N/A | N/A | X - 12821 | | 0.88 | 0.8380 | 1.61 | 0.4461 |
| N/A | N/A | X - 12822 | | 0.64 | 0.4222 | 1.17 | 0.7952 |
| N/A | N/A | X - 12828 | | 1.37 | 0.3924 | **2.43** | 0.0227 |
| N/A | N/A | X - 12831 - retired for glucuronide of C14H26O4 (1)* | | 0.93 | 0.5892 | 0.98 | 0.8804 |
| N/A | N/A | X - 12839 | | 0.89 | 0.6468 | 0.82 | 0.4835 |
| N/A | N/A | X - 12844 | | **0.62** | 0.0171 | 0.98 | 0.9299 |
| N/A | N/A | X - 12849 | | 0.79 | 0.3067 | 1.15 | 0.5433 |
| N/A | N/A | X - 12879 | | 0.97 | 0.9579 | 1.66 | 0.4320 |
| N/A | N/A | X - 12906 | | 1.42 | 0.4200 | 1.59 | 0.3118 |
| N/A | N/A | X - 13007 | | 1.23 | 0.4304 | **1.62** | 0.0789 |
| N/A | N/A | X - 13431 | | 1.06 | 0.9019 | **2.66** | 0.0615 |
| N/A | N/A | X - 13507 | | 0.63 | 0.1269 | 1.36 | 0.3219 |
| N/A | N/A | X - 13553 | | 0.94 | 0.8884 | 2.04 | 0.1538 |
| N/A | N/A | X - 13723 | | 0.76 | 0.3371 | **2.13** | 0.0140 |

| | | | | | | | |
|---|---|---|---|---|---|---|---|
| N/A | N/A | X - 13729 | | 0.79 | 0.5451 | 0.79 | 0.5597 |
| N/A | N/A | X - 13737 | | 0.78 | 0.2476 | 1 | 0.9889 |
| N/A | N/A | X - 13834 - retired for nonenedioate (C9:1-DC)* | | 1.03 | 0.8903 | **0.5** | 0.0009 |
| N/A | N/A | X - 13835 | | 1.04 | 0.9119 | 1.42 | 0.3639 |
| N/A | N/A | X - 13844 | | **0.51** | 0.0781 | 1.53 | 0.2791 |
| N/A | N/A | X - 14056 | | 0.51 | 0.1318 | 1.91 | 0.1594 |
| N/A | N/A | X - 14095 - retired for pyr-ala* | | 0.8 | 0.5220 | 0.89 | 0.7585 |
| N/A | N/A | X - 14096 - retired for pyr-thr* | | 0.56 | 0.1274 | 1.07 | 0.8567 |
| N/A | N/A | X - 14099 - retired for pyr-his* | | 0.88 | 0.7561 | 1.04 | 0.9278 |
| N/A | N/A | X - 14113 - retired for pyr-arg* | | 0.93 | 0.8453 | 1.63 | 0.2256 |
| N/A | N/A | X - 14141 - retired for alpha-glu-leu* | | 1.53 | 0.2085 | 0.72 | 0.3617 |
| N/A | N/A | X - 14196 - retired for pyr-tyr* | | 0.78 | 0.4523 | 1.09 | 0.7947 |
| N/A | N/A | X - 14224 | | 0.9 | 0.4552 | 1.17 | 0.2807 |
| N/A | N/A | X - 14254 | | 0.94 | 0.9141 | 1.08 | 0.8985 |
| N/A | N/A | X - 14263 | | 0.85 | 0.6264 | 0.87 | 0.6919 |
| N/A | N/A | X - 14264 | | 1.14 | 0.8075 | 0.86 | 0.7911 |
| N/A | N/A | X - 14302 - retired for pyr-ile* | | 0.77 | 0.4565 | 1.14 | 0.7166 |
| N/A | N/A | X - 14314 - retired for pyr-leu* | | 0.76 | 0.2651 | 1.17 | 0.5500 |
| N/A | N/A | X - 14320 | | 1.13 | 0.6199 | 0.84 | 0.5055 |
| N/A | N/A | X - 14337 | | 0.85 | 0.7686 | 1.41 | 0.5522 |
| N/A | N/A | X - 14364 - retired for pyr-phe* | | 0.63 | 0.2294 | 0.53 | 0.1220 |
| N/A | N/A | X - 14383 | | 1.54 | 0.3751 | 1.2 | 0.7190 |
| N/A | N/A | X - 14392 | | 1.29 | 0.5764 | 1.67 | 0.2864 |
| N/A | N/A | X - 14416 | | 1.08 | 0.8799 | 1.39 | 0.5283 |
| N/A | N/A | X - 14454 | | 1.95 | 0.2201 | 1.02 | 0.9691 |
| N/A | N/A | X - 14473 | | 1.01 | 0.9798 | 0.72 | 0.1918 |
| N/A | N/A | X - 14502 | | 0.73 | 0.6550 | 2.2 | 0.2797 |
| N/A | N/A | X - 14538 | | 0.83 | 0.6578 | 0.95 | 0.9031 |
| N/A | N/A | X - 14658 | | **0.29** | 0.0300 | **0.36** | 0.0810 |
| N/A | N/A | X - 14662 | | **0.61** | 0.0087 | 0.93 | 0.7175 |
| N/A | N/A | X - 14697 - retired for alpha-glu-ile* | | 1.09 | 0.8011 | 0.81 | 0.5619 |
| N/A | N/A | X - 14838 | | 0.67 | 0.3861 | 1.26 | 0.6277 |
| N/A | N/A | X - 14900 | | 1.94 | 0.1609 | 0.85 | 0.7327 |
| N/A | N/A | X - 14904 | | 1.33 | 0.2947 | 1.55 | 0.1208 |

| | | | | | | | |
|---|---|---|---|---|---|---|---|
| N/A | N/A | X - 15136 | | 0.79 | 0.4155 | 0.92 | 0.7730 |
| N/A | N/A | X - 15150 | | 1.35 | 0.3021 | **0.6** | 0.0921 |
| N/A | N/A | X - 15245 | | 0.77 | 0.5350 | 1.61 | 0.2739 |
| N/A | N/A | X - 15461 | | 1.04 | 0.9165 | 0.75 | 0.4493 |
| N/A | N/A | X - 15486 | | 0.69 | 0.3735 | 0.9 | 0.8057 |
| N/A | N/A | X - 15492 | | **0.66** | 0.0250 | 0.95 | 0.8006 |
| N/A | N/A | X - 15497 | | 0.4 | 0.1049 | 1.24 | 0.7098 |
| N/A | N/A | X - 15608 | | 1.21 | 0.5474 | 1 | 0.9994 |
| N/A | N/A | X - 15646 - retired for gamma-CEHC sulfate* | | 0.97 | 0.9594 | 1.69 | 0.3422 |
| N/A | N/A | X - 15666 | | 0.85 | 0.4616 | 0.84 | 0.4686 |
| N/A | N/A | X - 15843 | | 0.83 | 0.5957 | 0.82 | 0.5761 |
| N/A | N/A | X - 15853 | | 0.93 | 0.8746 | 1.14 | 0.7829 |
| N/A | N/A | X - 15854 | | 1.29 | 0.6308 | 1.29 | 0.6463 |
| N/A | N/A | X - 16060 | | 0.95 | 0.8672 | 1.13 | 0.7222 |
| N/A | N/A | X - 16071 | | 0.56 | 0.1128 | 0.96 | 0.9101 |
| N/A | N/A | X - 16343 | | 0.96 | 0.9075 | 0.77 | 0.5163 |
| N/A | N/A | X - 16391 | | 1.09 | 0.8337 | 1.35 | 0.4679 |
| N/A | N/A | X - 16580 | | 0.72 | 0.3445 | 0.57 | 0.1210 |
| N/A | N/A | X - 16654 | | 0.39 | 0.1331 | 0.61 | 0.4489 |
| N/A | N/A | X - 16946 | | 0.87 | 0.6199 | 0.81 | 0.4640 |
| N/A | N/A | X - 16947 - retired for glucuronide of C10H18O2 (7)* | | 0.96 | 0.6733 | 0.98 | 0.8544 |
| N/A | N/A | X - 17009 | | 1.13 | 0.7736 | 1.14 | 0.7739 |
| N/A | N/A | X - 17010 | | 1.19 | 0.4967 | 1.17 | 0.5454 |
| N/A | N/A | X - 17162 | | 0.94 | 0.8931 | 0.72 | 0.5169 |
| N/A | N/A | X - 17299 | | 0.56 | 0.4040 | 0.95 | 0.9387 |
| N/A | N/A | X - 17325 | | 0.79 | 0.5045 | 1.09 | 0.8125 |
| N/A | N/A | X - 17327 | | 0.83 | 0.6099 | 0.96 | 0.9108 |
| N/A | N/A | X - 17335 | | 0.75 | 0.2720 | **0.62** | 0.0866 |
| N/A | N/A | X - 17348 | | **2.34** | 0.0483 | 1.49 | 0.3815 |
| N/A | N/A | X - 17349 | | 0.63 | 0.4161 | 0.9 | 0.8596 |
| N/A | N/A | X - 17353 | | 1.05 | 0.8643 | 1.09 | 0.7536 |
| N/A | N/A | X - 17359 | | **0.65** | 0.0004 | 0.91 | 0.4531 |
| N/A | N/A | X - 17438 | | 0.9 | 0.7595 | **2.21** | 0.0213 |
| N/A | N/A | X - 17469 | | 1.12 | 0.7864 | 1.24 | 0.6099 |

| | | | | | | | |
|---|---|---|---|---|---|---|---|
| N/A | N/A | X - 17653 | | 0.85 | 0.654<br>1 | 0.94 | 0.873<br>9 |
| N/A | N/A | X - 17655 | | 1 | 1.000<br>0 | **0.63** | 0.047<br>7 |
| N/A | N/A | X - 17674 | | 0.84 | 0.700<br>2 | 1.16 | 0.745<br>5 |
| N/A | N/A | X - 17686 | | 0.81 | 0.432<br>4 | 1.27 | 0.384<br>0 |
| N/A | N/A | X - 17705 - retired for N-acetylglucosamine conjugate of C24H40O4 bile acid** | | 0.63 | 0.221<br>1 | 0.99 | 0.969<br>1 |
| N/A | N/A | X - 17709 | | 0.61 | 0.191<br>4 | 1.88 | 0.106<br>1 |
| N/A | N/A | X - 17749 | | 0.8 | 0.528<br>3 | 1.17 | 0.666<br>0 |
| N/A | N/A | X - 17808 | | 0.74 | 0.474<br>1 | **2.11** | 0.098<br>3 |
| N/A | N/A | X - 17838 | | 2.02 | 0.157<br>7 | 1.56 | 0.392<br>5 |
| N/A | N/A | X - 17842 | | 0.9 | 0.775<br>0 | 1.33 | 0.472<br>6 |
| N/A | N/A | X - 17852 | | 0.89 | 0.824<br>4 | 1.99 | 0.198<br>7 |
| N/A | N/A | X - 17869 | | 0.76 | 0.484<br>2 | 1.09 | 0.842<br>8 |
| N/A | N/A | X - 17876 | | 1.07 | 0.866<br>3 | 1.58 | 0.259<br>8 |
| N/A | N/A | X - 17877 | | 0.95 | 0.906<br>7 | 1.68 | 0.291<br>5 |
| N/A | N/A | X - 17895 | | 0.77 | 0.459<br>6 | 1.74 | 0.130<br>7 |
| N/A | N/A | X - 17910 | | 0.96 | 0.915<br>0 | 1.21 | 0.661<br>7 |
| N/A | N/A | X - 17919 | | 0.86 | 0.744<br>0 | 1.27 | 0.613<br>5 |
| N/A | N/A | X - 17960 | | 1.53 | 0.370<br>4 | 1.93 | 0.186<br>1 |
| N/A | N/A | X - 17969 | | 1.13 | 0.800<br>4 | 1.67 | 0.303<br>7 |
| N/A | N/A | X - 17978 | | 1.21 | 0.719<br>5 | 1.08 | 0.884<br>0 |
| N/A | N/A | X - 17983 | | 1.18 | 0.730<br>8 | 1.03 | 0.955<br>7 |
| N/A | N/A | X - 17984 | | 1.23 | 0.536<br>9 | 1.06 | 0.865<br>1 |
| N/A | N/A | X - 18059 | | 0.95 | 0.789<br>1 | 1.22 | 0.328<br>7 |
| N/A | N/A | X - 18162 | | 1.2 | 0.632<br>5 | 0.85 | 0.689<br>6 |
| N/A | N/A | X - 18165 | | 0.88 | 0.759<br>0 | 0.81 | 0.628<br>5 |
| N/A | N/A | X - 18167 | | 0.81 | 0.711<br>3 | 1.16 | 0.800<br>5 |
| N/A | N/A | X - 18278 | | 1.72 | 0.332<br>7 | 1.38 | 0.580<br>7 |
| N/A | N/A | X - 18345 | | 0.95 | 0.911<br>9 | 1.26 | 0.621<br>7 |
| N/A | N/A | X - 18349 | | 0.92 | 0.773<br>6 | 0.82 | 0.514<br>4 |
| N/A | N/A | X - 18410 | | 0.5 | 0.189<br>6 | 0.83 | 0.730<br>7 |
| N/A | N/A | X - 18888 | | 0.73 | 0.426<br>9 | 0.69 | 0.382<br>3 |
| N/A | N/A | X - 18889 | | 0.44 | 0.134<br>9 | 1.22 | 0.732<br>2 |
| N/A | N/A | X - 18935 | | 0.9 | 0.776<br>9 | 1.22 | 0.608<br>9 |
| N/A | N/A | X - 18938 | | 1.04 | 0.869 | 1.44 | 0.117 |

| | | | | | | | |
|---|---|---|---|---|---|---|---|
| | | | | | 6 | | 7 |
| N/A | N/A | X - 19220 | | 0.55 | 0.126 4 | 0.95 | 0.898 9 |
| N/A | N/A | X - 19232 | | 0.83 | 0.488 5 | 0.98 | 0.929 5 |
| N/A | N/A | X - 19299 | | 0.85 | 0.658 7 | 0.81 | 0.568 7 |
| N/A | N/A | X - 19369 | | 0.97 | 0.937 7 | 1.64 | 0.161 4 |
| N/A | N/A | X - 19434 | | **0.43** | 0.043 8 | 0.94 | 0.891 6 |
| N/A | N/A | X - 19452 | | **0.41** | 0.014 0 | 0.7 | 0.358 7 |
| N/A | N/A | X - 19561 | | 0.58 | 0.225 5 | 1.86 | 0.187 2 |
| N/A | N/A | X - 19746 | | 0.56 | 0.227 5 | 1.32 | 0.584 9 |
| N/A | N/A | X - 19751 | | 0.58 | 0.231 0 | 1.16 | 0.753 6 |
| N/A | N/A | X - 19763 | | 0.83 | 0.645 0 | 1.31 | 0.525 1 |
| N/A | N/A | X - 19914 | | 1 | 1.000 0 | 1.04 | 0.892 1 |
| N/A | N/A | X - 19917 | | 1 | | 1 | |
| N/A | N/A | X - 19920 | | 1.28 | 0.102 2 | 1.11 | 0.515 4 |
| N/A | N/A | X - 19921 | | 0.64 | 0.198 7 | 1.49 | 0.264 9 |
| N/A | N/A | X - 19923 | | 0.85 | 0.386 5 | 1.02 | 0.914 8 |
| N/A | N/A | X - 19924 | | 1.18 | 0.533 3 | 1.09 | 0.754 0 |
| N/A | N/A | X - 19928 | | 0.69 | 0.437 7 | 1.43 | 0.480 1 |
| N/A | N/A | X - 19929 | | 0.71 | 0.255 1 | 1.29 | 0.423 5 |
| N/A | N/A | X - 19931 | | 0.79 | 0.584 3 | 1.22 | 0.671 0 |
| N/A | N/A | X - 19932 - retired for pyr-trp* | | 0.71 | 0.374 5 | 0.97 | 0.934 7 |
| N/A | N/A | X - 19934 | | 1.06 | 0.758 0 | 1.07 | 0.727 9 |
| N/A | N/A | X - 19940 | | 0.71 | 0.345 3 | 1.06 | 0.876 8 |
| N/A | N/A | X - 20100 | | 1.23 | 0.647 8 | 1.44 | 0.434 5 |
| N/A | N/A | X - 20172 | | 1.28 | 0.648 5 | 1.23 | 0.707 9 |
| N/A | N/A | X - 20185 | | 1.11 | 0.812 8 | 1.29 | 0.568 0 |
| N/A | N/A | X - 20197 | | 1.33 | 0.568 3 | 1.4 | 0.514 6 |
| N/A | N/A | X - 20624 | | 0.93 | 0.875 6 | 1.29 | 0.596 4 |
| N/A | N/A | X - 20747 | | 1.62 | 0.341 0 | 1.02 | 0.965 5 |
| N/A | N/A | X - 20756 | | 0.56 | 0.126 1 | 1.26 | 0.555 9 |
| N/A | N/A | X - 20768 | | 0.71 | 0.224 3 | 1.05 | 0.861 9 |
| N/A | N/A | X - 21283 | | 0.95 | 0.888 2 | 1.29 | 0.542 1 |
| N/A | N/A | X - 21285 | | **0.55** | 0.011 5 | 0.69 | 0.122 1 |
| N/A | N/A | X - 21295 | | 0.52 | 0.268 9 | **3.02** | 0.076 0 |

| | | | | | | | |
|---|---|---|---|---|---|---|---|
| N/A | N/A | X - 21319 | | 1.36 | 0.3090 | 1.36 | 0.3409 |
| N/A | N/A | X - 21327 | | 1.02 | 0.9554 | **5.52** | 0.0000 |
| N/A | N/A | X - 21343 - retired for dodecadienoate (12:2)* | | 1.09 | 0.8419 | 0.8 | 0.6187 |
| N/A | N/A | X - 21353 | | 0.77 | 0.2892 | 0.98 | 0.9277 |
| N/A | N/A | X - 21410 | | 0.92 | 0.8287 | 0.69 | 0.3318 |
| N/A | N/A | X - 21441 | | 0.81 | 0.1065 | 0.99 | 0.9542 |
| N/A | N/A | X - 21448 | | 0.85 | 0.5731 | 0.89 | 0.6964 |
| N/A | N/A | X - 21467 | | 0.63 | 0.3074 | 1.2 | 0.7062 |
| N/A | N/A | X - 21470 | | 0.85 | 0.6916 | **0.49** | 0.0990 |
| N/A | N/A | X - 21471 | | 0.83 | 0.6066 | 1.01 | 0.9858 |
| N/A | N/A | X - 21668 - retired for glyco-beta-muricholate** | | **0.54** | 0.0692 | 1.03 | 0.9344 |
| N/A | N/A | X - 21729 | | 1.1 | 0.8513 | 1.61 | 0.3661 |
| N/A | N/A | X - 21785 | | 0.68 | 0.5688 | 1.03 | 0.9621 |
| N/A | N/A | X - 21787 | | 1.15 | 0.6711 | 1.44 | 0.2678 |
| N/A | N/A | X - 21788 | | 0.91 | 0.8724 | 0.82 | 0.7383 |
| N/A | N/A | X - 21796 | | 0.97 | 0.8393 | 0.89 | 0.5157 |
| N/A | N/A | X - 21821 | | 0.75 | 0.4065 | 1.04 | 0.9166 |
| N/A | N/A | X - 21826 - retired for glutamine conjugate of C8H12O2 (2)* | | 0.69 | 0.3117 | 0.82 | 0.6094 |
| N/A | N/A | X - 21851 | | 0.82 | 0.1452 | 0.93 | 0.6358 |
| N/A | N/A | X - 21959 | | 1.18 | 0.5593 | 1.62 | 0.1005 |
| N/A | N/A | X - 21963 | | 1.03 | 0.8922 | 1 | 0.9925 |
| N/A | N/A | X - 22030 | | 1.75 | 0.1365 | 1.67 | 0.1944 |
| N/A | N/A | X - 22035 - retired for pyr-pro* | | **0.4** | 0.0221 | 1.01 | 0.9736 |
| N/A | N/A | X - 22062 | | 1.27 | 0.4404 | 1.21 | 0.5516 |
| N/A | N/A | X - 22085 | | 0.65 | 0.4472 | 0.92 | 0.8946 |
| N/A | N/A | X - 22099 | | **0.63** | 0.0942 | 0.89 | 0.6743 |
| N/A | N/A | X - 22102 | | 0.46 | 0.1974 | 0.38 | 0.1366 |
| N/A | N/A | X - 22142 | | 0.51 | 0.1690 | 0.78 | 0.6254 |
| N/A | N/A | X - 22162 | | 0.7 | 0.3323 | **0.42** | 0.0261 |
| N/A | N/A | X - 22520 | | 1.05 | 0.8670 | 0.9 | 0.7258 |
| N/A | N/A | X - 22800 | | 0.94 | 0.8700 | 1.21 | 0.6232 |
| N/A | N/A | X - 22810 | | 0.91 | 0.7234 | 0.85 | 0.5296 |
| N/A | N/A | X - 23159 | | 0.67 | 0.1431 | 0.89 | 0.6844 |
| N/A | N/A | X - 23160 | | 0.86 | 0.668 | 0.97 | 0.929 |

| | | | | | | | |
|---|---|---|---|---|---|---|---|
| | | | | | 6 | | 2 |
| N/A | N/A | X - 23163 - retired for gamma-CEHC taurine* | | 0.65 | 0.1645 | 1.22 | 0.5343 |
| N/A | N/A | X - 23164 - retired for alpha-CEHC taurine* | | 0.68 | 0.1656 | 0.99 | 0.9670 |
| N/A | N/A | X - 23173 | | 0.83 | 0.6689 | 1.23 | 0.6607 |
| N/A | N/A | X - 23188 | | 1.38 | 0.2805 | 1.58 | 0.1403 |
| N/A | N/A | X - 23191 | | 0.89 | 0.7751 | **3.99** | 0.0012 |
| N/A | N/A | X - 23193 | | 0.81 | 0.1833 | **1.39** | 0.0413 |
| N/A | N/A | X - 23196 | | 1.23 | 0.4223 | **2.07** | 0.0087 |
| N/A | N/A | X - 23201 | | 0.96 | 0.9190 | 0.78 | 0.5465 |
| N/A | N/A | X - 23223 | | 1.02 | 0.9727 | 1.84 | 0.2594 |
| N/A | N/A | X - 23228 | | 0.93 | 0.7286 | 1.03 | 0.8801 |
| N/A | N/A | X - 23236 | | 0.91 | 0.7356 | 1.5 | 0.1373 |
| N/A | N/A | X - 23238 | | 0.68 | 0.1609 | 0.77 | 0.3572 |
| N/A | N/A | X - 23259 | | 1 | 1.0000 | **1.49** | 0.0180 |
| N/A | N/A | X - 23267 | | 0.72 | 0.4557 | **4.42** | 0.0015 |
| N/A | N/A | X - 23277 | | 0.9 | 0.6752 | **2.15** | 0.0035 |
| N/A | N/A | X - 23287 | | 0.92 | 0.8007 | 1.11 | 0.7364 |
| N/A | N/A | X - 23314 | | 1.03 | 0.9384 | 0.64 | 0.2075 |
| N/A | N/A | X - 23323 | | 1.14 | 0.7629 | **2.22** | 0.0762 |
| N/A | N/A | X - 23324 | | 1.17 | 0.6762 | 1.59 | 0.2340 |
| N/A | N/A | X - 23360 | | 0.99 | 0.9587 | **1.28** | 0.0719 |
| N/A | N/A | X - 23369 | | 0.56 | 0.1425 | 0.8 | 0.5913 |
| N/A | N/A | X - 23423 | | 0.91 | 0.7456 | 0.93 | 0.8110 |
| N/A | N/A | X - 23424 | | 0.87 | 0.5441 | 0.88 | 0.5820 |
| N/A | N/A | X - 23429 | | 0.93 | 0.8538 | 1.06 | 0.8865 |
| N/A | N/A | X - 23432 | | **2.74** | 0.0475 | 0.89 | 0.8275 |
| N/A | N/A | X - 23438 | | 0.83 | 0.3770 | 0.97 | 0.8813 |
| N/A | N/A | X - 23451 | | 0.49 | 0.2067 | **2.67** | 0.0905 |
| N/A | N/A | X - 23453 | | **3.25** | 0.0096 | 2.02 | 0.1326 |
| N/A | N/A | X - 23456 | | 1.08 | 0.8781 | 1.38 | 0.5567 |
| N/A | N/A | X - 23469 | | 1.05 | 0.9137 | 1.58 | 0.3414 |
| N/A | N/A | X - 23470 | | 1.2 | 0.6473 | **2.34** | 0.0379 |
| N/A | N/A | X - 23481 | | 0.88 | 0.4814 | 1.08 | 0.6610 |
| N/A | N/A | X - 23482 | | 0.86 | 0.6274 | **1.83** | 0.0648 |

| | | | | | | | |
|---|---|---|---|---|---|---|---|
| N/A | N/A | X - 23507 | | 0.93 | 0.7490 | 1.18 | 0.4914 |
| N/A | N/A | X - 23557 | | 0.8 | 0.6430 | 1.32 | 0.5861 |
| N/A | N/A | X - 23581 | | 0.55 | 0.2937 | 2.19 | 0.1860 |
| N/A | N/A | X - 23585 | | 1.03 | 0.9517 | 0.9 | 0.8414 |
| N/A | N/A | X - 23587 | | 0.79 | 0.6581 | 0.9 | 0.8470 |
| N/A | N/A | X - 23600 | | 1.15 | 0.7536 | 1 | 0.9935 |
| N/A | N/A | X - 23610 | | 0.91 | 0.7263 | 0.84 | 0.5424 |
| N/A | N/A | X - 23617 | | 0.68 | 0.1647 | 1.24 | 0.4538 |
| N/A | N/A | X - 23623 | | 0.82 | 0.3763 | 1.14 | 0.5769 |
| N/A | N/A | X - 23626 | | **0.57** | 0.0924 | 1.18 | 0.6375 |
| N/A | N/A | X - 23637 | | 0.84 | 0.5972 | 1.25 | 0.5066 |
| N/A | N/A | X - 23639 | | 1.13 | 0.7483 | 1.24 | 0.5983 |
| N/A | N/A | X - 23644 | | 0.5 | 0.1664 | 0.97 | 0.9559 |
| N/A | N/A | X - 23652 | | 0.89 | 0.6565 | 1.18 | 0.5473 |
| N/A | N/A | X - 23655 | | 1.31 | 0.3958 | 1.05 | 0.8894 |
| N/A | N/A | X - 23662 | | 0.74 | 0.5629 | 0.76 | 0.6038 |
| N/A | N/A | X - 23665 | | 0.98 | 0.9606 | 1.05 | 0.8940 |
| N/A | N/A | X - 23680 | | 1.13 | 0.7230 | 0.86 | 0.6786 |
| N/A | N/A | X - 23710 | | 0.91 | 0.8216 | 0.93 | 0.8550 |
| N/A | N/A | X - 23728 | | 1 | 0.9993 | **2.68** | 0.0266 |
| N/A | N/A | X - 23729 | | 0.87 | 0.6462 | 1.06 | 0.8647 |
| N/A | N/A | X - 23732 | | 1.25 | 0.6047 | **2.25** | 0.0763 |
| N/A | N/A | X - 23734 | | 0.89 | 0.7210 | 1 | 0.9882 |
| N/A | N/A | X - 23736 | | 0.75 | 0.3788 | 0.93 | 0.8400 |
| N/A | N/A | X - 23737 | | 0.9 | 0.7393 | 1.46 | 0.2490 |
| N/A | N/A | X - 23739 | | 0.72 | 0.5322 | 1.7 | 0.3283 |
| N/A | N/A | X - 23747 - retired for diacetylspermidine* | | 0.89 | 0.6873 | 0.97 | 0.9237 |
| N/A | N/A | X - 23753 | | 1.05 | 0.9008 | **2.64** | 0.0179 |
| N/A | N/A | X - 23756 - retired for octadecenedioylcarnitne (C18:1-DC)* | | 1.62 | 0.2369 | 1.21 | 0.6408 |
| N/A | N/A | X - 23764 | | 1.21 | 0.5788 | 1.32 | 0.4344 |
| N/A | N/A | X - 23767 | | 0.6 | 0.3481 | 0.87 | 0.8039 |
| N/A | N/A | X - 23776 | | 0.57 | 0.3648 | 0.93 | 0.9054 |
| N/A | N/A | X - 23782 | | 0.72 | 0.3158 | 1.03 | 0.9251 |
| N/A | N/A | X - 23908 | | 0.97 | 0.952 | **0.3** | 0.015 |

| | | | | | | | |
|---|---|---|---|---|---|---|---|
| | | | | | 2 | | 4 |
| N/A | N/A | X - 23919 | | 1.06 | 0.8081 | 1.07 | 0.8022 |
| N/A | N/A | X - 23925 | | 1.54 | 0.2258 | 0.82 | 0.5811 |
| N/A | N/A | X - 23939 | | 0.94 | 0.1063 | 1 | 1.0000 |
| N/A | N/A | X - 23942 | | 0.73 | 0.2677 | 1.03 | 0.9154 |
| N/A | N/A | X - 23974 | | 0.94 | 0.8026 | 0.85 | 0.5430 |
| N/A | N/A | X - 24027 | | 0.88 | 0.3017 | 1.04 | 0.7242 |
| N/A | N/A | X - 24077 | | 0.46 | 0.1560 | **3.62** | 0.0238 |
| N/A | N/A | X - 24084 | | 0.91 | 0.8209 | 0.95 | 0.9042 |
| N/A | N/A | X - 24089 | | **0.61** | 0.0605 | 1.1 | 0.7371 |
| N/A | N/A | X - 24137 | | 1.05 | 0.8681 | 1.06 | 0.8404 |
| N/A | N/A | X - 24143 | | 1.06 | 0.8488 | 1.01 | 0.9828 |
| N/A | N/A | X - 24177 | | 0.75 | 0.6176 | 1.28 | 0.6854 |
| N/A | N/A | X - 24207 - retired for pheophytin A | | 0.97 | 0.9534 | 1.41 | 0.5815 |
| N/A | N/A | X - 24208 | | **0.51** | 0.0733 | 1.32 | 0.4702 |
| N/A | N/A | X - 24216 | | **0.37** | 0.0284 | **0.39** | 0.0451 |
| N/A | N/A | X - 24220 | | 0.73 | 0.1973 | 0.9 | 0.6887 |
| N/A | N/A | X - 24231 | | 0.95 | 0.9085 | 1.57 | 0.3712 |
| N/A | N/A | X - 24246 | | 0.61 | 0.3182 | 0.7 | 0.4912 |
| N/A | N/A | X - 24272 | | 0.99 | 0.9228 | 1.22 | 0.1038 |
| N/A | N/A | X - 24329 | | 0.66 | 0.1723 | 1.09 | 0.7859 |
| N/A | N/A | X - 24348 | | **0.34** | 0.0108 | 1.38 | 0.4518 |
| N/A | N/A | X - 24359 | | 0.73 | 0.6364 | 1.12 | 0.8657 |
| N/A | N/A | X - 24400 | | 0.8 | 0.5223 | 0.96 | 0.9100 |
| N/A | N/A | X - 24404 - retired for umbelliferone | | 0.66 | 0.2520 | 0.92 | 0.8226 |
| N/A | N/A | X - 24408 | | 0.81 | 0.5387 | 1.3 | 0.4659 |
| N/A | N/A | X - 24410 | | 1.32 | 0.5951 | 1.19 | 0.7505 |
| N/A | N/A | X - 24412 | | 0.89 | 0.6197 | 0.71 | 0.1571 |
| N/A | N/A | X - 24413 | | 0.74 | 0.6373 | 0.91 | 0.8949 |
| N/A | N/A | X - 24425 | | 0.88 | 0.7417 | 1.73 | 0.1717 |
| N/A | N/A | X - 24431 | | 0.95 | 0.9132 | 1.25 | 0.6272 |
| N/A | N/A | X - 24452 | | **0.41** | 0.0100 | 1.1 | 0.7874 |
| N/A | N/A | X - 24455 | | 0.85 | 0.3791 | 1.16 | 0.4595 |
| N/A | N/A | X - 24456 | | 0.86 | 0.4861 | 1.16 | 0.5218 |

| | | | | | | | |
|---|---|---|---|---|---|---|---|
| N/A | N/A | X - 24465 | | **0.55** | 0.0577 | 0.92 | 0.7922 |
| N/A | N/A | X - 24474 | | 1.03 | 0.9118 | 1.29 | 0.3961 |
| N/A | N/A | X - 24542 | | **0.23** | 0.0048 | 1.89 | 0.2344 |
| N/A | N/A | X - 24544 | | 0.82 | 0.6920 | 0.89 | 0.8216 |
| N/A | N/A | X - 24545 | | 0.57 | 0.1219 | 0.56 | 0.1159 |
| N/A | N/A | X - 24546 | | 0.9 | 0.7600 | **0.54** | 0.0872 |
| N/A | N/A | X - 24550 - retired for PEG-glycoside, fragment 07* | | 0.81 | 0.4078 | 1.01 | 0.9625 |
| N/A | N/A | X - 24551 - retired for PEG-glucuronide, fragment 04* | | 0.92 | 0.7088 | 0.99 | 0.9737 |
| N/A | N/A | X - 24552 - retired for PEG-glycoside, fragment 05* | | 0.68 | 0.1605 | 1.08 | 0.7962 |
| N/A | N/A | X - 24555 - retired for PEG-glycoside, fragment 04* | | 0.83 | 0.4712 | 1.07 | 0.8145 |
| N/A | N/A | X - 24556 | | 0.83 | 0.6155 | 1.79 | 0.1161 |
| N/A | N/A | X - 24557 - retired for PEG-glucuronide, fragment 06* | | 0.99 | 0.9527 | 0.97 | 0.8730 |
| N/A | N/A | X - 24558 - retired for PEG-glycoside, fragment 06* | | 0.76 | 0.3202 | 0.97 | 0.9065 |
| N/A | N/A | X - 24559 - retired for PEG-glycoside, fragment 03* | | 0.82 | 0.3917 | 1.06 | 0.8167 |
| N/A | N/A | X - 24560 - retired for PEG-glycoside, fragment 08* | | 1.02 | 0.9186 | 1 | 0.9953 |
| N/A | N/A | X - 24568 - retired for N4-acetylsulfamethoxazole* | | 0.77 | 0.4473 | **0.29** | 0.0008 |
| N/A | N/A | X - 24569 - retired for N4-acetyl-5-hydroxysulfamethoxazole* | | 0.84 | 0.2967 | **0.64** | 0.0164 |
| N/A | N/A | X - 24574 | | 0.72 | 0.4498 | 0.54 | 0.1728 |
| N/A | N/A | X - 24586 | | **0.43** | 0.0083 | 0.83 | 0.5857 |
| N/A | N/A | X - 24588 | | **0.32** | 0.0039 | 0.94 | 0.8926 |
| N/A | N/A | X - 24590 | | 0.78 | 0.3164 | 1.19 | 0.4895 |
| N/A | N/A | X - 24608 | | 0.77 | 0.4059 | 0.6 | 0.1143 |
| N/A | N/A | X - 24609 | | **0.56** | 0.0221 | 0.85 | 0.5530 |
| N/A | N/A | X - 24626 | | 0.75 | 0.2529 | 0.89 | 0.6641 |
| N/A | N/A | X - 24630 | | 0.94 | 0.4853 | 1 | 1.0000 |
| N/A | N/A | X - 24640 | | 1.28 | 0.5952 | **2.88** | 0.0281 |
| N/A | N/A | X - 24641 | | 0.72 | 0.5406 | 2.23 | 0.1447 |
| N/A | N/A | X - 24643 | | 1 | 1.0000 | 1 | 1.0000 |
| N/A | N/A | X - 24655 - retired for genistein sulfate | | **0.33** | 0.0145 | 1.62 | 0.3039 |
| N/A | N/A | X - 24658 | | 0.88 | 0.7169 | 1.08 | 0.8304 |
| N/A | N/A | X - 24659 | | **0.39** | 0.0086 | 0.75 | 0.4493 |
| N/A | N/A | X - 24660 | | 0.46 | 0.1139 | 1.36 | 0.5446 |
| N/A | N/A | X - 24669 | | 0.45 | 0.1267 | 1.53 | 0.4300 |
| N/A | N/A | X - 24670 | | **0.41** | 0.013 | 1.06 | 0.871 |

| | | | | | | | |
|---|---|---|---|---|---|---|---|
| | | | | | 5 | | 3 |
| N/A | N/A | X - 24682 - retired for pyr-pro* | | **0.27** | 0.012 3 | 0.79 | 0.668 6 |
| N/A | N/A | X - 24683 | | 0.64 | 0.261 5 | 1.13 | 0.767 2 |
| N/A | N/A | X - 24686 | | 0.56 | 0.113 6 | 0.96 | 0.906 7 |
| N/A | N/A | X - 24693 - retired for histidine betaine (hercynine)* | | 1.05 | 0.924 5 | 1.45 | 0.465 0 |
| N/A | N/A | X - 24697 | | 1.05 | 0.897 5 | 1.24 | 0.568 1 |
| N/A | N/A | X - 24704 | | **0.45** | 0.017 8 | 0.81 | 0.554 0 |
| N/A | N/A | X - 24711 | | **0.37** | 0.002 8 | 1.2 | 0.578 9 |
| N/A | N/A | X - 24712 | | **0.42** | 0.002 9 | 0.8 | 0.473 6 |
| N/A | N/A | X - 24715 | | 0.77 | 0.362 1 | 1.03 | 0.927 9 |
| N/A | N/A | X - 24723 | | 1 | 0.980 9 | 1 | 1.000 0 |
| N/A | N/A | X - 24728 | | 0.92 | 0.866 8 | 1 | 0.996 8 |
| N/A | N/A | X - 24729 | | 1.46 | 0.111 5 | 1.17 | 0.526 9 |
| N/A | N/A | X - 24738 - retired for 3-hydroxystachydrine* | | 0.54 | 0.265 8 | 0.94 | 0.908 5 |
| N/A | N/A | X - 24753 | | 0.79 | 0.536 5 | **2.07** | 0.061 7 |
| N/A | N/A | X - 24762 | | 0.47 | 0.185 8 | 0.86 | 0.806 9 |
| N/A | N/A | X - 24766 | | **0.62** | 0.078 4 | 1.07 | 0.803 4 |
| N/A | N/A | X - 24767 | | 1.13 | 0.750 2 | **1.95** | 0.091 8 |
| N/A | N/A | X - 24809 | | 0.94 | 0.876 7 | 0.65 | 0.303 8 |
| N/A | N/A | X - 24813 | | 1.16 | 0.503 9 | 1.35 | 0.199 0 |
| N/A | N/A | X - 24829 | | 0.85 | 0.644 3 | 1.1 | 0.795 7 |
| N/A | N/A | X - 24831 | | **0.6** | 0.022 0 | **1.47** | 0.096 3 |
| N/A | N/A | X - 24832 | | 0.68 | 0.131 8 | 1.49 | 0.125 6 |
| N/A | N/A | X - 24840 | | 0.65 | 0.231 2 | 1.3 | 0.477 2 |
| N/A | N/A | X - 24853 | | 1.13 | 0.674 6 | 0.78 | 0.421 2 |
| N/A | N/A | X - 24854 | | **0.46** | 0.048 2 | 0.82 | 0.618 8 |
| N/A | N/A | X - 24855 | | 0.98 | 0.947 6 | 0.74 | 0.425 8 |
| N/A | N/A | X - 24931 | | 0.79 | 0.439 5 | 0.79 | 0.466 9 |
| N/A | N/A | X - 24932 | | 0.48 | 0.215 4 | 1.43 | 0.565 4 |
| N/A | N/A | X - 24947 | | **0.54** | 0.052 2 | **0.57** | 0.078 5 |
| N/A | N/A | X - 24948 | | 1.02 | 0.968 5 | 0.93 | 0.876 5 |
| N/A | N/A | X - 24974 | | 0.64 | 0.172 9 | **0.48** | 0.030 8 |
| N/A | N/A | X - 24977 | | 0.33 | 0.111 5 | **0.3** | 0.097 5 |
| N/A | N/A | X - 24978 | | 1.13 | 0.715 7 | 0.95 | 0.892 5 |

| | | | | | | | |
|---|---|---|---|---|---|---|---|
| N/A | N/A | X - 24983 | | 0.74 | 0.2085 | 0.86 | 0.5350 |
| N/A | N/A | X - 24989 | | 1.1 | 0.7792 | 1.29 | 0.4745 |
| N/A | N/A | X - 24991 | | 1.17 | 0.7046 | 1.12 | 0.7854 |
| N/A | N/A | X - 24996 | | **0.46** | 0.0057 | 1.1 | 0.7491 |
| N/A | N/A | X - 25005 | | 0.76 | 0.3725 | 1.22 | 0.5271 |
| N/A | N/A | X - 25007 | | 1.13 | 0.7052 | 1.51 | 0.2355 |
| N/A | N/A | X - 25009 | | 0.61 | 0.1128 | 1.32 | 0.3946 |
| N/A | N/A | X - 25010 | | 0.74 | 0.2707 | 0.92 | 0.7594 |
| N/A | N/A | X - 25035 | | **0.4** | 0.0896 | **3.71** | 0.0221 |
| N/A | N/A | X - 25053 | | **0.39** | 0.0609 | 1.48 | 0.4605 |
| N/A | N/A | X - 25076 | | 1.33 | 0.5756 | 1.72 | 0.3106 |

104
